# Striatal activity maintains a short-term action-outcome memory to guide future choice

**DOI:** 10.64898/2026.06.16.731709

**Authors:** Allison E. Girasole, Gil Mandelbaum, Livia C. Murray, Celia C. Beron, Madeline A. Albanese, Rebekah Y. Zhang, Bastijn J. G. van den Boom, Rebecca N. Alvarado, Daniel R. Hochbaum, Trevor M. Haynes, Maria Diaz Bobillo, Wengang Weng, Bernardo L. Sabatini

**Affiliations:** Howard Hughes Medical Institute, Department of Neurobiology, Harvard Medical School, Boston MA, 02115, USA; Kempner Institute for the Study of Natural and Artificial Intelligence, Harvard University, Allston, MA, 02134, USA; Society of Fellows, Harvard University, Cambridge, MA 02138, USA; Division of Endocrinology, Diabetes and Metabolism, Beth Israel Deaconess Medical Center, Harvard Medical School, Boston, MA 02215, USA

## Abstract

Adaptive behavior requires animals to use the outcomes of recent actions (action-outcomes) to guide future decisions. While the striatum is critical for decision-making, it is currently unclear how it is involved in the storage and retrieval of short-term associative memories. Here, we implemented a head-fixed memory-guided decision-making task in which mice use the outcome of a previous choice to determine whether to repeat or switch their next action. We show that dopamine fluctuations in the ventrolateral striatum are modulated by reward receipt or omission and recent outcome history, while the activity of direct and indirect pathway striatal projection neurons encodes recent action-outcome associations and predicts future switch/repeat choices. Closed-loop optogenetic activation and inhibition of direct and indirect pathway neurons during either the action-outcome association period or the delay preceding the next choice bidirectionally biased future actions away from those favored by reward history. Together, these findings suggest that striatal activity maintains a short-term action-outcome associative memory that links completed actions to future motor plans during adaptive decision-making.

## INTRODUCTION

Adaptive behavior requires animals to select actions that obtain resources while avoiding actions that lead to harm. Reinforcement learning provides a framework for understanding how experience shapes these choices: actions followed by positive outcomes become more likely to be repeated, whereas actions followed by negative outcomes become more likely to be avoided or updated (Sutton and Barto, 1998). Recurrent loops between cortex, basal ganglia (BG), and thalamus are central to reinforcement learning by supporting the selection and execution of goal-oriented actions, including the repetition of behaviors that lead to positive outcomes (Maia and Frank, 2011; Fee, 2012; Averbeck and O’Doherty, 2022). In this framework, dopamine-dependent plasticity in cortico-BG circuits is thought to gradually shape action policies over repeated experience (Shen et al., 2008; Maia and Frank, 2011; Yagishita et al., 2014; Iino et al., 2020; Qi et al., 2025). However, adaptive behavior also requires a more immediate computation: once an animal has executed an action and experienced its outcome, that information can rapidly impact the selection of the immediate next action. This updated action plan must then be maintained in the inter-action interval as a short-term motor memory.

Emerging work from rodents and non-human primates suggests that circuits within the striatum, the primary input nucleus of the BG, carry such short-timescale memory-related signals necessary for bridging previous actions to future behaviors through online activity (Hikosaka et al., 1989; Kim et al., 2013; Akhlaghpour et al., 2016; Nonomura et al., 2018; Tang et al., 2021; Wang et al., 2021; Yoshizawa et al., 2023; van Beest et al., 2024; Reinhold et al., 2025). Recent work using instructed memory-guided licking tasks have shown that cortico-basal ganglia-thalamo-cortical circuits contribute to persistent activity in frontal motor areas during the maintenance of short-term memory and to the selection of upcoming actions (Tang et al., 2021; Wang et al., 2021). Further, activity in posterior dorsomedial striatum encodes interactions between context, action, and outcome during sensory-motor learning, providing information necessary to guide future behavior (Reinhold et al., 2025). These short-timescale signals are likely shaped by direct (dSPN) and indirect (iSPN) pathway spiny projection neurons, which receive convergent cortical, thalamic, and dopaminergic input and provide output through distinct BG pathways (Gerfen and Surmeier, 2011; Hunnicutt et al., 2016; Watabe-Uchida et al., 2012). Once viewed as antagonistic for movement (Albin et al., 1989; Alexander et al., 1986; DeLong, 1990), dSPNs and iSPNs are increasingly understood as dynamically coactive populations whose activity regulates multiple dimensions of behavior (Cui et al., 2013; Isomura et al., 2013; Jin et al., 2014; Barbera et al., 2016; Markowitz et al., 2018; Parker et al., 2018). Prior studies have largely examined striatal contributions to sensory-instructed actions or gradual learning. However, it remains unresolved whether striatal activity maintains information about a recently completed action and its outcome to guide the selection of a future action when current sensory cues do not specify the correct choice.

Here, we use the term “short-term action-outcome memory” to refer to a transient neural representation that links a recently completed action with its outcome and biases a future choice. We tested the hypothesis that dSPN and iSPN activity contributes to this short-term action-outcome memory by linking recent choices and outcomes to future motor plans. These studies were targeted to the ventrolateral striatum (VLS), a region anatomically and functionally linked to orofacial behaviors (Lee et al., 2020; Bakhurin et al., 2020; Lee and Sabatini, 2021; Chen et al., 2021; Tang et al., 2021). We developed a memory-guided task in which mice selected between two lick spouts and used the outcome of the previous choice to determine whether to repeat or switch on the next trial, similar to established paradigms (Bari et al., 2019; Hattori et al., 2019; Su and Cohen, 2022; Hattori et al., 2023). This task separates the action-outcome association period of one trial from the choice on the next trial, allowing us to interrogate whether pathway-specific striatal activity maintains recently-acquired outcome information across the delay between choices. Using pathway-specific fiber photometry recordings from dSPNs, iSPNs, and dopamine release in the VLS, together with closed-loop optogenetic manipulations, we find that both striatal pathways encode recent action-outcome history, predict whether animals will switch or repeat their next choice, and causally influence future action selection. These findings support a model in which striatal SPN activity maintains and updates a short-term memory of the upcoming motor plan.

## RESULTS

### A head-fixed dynamic foraging task for memory-guided decision making

To examine whether striatal circuits shape action selection, we implemented a memory-guided decision-making task that requires animals to choose their next action based on the outcome of their previous action (**Figure 1A**). Following an enforced delay and neutral “go cue”, head-restrained, water-restricted mice indicate their choice by licking to one of two lick spouts located on either side of their mouths (i.e. they choose to “lick left” or “lick right”). The first lick after the cue is considered the “choice lick”. If the choice is correct, a drop of water is delivered that can be collected with subsequent licks. The task structure is organized into blocks in which only one spout is correct and delivers a water drop if chosen (i.e. the rewarded spout). After a block of random length from 4 to 8 rewards, the identities of the rewarded and unrewarded spouts reverse without any additional cue indicating the block transition.

**Figure 1.**
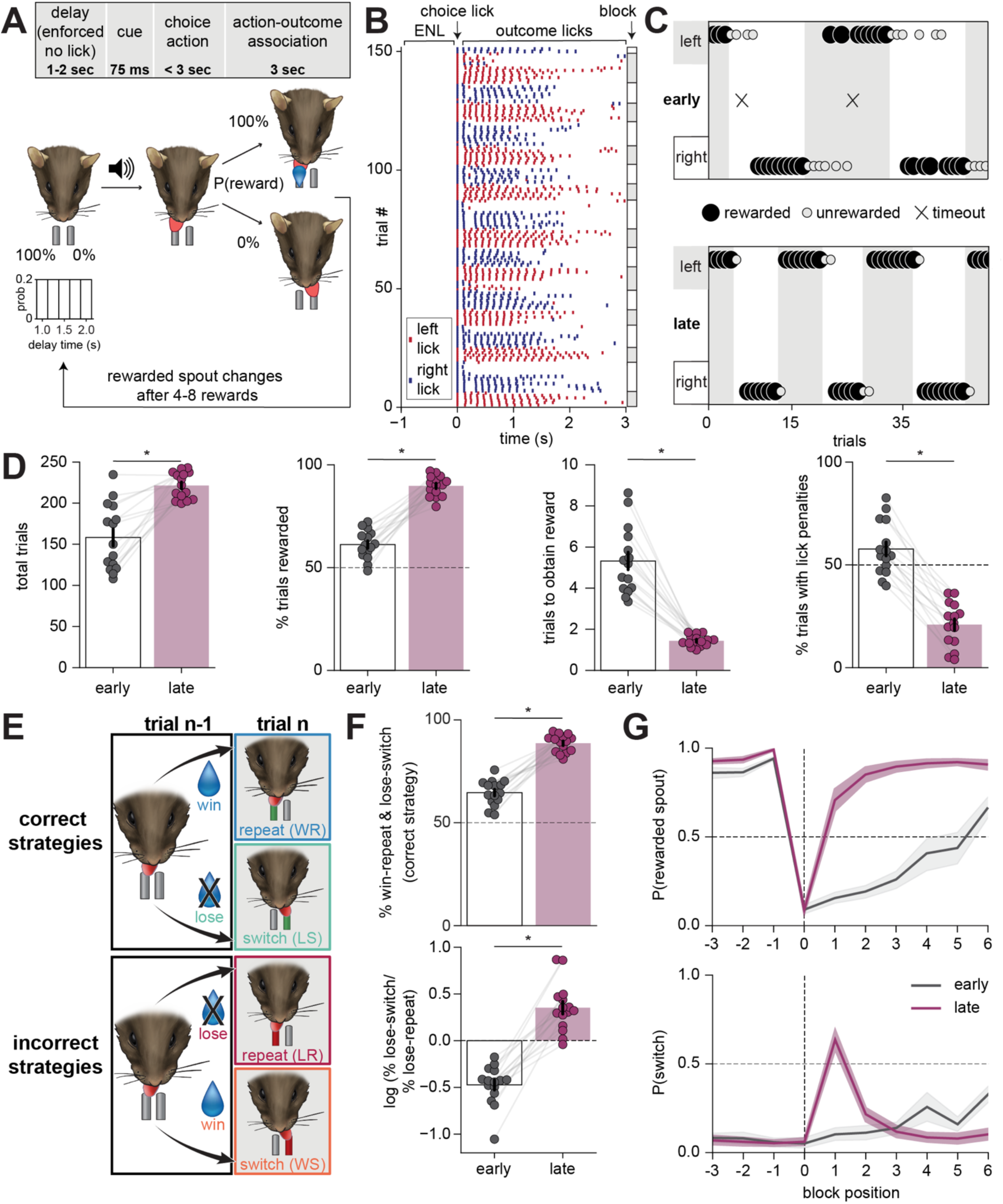
A head-fixed dynamic foraging task for memory guided decision making. **A.** *top:* Schematic showing that a trial begins with an “enforced no-lick (ENL)” delay period of variable length (1-2 seconds) that ends with an auditory cue (75 ms) after which mice have up to 3 seconds to report their choice by licking either the right or left spout. If the correct spout is chosen, a water drop is delivered. *bottom:* Graphical schematic of the trial and block structure showing the distribution of delay/ENL period durations and that the identity of the rewarded spout switches every 4-8 rewards. **B.** Raster plot from a behavioral session of a well-trained animal showing individual licks to left (red) and right (blue) spouts as a function of time relative to the choice lick (black arrow, at time=0 seconds). Block color (grey or white) indicates the identity of the rewarded spout (left or right, respectively). **C.** Representative behavior from a single session of one mouse early (1 week, *top*) and late (6 weeks, *bottom*) into training. Each dot corresponds to one trial with the vertical position indicating the chosen spout and the dot size the trial outcome (large=rewarded, small=unrewarded, no selection=timeout). Data are shown for 50 consecutive trials. **D.** Comparison of task performance early (1 week) and late (4-6 weeks) into training. *left*: Total number of trials per session. *middle/left:* Percentage of rewarded trials per session. *middle/right:* Average number of trials required to receive the first reward after a block switch. *right:* Percentage of trials that incurred lick penalties during the ENL per session. **E.** Schematic of task strategy. Mice select their next action based on their previous choice (lick left vs. right) and its outcome (reward vs. no reward). *top*: The optimal strategy to maximize rewards is to repeat the same choice after a rewarded trial (win-repeat, blue) and switch to the other spout after an unrewarded trial (lose-switch, green). *bottom:* Win-switch (orange) and lose-repeat (red) represent suboptimal, incorrect strategies. **F.** Comparison of task strategy early and late in training. *top:* Percentages of trials in which the animal used the correct strategy (win-repeat and lose-switch). *bottom:* Log of the ratio of lose-switch to lose-repeat trials. **G.** Probability of selecting the rewarded spout (*top*) and the probability of switching between spouts on the current trial relative to the last trial (*bottom*) as a function of trial position around block transitions (x=0). Solid lines represent aggregated probabilities across mice early (grey) and late (purple) in training and shading represents ± SEM across mice. **D-F.** N=15 mice, n=42 sessions early in training, n=45 sessions late in training. Bar plots represent mean across mice ± SEM. Each circle represents one mouse (an average of 2-3 sessions early and 3 sessions late in training per mouse). For all panels: ∗*p* < 0.05.

Each trial has distinct epochs consisting of: 1) a variable delay “enforced no-lick (ENL) period” (1-2 seconds) in which mice are required to withhold their licks and which restarts if the animals licks during it; 2) an auditory “go cue” (75 ms, 5 kHz tone) whose end signals the beginning of the response epoch; 3) a “selection” epoch (up to 3 seconds long) in which mice report their choice action by licking the left or right spout; 4) an outcome evaluation epoch (3 seconds) in which the mice either receive and consume (win) or do not receive (lose) a water reward (**Figure 1A**). We refer to this outcome evaluation epoch as the “action-outcome association” period, as the outcome (presence or absence of a reward) is linked to the executed choice action (lick left or lick right). The optimal task strategy to maximize rewards is to implement win-repeat (WR) and lose-switch (LS) behaviors and to avoid win-switch (WS) and lose-repeat (LR) behaviors (**Figure 1B-C, E**) (Nowak and Sigmund, 1993). That is, if a mouse received a reward on one trial (i.e. a win) they should choose the same spout again on the next trial (repeat). Conversely, if they did not receive a reward (i.e. a loss), they should change the chosen spout (switch) on the next trial. Thus, the identity of the last action taken and its ensuing outcome fully determine the subsequent optimal action.

Mice learned the task structure within 4-6 weeks (**Figure 1B-D**). To quantify learning, we made within-mouse comparisons “early” or “late” in training (N=15 mice, 42 sessions early, 45 sessions late). Early sessions were those after 1 week of training (days 5-7), whereas late sessions were those after 4-6 weeks of training (**Methods**). Trained mice completed more trials per session (early: 158.3±10.4 vs. late: 221.5±4.2 trials, p=0.0001), achieved a higher percentage of rewarded trials (early: 61.2±2%; late: 90.0±1%, p=6.1e-05), and needed fewer trials to obtain a reward after the reward contingencies changed at block boundaries, or in other words, fewer trials to switch between spouts (early: 5.3±0.4; late: 1.4±0.1 trials, p=6.1e-05) (**Figure 1D**). Importantly, trained mice learned to withhold their licks between trials and had a lower percentage of trials with lick penalties during the ENL period (early: 57.7±3.4% vs. late: 20.9±2.8%, p=6.1e-05) or cue period (**Figure 1D; Supp Figure 1A**), indicating they learn to adhere to the trial structure. Well-trained mice were not biased in choosing the right or left spout and male and female mice performed the task to similar proficiency (**Supp Figure 1D-E**). **Table S1** summarizes all statistical comparisons.

In addition to learning the task structure, mice acquired the task strategy over the course of training (**Figure 1E-G**). Trained mice used the correct strategy, executing WR and LS sequences, on a larger fraction of trials compared to novice mice (early: 66.6±1.6% vs. late: 88.6%±1.1%, p=6.1e-05) (**Figure 1F; Supp Figure 1B**). Early-in-training mice correctly performed WR but failed to execute LS, instead persisting on the unrewarded spout (i.e. lose-repeat) (log(%LS/%LR) early: −0.47±0.1; late: 0.35±0.1, p=6.1e-05) (**Figure 1F**). Thus, mice quickly learned to choose the same spout as their previous choice after they receive a reward but required additional training to learn to behave flexibly and switch actions following a negative outcome. Finally, trained mice rarely switched their spout choices after a win (i.e., they rarely WS) (**Figure 1G; Supp Figure 1C**). Taken together, mice learn to implement WR and LS strategies and to withhold licks between trials as necessary to efficiently obtain rewards in this task.

### Choice is guided by the previous action-outcome association

The trial structure was designed to separate the action-outcome association on one trial (trial n) from the next choice action (trial n+1) by a variable delay period (**Figure 2A**, N=15 mice). In rewarded trials, well-trained mice licked more and over a longer duration compared to unrewarded trials, in which licking tapered off quickly following an unrewarded action (16.3±1.0 vs. 3.3±0.4 licks on trial n, p=6.1e-05) (**Figure 2B-C; Supp Figure 2A**). Furthermore, on “win” trials, almost all licks were directed toward the initial choice side (92.3±2.3% of licks), whereas the lick direction was more variable on “lose” trials (74.3±3.3% of licks toward the initial choice side) (**Figure 2D**). As the action-outcome association determines the optimal next choice, these directional changes following a loss may suggest that animals quickly begin updating their behavioral strategy for their next choice. Consistent with this possibility, licks during the ENL period were more likely to follow the direction that the animal would eventually choose in the subsequent trial (n+1), deemed the “chosen” side (**Figure 2E**). This effect was seen in ENL periods following both win (0.76±0.03: chosen vs. 0.24±0.03: not-chosen, p=6.1e-05) and lose trials (0.84±0.03: chosen vs. 0.16±0.0.3: not-chosen, p=6.1e-05). This finding persisted when restricting analysis to the first and last licks within a penalty-lick bout (**Supp Fig 2B**). This suggests that the motor plan for the next action is already beginning to be formed in the time between the last lick of the action-outcome association period and the start of the delay period. Further, it indicates that the licks during the ENL are interpreted differently from unrewarded licks following the go cue, confirming that the animals understand the task structure. Finally, the response time (measured from the end of the cue to the first lick) was slower for LS compared to WR trials (149±13ms vs. 111±7ms, respectively, p=2.0e-4) (**Figure 2F**) independent of the presence or absence of penalized ENL period licks (**Supp Figure 2C**). Thus, licks occurring during the inter-trial interval are informative of the animal’s upcoming choice and indicate that a representation of recent action-outcome experience may be maintained across trials.

**Figure 2.**
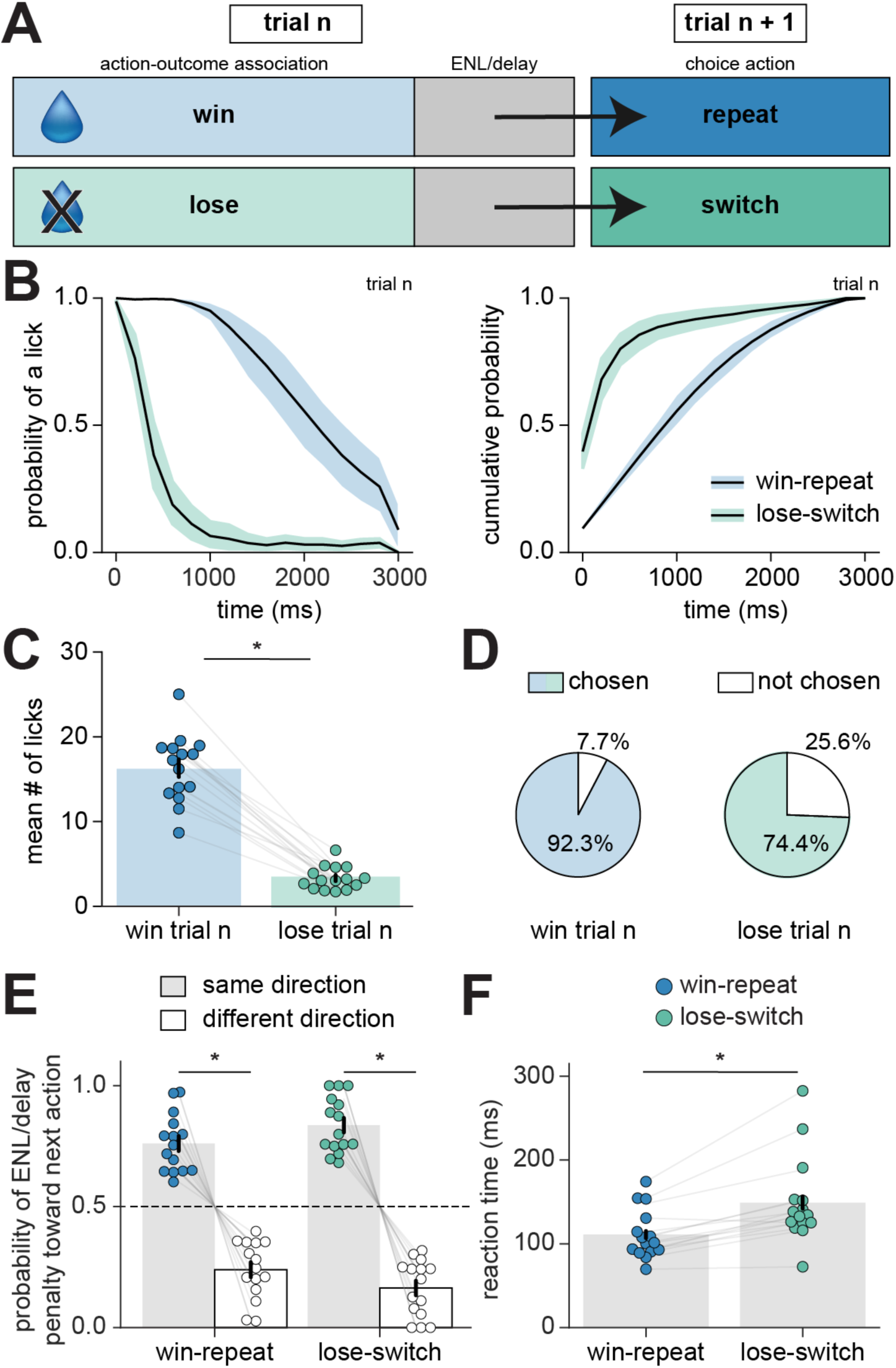
Choice is guided by the previous action-outcome association. **A.** Schematic of the optimal task strategies (win-repeat and lose-switch) across two trials (trial n and n+1). The action-outcome association period (trial n) informs the choice action of the next trial (trial n+1). **B.** Probability distribution (*left*) and cumulative probability distribution (*right*) of licking across the action-outcome association period in 200 ms bins for win-repeat (blue) and lose-switch (green) trials. t=0 represents the choice lick. Solid lines are the average of mice; shading represents 95% confidence. **C.** Average number of licks on win-repeat (blue) and lose-switch (green) trials during the action-outcome association period on trial n. **D.** Pie charts depicting the percentages of licks during the action-outcome association period (trial n) in the direction that was the same (chosen) or different (not chosen) as the choice action on trial n for win-repeat (blue) and lose-switch trials (green). Percentages represent averages across mice. **E.** On trials with lick penalties during the delay period, the probability that a lick penalty occurs in the direction of the next choice action (grey) or not (white) is shown for win-repeat (left bars, blue dots) and lose-switch (right bars, green dots) trials. **F.** Average response times (from the start of the cue until the first lick) for win-repeat (blue dots) and lose-switch (green dots) on trial n+1. Each dot represents the average reaction time per mouse. **A-F**. N=15 mice, n=45 sessions late in training. Bar plots represent mean across mice ± SEM. Each circle represents one mouse. For all panels: ∗*p* < 0.05.

### Dopamine in the ventrolateral striatum encodes reward and action history

The mouse behavior described above suggests that updating or switching the motor plan between trials is triggered by a loss event. As block transitions are uncued, unexpected outcome changes at these transitions likely evoke dopamine (DA) transients in the striatum that reflect reward prediction errors (RPE) (Schultz et al., 1997; Bromberg-Martin et al., 2010). To quantify the magnitude of RPE, we used fiber photometry to monitor DA release in mice performing the dynamic decision-making task. We used adeno-associated viruses (AAVs) to express red GRAB_DA_ (AAV9-hSyn-RDAm, abbreviated rDAm) (Sun et al., 2020), a sensor that serves as a proxy of DA release, in the ventrolateral striatum (VLS) of *Drd1a-Cre* (dSPNs) or *Adora2a-Cre* (iSPNs) mice and implanted optical fibers (**Figure 3A-B**, N=17, 6 D1-Cre, 11 A2a-Cre). While our virus is Cre-independent, we performed these experiments in the same BAC lines used for pathway-specific dSPN and iSPN (*Drd1a-Cre* or *Adora2a-Cre,* respectively) calcium recordings to match behavioral and genetic conditions. Coordinates were selected to target a region of VLS previously shown anatomically and functionally to contribute to orofacial, including tongue, movements (Lee et al., 2020; Bakhurin et al., 2020; Lee and Sabatini, 2021; Chen et al., 2021; Tang et al., 2021). All analyses were performed on z-scored DA signals aligned to task-specific events (**Figure 3C-D; Methods**). The amplitudes of DA transients were quantified by aligning each trial to the event of interest (e.g., ENL delay period, cue, selection lick, or outcome lick) and calculating the peak (ΔF_peak_) and average signal (ΔF_avg_) within a fixed time window after the event (**Supp Figure 3A-B**). Consistent with previous reports (Beron and Sabatini, 2023; Chantranupong et al., 2023), we saw no systematic differences in DA encoding across the right and left hemispheres (**Supp Figure 3A-B**); therefore, data were pooled across hemispheres and grouped according to whether licks were to the spout ipsilateral or contralateral to the fiber optics.

**Figure 3.**
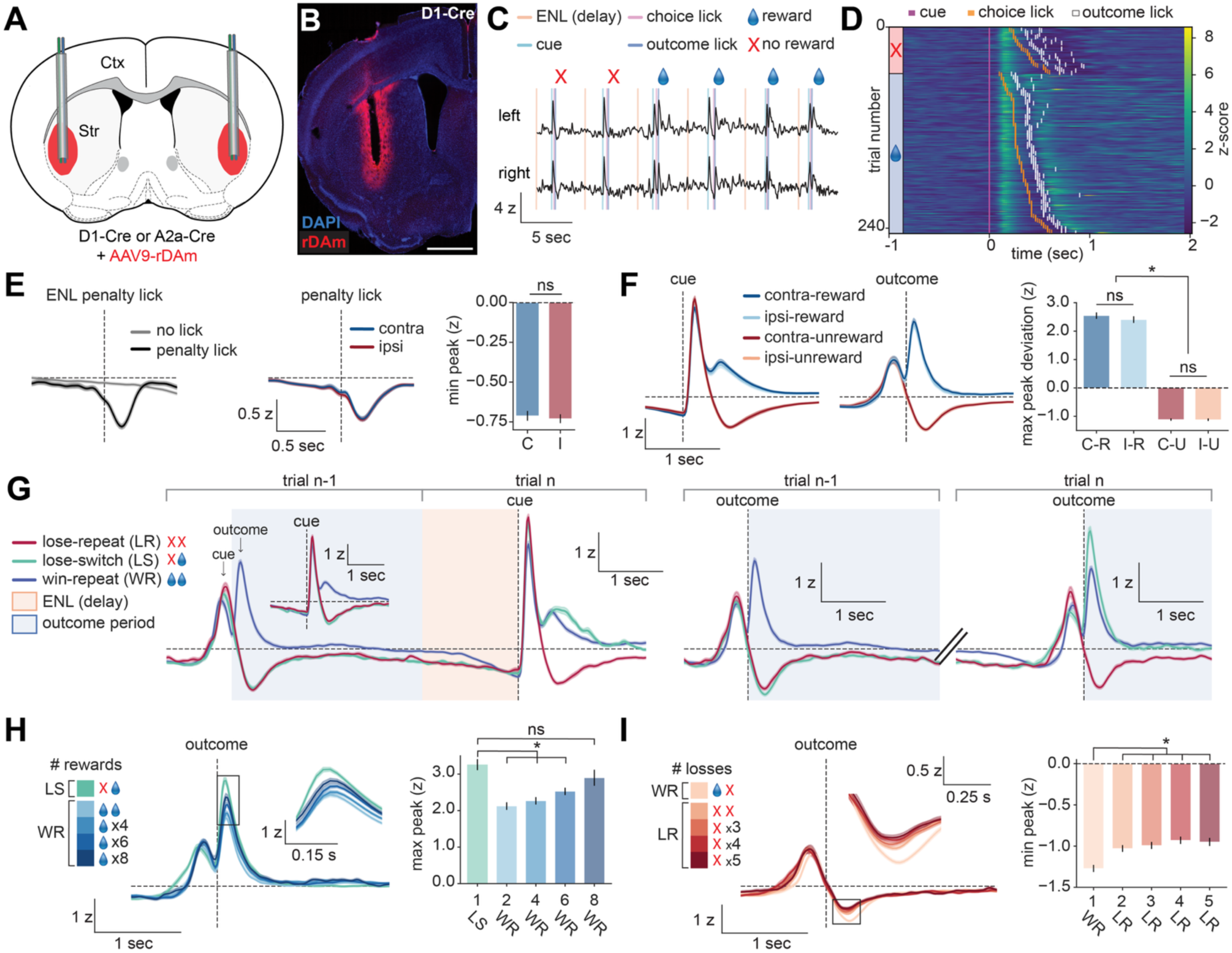
Dopamine in the ventrolateral striatum encodes reward and action history. **A.** Coronal schematic of the experimental approach showing bilateral AAV9-hSyn-rDAm viral expression and fiber implantation targeting the ventrolateral striatum. **B.** Low-magnification image showing rDAm expression (red) and DAPI fluorescence (blue) in the striatum. Scale bar: 1 mm. **C.** Representative z-scored photometry traces of striatal DA recorded bilaterally from the right and left hemisphere during behavior. Trial specific events (ENL/delay start, cue, choice lick, and outcome lick) are denoted. Symbols: water drop represents a rewarded trial; X represents an unrewarded trial. **D.** Representative behavioral session showing DA dynamics aligned to cue onset (pink). Each row represents one trial (z-scored signal). The timing of choice action licks (orange) and outcome licks (white) are denoted. Trials are sorted by lick latency and rewarded (blue with water symbol) or unrewarded (red with X) trial outcomes. **E.** DA dynamics aligned to spontaneous penalty licks during the ENL period. *left*: Activity with (black) and without (grey) penalty licks aligned to the first penalty lick during the ENL period. *middle*: DA dynamics during contralateral (C, blue) and ipsilateral (I, red) penalty licks. *right*: minimum z-score amplitude following penalty lick onset. **F.** DA dynamics aligned to the cue or outcome lick during a correct (rewarded) or incorrect (unrewarded) choice for licks to the contralateral (dark blue/dark red) or ipsilateral (light blue/orange) spout. *right:* Quantifications of the maximum or minimum z-score deviation of the response magnitudes following the outcome lick for contralateral (C) vs. ipsilateral (I) and rewarded (R) vs. unrewarded (U) trial types. **G.** DA dynamics across consecutive trials as a function of outcome history. *left:* DA signals aligned to the cue on trial n, with activity from the preceding trial (n−1) shown to illustrate DA dynamics during the prior outcome period and subsequent ENL period. Inset shows DA signals aligned to the cue on trial n-1. *right:* DA signals aligned to the outcome lick on trial n-1 and trial n. Traces are grouped by trial history: win-repeat (WR), lose-repeat (LR), and lose-switch (LS). Blue shading denotes the outcome period; orange shading denotes the ENL period. **H.** DA dynamics aligned to the outcome lick following the indicated number of consecutive rewarded choice actions. The inset highlights that the first rewarded action reaches the maximum peak z-score. *right:* Quantification of the maximum peak z-score amplitudes of consecutive rewarded choice actions. **I.** DA dynamics aligned to the outcome lick following the indicated number of unrewarded choice actions. The inset highlights that the first unrewarded action reaches the minimum peak z-score. *right:* Quantification of the minimum peak z-score amplitudes of consecutive unrewarded choice actions. **A-I**. N=17 mice (6 D1-Cre and 11 A2a-Cre), n=110 sessions (60,276 trials). Signals are z-scored, aligned to event of interest, and averaged across mice. Shaded bars around mean signal represents SEM. Bar plots represent mean across mice ± SEM. For all panels: ns, not significant; ∗*p* < 0.05.

As reported previously, DA transients showed two dominant phases within a trial (Schultz et al., 1997; Menegas et al., 2017; Amo et al., 2022). The first was a positive DA peak immediately following the cue, whereas the second was an outcome-dependent positive or negative deviation that followed the first outcome lick (**Figure 3C-D, F; Supp Figure 3B**). ENL delay penalty licks, which were not rewarded and extended the inter-trial interval, were associated with negative deflections that were identical for ipsilateral and contralateral choices (ΔF_peak_: contra: −0.71±0.03, ipsi: −0.73±0.02, p=0.21) (**Figure 3E; Supp Figure 3A**) (Beron and Sabatini, 2023). Thus, although all unrewarded licks can evoke a DA dip, only dips that occur in the appropriate behavioral context appear to update the upcoming motor plan. The previously described propensity of most ENL delay licks to predict the animal’s upcoming choice direction is consistent with this interpretation (**Figure 2D**). Further, neither ΔF_peak_ nor ΔF_avg_ of ENL or cue associated DA transients differed for ipsilateral vs. contralateral licks, indicating that DA signals were not lateralized by lick direction (**Figure 3E; Supp Figure 3A**) (Beron and Sabatini, 2023), in contrast to prior studies showing that DA axons in the dorsomedial striatum preferentially encode contralateral movements (Parker et al., 2016; Tsutsui-Kimura et al., 2020; Hart et al., 2024). As expected, DA during the outcome-associated period was strongly modulated by the presence or absence of reward (**Figure 3F**). On rewarded trials, DA transients rapidly increased following the first outcome lick, whereas on unrewarded trials DA decreased or “dipped” in the same period (ΔF_peak_: contra-rewarded: 2.54±0.1, ipsi-rewarded: 2.45±0.1, p=0.1; contra-unrewarded: −1.12±0.04, ipsi-unrewarded: −1.11±0.04, p=1.0) (**Figure 3F; Supp Figure 3C**). Together, these results show that DA transients in this motor region of the striatum are neither simply motor-related nor lateralized. Instead, their bidirectional encoding of reward receipt and omission may provide an outcome signal that informs subsequent choices The task structure required the mouse to combine and retain information about action and outcome across the inter-trial interval to determine the next action. Although there is a simple optimal strategy, mice occasionally deviate from it by repeating choices after loss outcomes (i.e. lose-repeat), which inevitably leads to a second loss. We examined DA signaling across different types of trial pairs (WR-win, LS-win, and LR-lose) to understand which aspects of action, outcome, and future outcome (perhaps expected) are influenced by and potentially determine action/outcome sequences. To standardize the timing of behavioral events, we isolated pairs of trials with an ENL period of 1.5 s and no ENL penalty licks (**Figure 3G**). Compared to loss trials, win trials were followed by sustained DA signals that remained elevated after the outcome period, revealing a prolonged outcome-dependent signal during the inter-trial interval (Beron and Sabatini, 2023). Furthermore, the outcome on trial n-1 affected the magnitude of DA transients on the current trial (trial n), with LS-win trials, in which animals received reward after a prior loss, evoking larger positive DA responses than WR-win trials (**Figure 3G**).

We extended this analysis to examine how DA responses evolved across longer reward and loss sequences. The reward-evoked peak DA signal was larger for the first reward following a loss (LS-win trial) than for consecutive rewarded trials (ΔF_peak_: 3.3±0.2, 2.1±0.1, 2.3±0.1, 2.5±0.1, or 2.9±0.2 for repeated win number 1, 2, 4, 6, or 8 in a series) (**Figure 3H; Supp Figure 3D**). This is consistent with DA signaling reflecting an RPE-like signal, in which the first reward following a loss is unexpected relative to recent experiences (Schultz et al., 1997; Schultz and Romo, 1988; Bromberg-Martin et al., 2010). Interestingly, the outcome-evoked DA peak consistently increased following longer reward sequences after the first repeated reward (**Figure 3H**). This pattern may reflect sensitivity to the distribution of block lengths: as rewards accumulate, animals may increasingly expect a block switch, making continued reward delivery on the same spout relatively unexpected. A complementary pattern was observed during consecutive losses. The first single loss following a reward (WR-lose, which occurs at the block boundary) evoked the largest negative DA deflection compared to subsequent repeated losses evoked by serial LR-lose trials (ΔF_peak_: −1.3±0.04, −1.0±0.04, −1.0±0.04, −0.93±0.04, or −0.95±0.05 for loss number 1, 2, 3, 4, or 5 in a series) (**Figure 3I; Supp Figure 3E**). Altogether, these results indicate that outcome-evoked DA does not simply report the current outcome, but rather reflects the outcome in the context of recent reward history, consistent with an expectation-dependent signal shaped by the animal’s evolving estimate of block structure. Furthermore, under the assumption that DA transients during the action-outcome association period reflect RPEs, these analyses suggest that RPEs are largest when reward outcomes change and diminish with repeated experience of the same outcome.

### dSPN and iSPN calcium activity encodes action-specific outcome value and reward history

dSPNs and iSPNs are the primary output neurons of the striatum and are major postsynaptic targets of midbrain DA neurons (Gerfen and Surmeier, 2011). DA is expected to differentially modulate the excitability of Type 1 DA receptor-expressing dSPNs and Type 2 DA receptor-expressing iSPNs (Gerfen et al., 1990; Tritsch and Sabatini, 2012; Lahiri and Bevan, 2020), raising the possibility that outcome-and history-dependent DA signals could be transformed into pathway-specific representations of action value (Shin et al., 2018; Maltese et al., 2021). We therefore hypothesized that dSPNs and iSPNs encode action choice and outcome in a history-dependent manner.

To examine this possibility, we used fiber photometry to record population calcium activity from dSPNs or iSPNs in the VLS of *Drd1a-Cre* or *Adora2a-Cre* mice, respectively, across behavior (**Figure 4A**, N=8 D1-Cre, N=18 A2a-Cre). An AAV encoding FLEX-jGCaMP8m (AAV9-syn-FLEX-jGCaMP8m-WPRE) (Zhang et al., 2023) was bilaterally injected into the VLS and optical fibers were implanted above the injection sites. Post hoc histology confirmed targeting and viral expression (**Figure 4B; Supp Figure 4A**). High signal-to-noise fluorescence transients were observed around task-relevant behavioral events that were similar across hemispheres (**Figure 4C**). All analyses were performed on z-scored calcium signals aligned to task-specific events, and quantification was performed as described above (**Supp Figure 4B-C; Methods**). Like the DA transients, signals in the left and right hemispheres did not differ for dSPNs or iSPNs (**Supp Figure 4B-C, F-G)**. Thus, we pooled data across hemispheres and grouped data for each cell class according to whether licks were to the spout ipsilateral or contralateral to the fiber optics.

**Figure 4.**
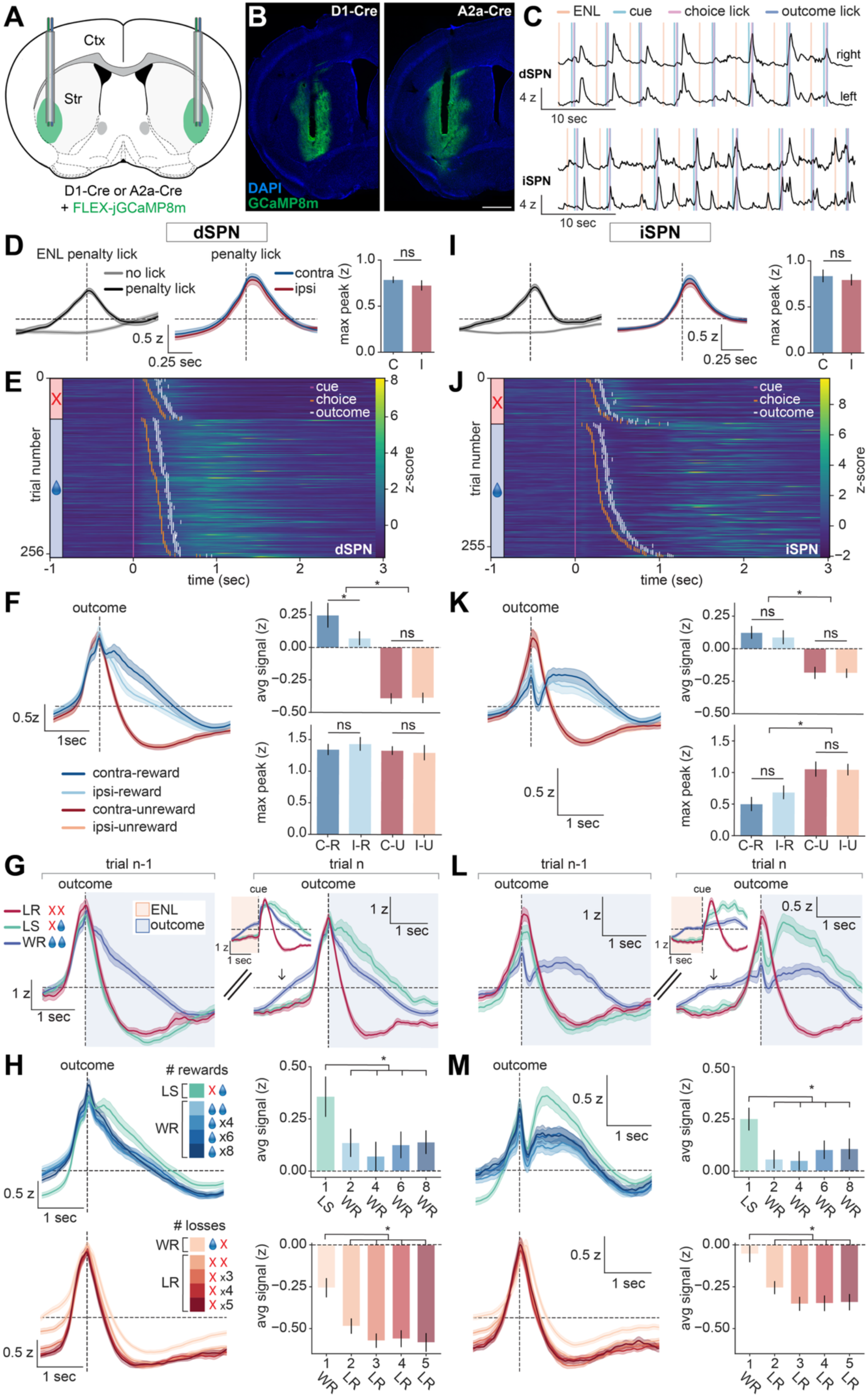
dSPN and iSPN calcium activity encodes action-specific outcome value and reward history. **A.** Coronal schematic of the experimental approach showing bilateral AAV9-FLEX-jGCaMP8m viral expression and fiber implantation targeting dSPNs or iSPNs in the ventrolateral striatum. **B.** Low-magnification image showing GCaMP8m expression (green) and DAPI fluorescence (blue) in dSPNs or iSPNs. Scale bar: 1 mm. **C.** Representative z-scored photometry traces from dSPNs (*top*) or iSPNs (*bottom*) recorded bilaterally from the right and left hemisphere during behavior. Trial specific events (ENL/delay start, cue, choice lick, and outcome lick) are denoted. **D.** Calcium activity in dSPNs aligned to spontaneous penalty licks during the ENL period. *left:* Activity with (black) and without (grey) penalty licks aligned to the first penalty lick during the ENL period. *middle*: Activity during contralateral (C) and ipsilateral (I) penalty licks. *right*: Peak z-score amplitude at penalty lick onset. **E.** Representative behavioral session showing calcium activity in dSPNs aligned to cue onset (pink). Each row represents one trial (z-scored signal). The timing of choice action licks (orange) and outcome licks (white) are denoted. Trials are sorted by lick latency and rewarded (blue with water symbol) or unrewarded (red with X) trial outcomes. **F.** *left:* Calcium activity in dSPNs aligned to the outcome lick during a correct (rewarded) or incorrect (unrewarded) choice for licks to the contralateral (dark blue/dark red) or ipsilateral (light blue/orange) spout. *right:* Quantifications (*top*: average z-scored signal; *bottom*: peak z-score) of the response magnitudes for contralateral (C) vs. ipsilateral (I) and rewarded (R) vs. unrewarded (U) trial types. **G.** Calcium activity in dSPNs across consecutive trials as a function of outcome history. Traces are aligned to the outcome lick on trial n-1 and trial n. Inset shows activity in dSPNs signals aligned to the cue on trial n. Activity diverges according to trial history before the outcome lick on trial n (see arrow). Traces are grouped by trial history: win-repeat (WR), lose-repeat (LR), and lose-switch (LS). Blue shading denotes the outcome period. **H.** *left*: Calcium activity in dSPNs aligned to the outcome lick following the indicated number of consecutive rewarded (*top,* blue) or unrewarded (*bottom,* red) choice actions. *right:* Quantifications (*top/bottom*: average signal) of the response magnitudes of consecutive rewarded (*top*) or unrewarded (*bottom*) choice actions. **I-M**. As in E-H, except for iSPN calcium recordings. **A-M**. N=8 D1-Cre mice, n=65 sessions (35,538 trials). N=18 A2a-Cre mice, n=120 sessions (65,846 trials). Signals are z-scored, aligned to event of interest, and averaged across mice. Shaded bars around mean signal represents SEM. Bar plots represent mean across mice ± SEM. For all panels: ns, not significant; ∗*p* < 0.05.

We found that dSPNs were active at times of spontaneous licks, with positive signal deflections that occurred during the ENL period and did not differentiate between ipsilateral or contralateral movements (**Figure 4D; Supp Figure 4B**). Thus, in this context, dSPN activity reflected motor actions but was not lateralized, similar to DA dynamics. dSPN calcium was modulated by task-related events, increasing following the cue, through the choice lick, and peaking at the time of the outcome lick at which the reward outcome is revealed (**Figure 4E; Supp Figure 4C**). The peak (as reflected in ΔF_peak_) calcium signals in dSPNs aligned to the outcome lick displayed little contextual sensitivity and was similar for rewarded and unrewarded trials to the ipsilateral and contralateral side (**Figure 4F**), as seen previously (Tang et al., 2021). In contrast, the average calcium signal in the action outcome-association period (as reflected in ΔF_avg_) was highly dependent on outcome and choice direction: ΔF_avg_ was larger for rewarded contralateral than ipsilateral trials (ΔF_avg:_ contra-rewarded: 0.24±0.1, ipsi-rewarded 0.06±0.06, p=0.03) (**Figure 4F**). Both were larger than the net negative ΔF_avg_ seen in unrewarded trials, which was insensitive to action direction (ΔF_avg_: contra-unrewarded: −0.40 ± 0.04, ipsi-unrewarded: −0.39±0.04, p=0.74) (**Figure 4F**). The effect persisted when controlling for reward history by analyzing consecutive same-spout selections but differing by rewarded outcomes (**Supp Figure 4D**).

To contextualize these patterns with our DA results, we examined dSPN activity across consecutive trials by comparing WR-win, LS-win, and LR-lose trials (**Figure 4G**). When aligned to the outcome lick on trial n-1, dSPN activity differed according to the outcome, with rewarded trials producing more sustained activity compared to unrewarded trials. Although activity in win and loss trials converged by the end of the outcome period of trial n-1, dSPN activity increased again *before the cue and outcome lick on trial n*, indicating that recent outcome history shapes dSPN activity as the timing of the subsequent choice lick approaches (**Figure 4G**). Following the outcome lick, dSPN activity was elevated in LS-win trials, in which mice received reward after a prior loss, but was suppressed in LR-lose trials, in which mice experienced repeated unrewarded outcomes (**Figure 4G**). These results indicate that dSPN population activity reflects not only the current outcome, but also the recent sequence of outcomes leading to that trial.

We extended this analysis to examine how dSPN calcium responses evolved across longer reward and loss sequences. As with reward and loss responses overall, we found that average (ΔF_avg_), but not peak (ΔF_peak_), calcium signals in dSPNs were sensitive to the history of outcomes on previous trials (**Figure 4H**). The reward-evoked average calcium signal was larger for the first reward after a LS-win trial than for subsequent consecutive rewarded trials (ΔF_avg_: 0.36±0.1, 0.14±0.07, 0.07±0.07, 0.13±0.07, or 0.14±0.06 for win numbers 1, 2, 4, 6, or 8 in a series) (**Figure 4H; Supp Figure 4E)**, consistent with the representation of RPE-like signals conveyed from DA. Furthermore, the same outcome for the same action becomes expected after the first reward. In contrast, the first single loss following a reward (WR-lose trials, typically at the block boundary) evoked the smallest negative deflection in calcium signal, which became progressively more negative over ∼3 consecutive losses (ΔF_avg_: −0.26±0.06, −0.48±0.05, −0.57±0.05, −0.56±0.05, or −0.58±0.06 for loss numbers 1, 2, 3, 4, or 5 in a series) (**Figure 4H; Supp Figure 4E**), inconsistent with RPE-like encoding conveyed from DA during repetitive loss trials. Notably, the pre-cue baseline calcium levels *before the subsequent cue and outcome* are suppressed following a single and multiple losses, suggesting cumulative suppression across trials and that traces of previous outcome history remain during the inter-trial interval (**Figure 4H**). Together, these results indicate that dSPN activity is not determined solely by the current outcome but is shaped by outcome history in a manner that is partly consistent with RPE-like signaling for rewarded outcome, but not for losses.

We performed analogous recordings and analyses in iSPNs. As with dSPNs, spontaneous licks correspond to robust activity in iSPNs that was insensitive to lick direction (**Figure 4I; Supp Figure 4F**). Co-activation of dSPNs and iSPNs during spontaneous licking suggests that motor actions are encoded by joint, rather than opponent, pathway activity (Cui et al., 2013; Isomura et al., 2013; Jin et al., 2014; Barbera et al., 2016; Parker et al., 2018; Markowitz et al., 2018). Task-related events were also robustly modulated calcium activity in iSPNs (**Figure 4J; Supp Figure 4G**) but the contextual sensitivity differed between classes. Unlike in dSPNs, peak outcome calcium signals in iSPNs are context sensitive and are higher on unrewarded trials compared to rewarded trials for both contralateral and ipsilateral licks (ΔF_peak:_ contra-unrewarded: 1.06±0.12, ipsi-unrewarded: 1.05±0.09, contra-rewarded: 0.5±0.12, ipsi-rewarded: 0.69±0.11) (**Figure 4K**), consistent with previous findings in the dorsomedial striatum (Nonomura et al., 2018). This effect persisted when controlling for reward history (**Supp Figure 4H**). Average calcium signal showed the opposite effect and was larger for rewarded trials due to a prolonged signal that emerged later in the trial (**Figure 4K**). As for dSPNs, across consecutive rewards and losses, calcium activity in iSPNs also showed history-dependent modulation in the average calcium signal across multiple trials (**Figure 4L-M**). Small but significant changes were also observed in the peak amplitude when comparing consecutive rewards (**Supp Figure 4I; Table S1**). Finally, shifts in baseline activity dependent on the outcome of the previous trial were also observed, indicating that iSPNs, like dSPNs, track recent outcome history across trials (**Figure 4L-M**).

Together, these results demonstrate that dSPN and iSPN population activity encodes action-outcome associations and recent outcome history in pathway-specific manners. dSPNs preferentially encode rewarded contralateral actions, whereas iSPNs show enhanced peak responses to unrewarded outcomes without clear action-direction specificity. Although these signals did not simply mirror DA, both pathways exhibited history-dependent modulation across consecutive rewards and losses. The history dependence of pre-cue baseline levels further suggests that SPN population activity carries information about recent outcomes across trials. Thus, downstream of outcome- and history-dependent DA signaling, dSPN and iSPN activity are plausible candidates to maintain action-outcome information across the inter-trial interval to guide upcoming choices.

### Future action and recent outcome history can be decoded from dSPN and iSPN population activity

Photometry recordings showed that dSPNs and iSPNs were modulated by both reward- and action-related variables, including movement execution and choice direction, and exhibited outcome-dependent baseline shifts across successive trials. Given these results and prior work involving striatal activity in value-based decision making (Tai et al., 2012; Donahue, 2018; Nonomura et al., 2018; Shin et al., 2018; Tang et al., 2021; Bolkan et al., 2022; van Beest et al., 2024), we hypothesized that striatal population activity would encode recent action-outcome history and carry information relevant to subsequent choices. We therefore asked whether SPN activity predicted future actions, consistent with a role in guiding upcoming behavior, and encoded past trial outcomes, consistent with a representation of the action–outcome history that could inform those choices. First, we confirmed that outcome-period calcium activity robustly discriminated the outcome of the ongoing trial, with logistic regression classifiers decoding win (reward) vs. lose (unrewarded) trials with peak receiver operating characteristic area under the curve (ROC AUC) values of 0.82 for dSPNs and 0.76 for iSPNs (chance=0.5; **Supp Figure 5A-D; Methods**). To test future-action decoding, we examined whether striatal calcium activity during the outcome period of an unrewarded trial (trial n) predicts whether the animal will repeat or switch its action on the next trial (n+1). This analysis was performed on loss trials, following which animals either update their strategy and switch to the opposite spout (LS-win, rewarded on trial n+1) or erroneously repeat the same action (LR-lose, unrewarded on trial n+1) (**Figure 5A**). The low rate of WS trials precluded a parallel analysis of future-action decoding following rewarded trials.

**Figure 5.**
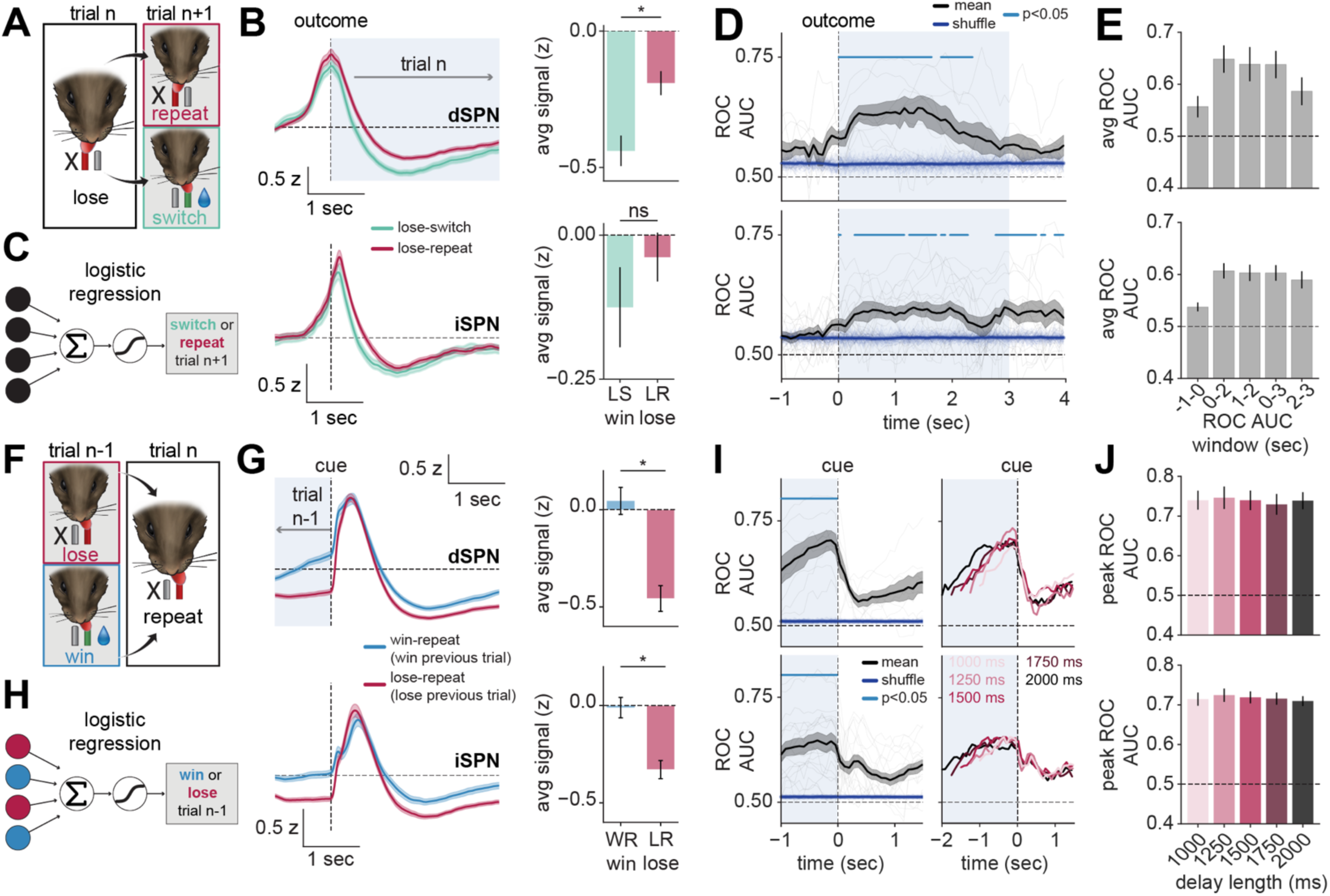
Future action and recent outcome history can be decoded from dSPN and iSPN population activity. **A.** Behavioral schematic illustrating lose-repeat and lose-switch trials. A loss on trial n is followed on trial n+1 by either repeating the same action and losing again (LR-lose) or switching actions and receiving reward (LS-win). **B.** *left*: Calcium activity in dSPNs (*top*) or iSPNs (*bottom*) aligned to the outcome lick on trial n, segregated by next-trial behavior (switch, green; repeat, magenta). Shaded region denotes analysis window to predict next-trial choice (trial n+1) for dSPNs and iSPNs. *right:* Quantification of the average z-scored signal (trial n window) for lose-switch (LS-win) and lose-repeat (LR-lose) conditions. **C.** Schematic of a logistic regression classifier trained to predict next-trial choice behavior from striatal activity during the outcome period of trial n (switch or repeat on trial n+1). To control for trial n outcome, the classifier was trained only on loss trials, comparing lose-switch and lose-repeat trials. **D.** Time-resolved ROC AUC analysis quantifying decoding of next-trial choice (switch vs. repeat) from calcium in dSPNs (*top*) or iSPNs (*bottom*). Thick black line: mean ROC AUC; thin black lines: individual mice; thick blue line: mean shuffle, thin blue lines: individual shuffles. Shaded region indicates analysis window. Significant time points are indicated (cyan line, *p* < 0.05, see Methods). **E.** Average mean ROC AUC across varying time windows for distinguishing switch vs. repeats actions on trial n+1 from the dSPN (*top*) and iSPN (*bottom*) signal during the trial n outcome period. **F.** Behavioral schematic illustrating lose-repeat and win-repeat trials. On trial n, the same action is repeated following either a loss (lose-repeat) or a win (win-repeat) on trial n-1. Both conditions result in a loss on trial n, isolating the effect of previous outcome history. **G** Direct (*top/left*) or indirect (*bottom/left*) pathway activity aligned to the cue on trial n, segregated by previous trial outcome (win-repeat, blue; lose-repeat, magenta). Shaded region denotes the analysis window used to predict previous-trial outcome (trial n-1) for dSPNs and iSPNs. *right*: Quantification of the average z-scored signal (trial n-1 window) for win-repeat (WR win) and lose-repeat (LR lose) conditions. **H.** Schematic of a logistic regression classifier trained to predict previous-trial outcome from striatal activity during the delay period of trial n (win or loss on trial n-1). To control for trial n choice and outcome, the classifier was trained only on loss trials in which animals repeated the previous choice, comparing win-repeat (WR-lose) and lose-repeat (LR-lose) trials. **I.** Time-resolved ROC AUC analysis quantifying decoding previous trial outcome from calcium activity in dSPNs (*top*) or iSPNs (*bottom*). *left*: ROC AUC across the 1 second period preceding cue onset and collapsed across delay lengths. Thick black line: mean ROC AUC; thin black lines: individual mice; thick blue line: mean shuffle, thin blue lines: individual shuffles. Shaded region indicates analysis window. Significant time points are indicated (cyan line, *p* < 0.05, see Methods). r*ight*: Mean ROC AUC across individual delay durations (1000 ms-2000 ms) for dSPNs (*top*) and iSPNs (*bottom*). **J**. Peak ROC AUC across delay lengths (1000ms-2000ms) for distinguishing previous trial outcome on trial n-1 from the dSPN (*top*) and iSPN (*bottom*) signal during the cue period. **A-J.** N=8 D1-Cre mice, n=65 sessions (35,538 trials). N=18 A2a-Cre mice, n=120 sessions (65,846 trials). Signals are z-scored, aligned to event of interest, and averaged across mice. Shaded bars around mean signal represents SEM. Bar plots represent mean across mice ± SEM. For all panels: ns, not significant; ∗*p* < 0.05. Abbreviations: ROC AUC: (receiver operating characteristic area under the curve).

For both dSPNs and iSPNs, calcium activity on the loss trial itself differed according to the animal’s choice on the next trial. Beginning at the outcome lick on trial n, before the animal had either switched or repeated its action on trial n+1, trials that would be followed by a switch were associated with a more negative signal, although this only reached statistical significance for dSPNs (ΔF_avg_ dSPN: LS-win: - 0.44±0.06 vs. LR-lose: −0.19±0.05, p=0.008, N=8 D1-Cre mice; ΔF_avg_ iSPN: LS-win: −0.13±0.07 vs. LR-lose: −0.04±0.04, p=0.074, N=18 A2a-Cre mice) (**Figure 5B**). This effect was consistent for ipsilateral and contralateral actions (**Supp Figure 5E**). To test whether this difference could be used to predict future behavior, we trained logistic regression classifiers to discriminate switch versus repeat actions on trial n+1 based on calcium activity during the outcome period of trial n (**Figure 5C**). Time-resolved ROC analysis revealed that decoding performance rose significantly above chance and above shuffled controls during the outcome window, peaking at 0.65 ROC AUC for dSPNs and 0.60 ROC AUC for iSPNs (**Figure 5D**). Decoding performance was strongest within the first ∼2 seconds of the action-outcome association period, although the entire period represented above chance performance (**Figure 5E**). Thus, both dSPN and iSPN population activity during the action-outcome association period predicts future choice actions, with stronger predictive power observed in dSPNs.

We separately examined which aspects of trial history were encoded in the inter-trial interval calcium activity of iSPNs and dSPNs, and whether this activity was sufficient to decode recent outcome history, independent of current action. To isolate outcome history from motor output, we compared WR-lose and LR-lose trials. For both conditions, animals repeated the same action and experienced the same outcome on trial n, such that they only differ in whether the preceding trial (n-1) was rewarded or unrewarded (**Figure 5F**). For both dSPNs and iSPNs, average calcium signals in the 1 s before the cue were more negative for LR-lose vs. WR-lose trials (ΔF_avg_ dSPN: LR-lose: −0.46±0.07 vs. WR-lose: 0.04±0.07, p=0.008; ΔF_avg_ iSPN: LR-lose: −0.33±0.05 vs. WR-lose: 0.01±0.05, p=3.8e-5). These effects were again identical for ipsilateral and contralateral actions (**Figure 5G; Supp Figure 5F**). Logistic regression decoded previous-trial outcome from delay-period activity above chance across delay lengths, with peak ROC AUC values of 0.71 for dSPNs and 0.66 for iSPNs (**Figure 5G–J**). As in the future-choice decoding analysis, decoding performance was stronger in dSPNs than for iSPNs. Together, these findings indicate that striatal population activity carries information about both upcoming actions and recent outcomes across trials, positioning SPNs to support memory-guided decision making.

### Cell-type specific activation and inhibition of dSPNs and iSPNs during the action-outcome association period bidirectionally modulate action updating

Given that calcium activity in SPNs exhibited outcome-dependent dynamics that predicted future choices and also encode task-relevant history information (**Figure 5**), we hypothesized that activity in dSPNs and iSPNs during the action-outcome association period contributes causally to the short-term maintenance of information used to guide the next choice. Prior work has shown that transient activation of dSPNs and iSPNs can bias lateralized choices in opposite directions, consistent with pathway-specific effects on action selection and action value (Tai et al., 2012). We therefore reasoned that if VLS SPN activity maintains an upcoming motor choice memory across trials, then perturbing dSPNs or iSPNs during the action-outcome association period should bias the animal’s subsequent choice in pathway- and direction-specific ways. As the task allows for precise temporal separation of task-specific events, we used optogenetics to manipulate striatal activity specifically during the action-outcome association period. We expressed ChR2-mCherry or eYFP in dSPNs or iSPNs in the VLS and implanted tapered optical fibers bilaterally (Pisanello et al., 2014, 2017; Pisano et al., 2019) (**Figure 6A**). Post-mortem histology confirmed viral expression and targeting (**Figure 6B; Supp Figure 6A**). To test whether activity during the action-outcome association period is sufficient to influence the next choice, we trained mice to task proficiency and stimulated dSPNs or iSPNs on ∼15% of trials. Stimulation was delivered only on WR-win trials and was triggered by the correct outcome lick of trial n, *after* the choice lick and *during* the action-outcome association period (N=18 D1-ChR2, 4 D1-eYFP, 6 A2a-ChR2, 6 A2a-eYFP) (**Figure 6C; Methods**).

**Figure 6.**
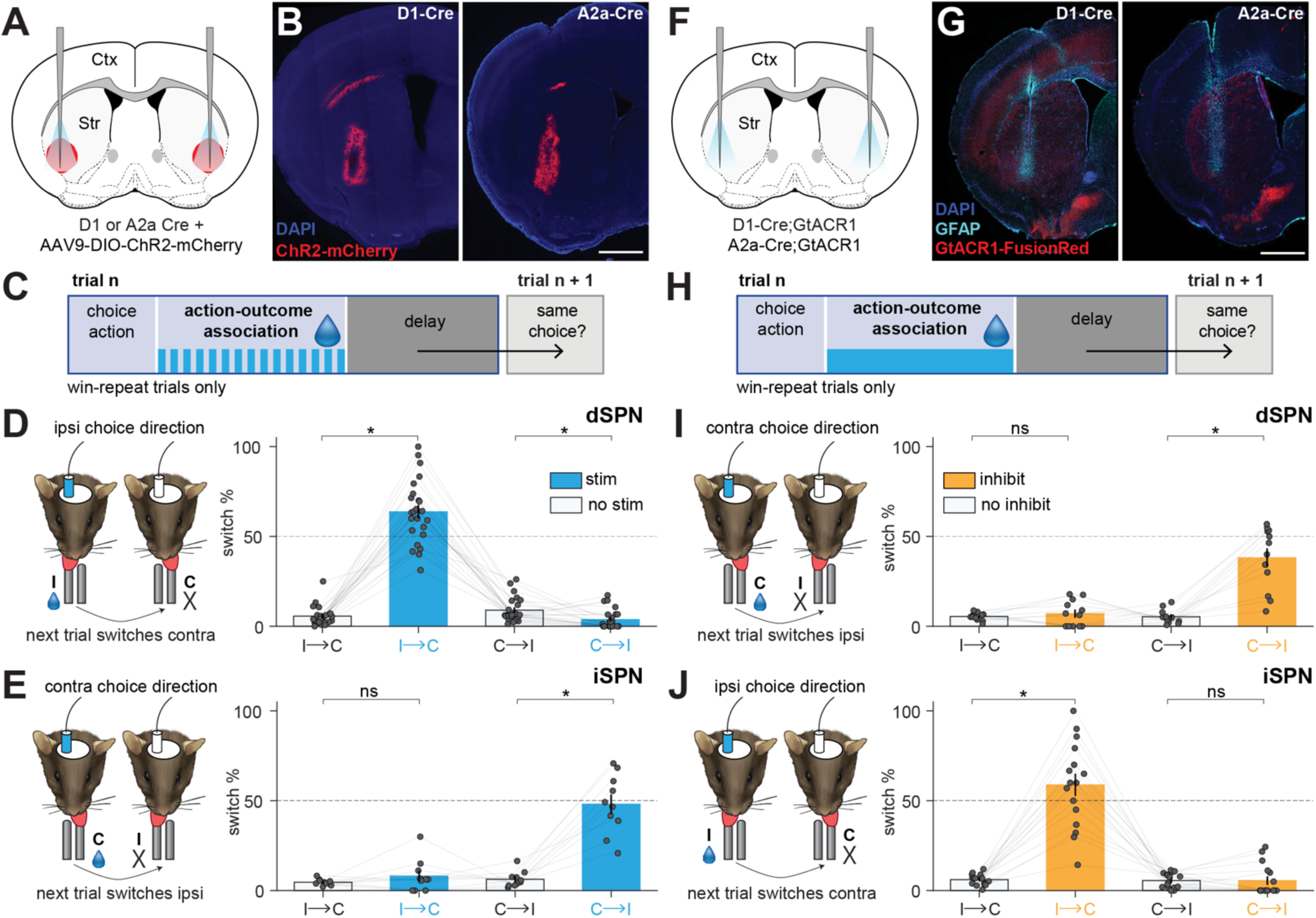
Cell-type specific activation and inhibition of dSPNs and iSPNs during the action-outcome association period bidirectionally modulate action updating. **A**. Coronal schematic showing viral targeting of AAV9-DIO-ChR2-mCherry and tapered fiber implantation in the ventrolateral striatum. **B** Representative low-magnification histology showing ChR2-mCherry (red) and DAPI fluorescence (blue) in dSPNs and iSPNs. Scale bar: 1 mm. **C.** Schematic of the task structure showing the period of optogenetic activation. Stimulation was delivered only on rewarded trials during the action-outcome association period of trial n (20 Hz, 1 mW, 3 seconds, 488 nm blue light). **D.** Schematic (*left*) and summary (*right*) of the effects that dSPN activation during rewarded trial n has on the action selected in trial n+1. dSPN stimulation promotes choice of the spout contralateral to the stimulated site such that it greatly increases the percent of ipsilateral to contralateral spout (I→C) reward-switch trials, quantified as switch percentage (switch %). Blue indicates stimulation trials; white indicates no stimulation trials. **E.** Same as in D, except for iSPNs. iSPN stimulation promotes choice of the spout ipsilateral to the stimulated site such that it greatly increases the fraction of contralateral to ipsilateral spout (C→I) reward-switch trials. **F.** Coronal schematic showing tapered fiber implantation in the ventrolateral striatum. **G.** Representative low-magnification histology showing GtACR1-FusionRed (red), GFAP (cyan), and DAPI fluorescence in dSPNs and iSPNs. Scale bar: 1 mm. **H.** Schematic of the task structure showing the period of optogenetic inhibition. Inhibition was delivered during the action-outcome association period only on rewarded trials (continuous inhibition, 0.5 mW, 3 seconds, 488 nm blue light). **I.** Schematic (*left*) and summary (*right*) of the effects that dSPN inhibition during rewarded trial n has on the action selected in trial n+1. dSPN inhibition promotes choice of the spout ipsilateral to the stimulated site such that it greatly increases the percent of contralateral to ipsilateral spout (C→I) reward-switch trials. Orange indicates inhibition trials; white indicates no inhibition trials. **J.** Same as in I, except for iSPNs. iSPN inhibition promotes choice of the spout contralateral to the stimulated site such that it greatly increases the fraction of ipsilateral to contralateral spout (I→C) reward-switch trials. **D-E, I-J.** N=18 mice, n=75 sessions (D1-ChR2); N=6 mice, n=32 sessions (A2a-ChR2); N=8 mice, n=27 sessions (D1;GtACR1); N=11 mice, n=24 sessions (A2a;GtACR1). Each dot represents one hemisphere per mouse. Bar plots represent mean across mice ± SEM. For all panels: ns, not significant; ∗*p* < 0.05.

dSPN stimulation during the action-outcome association period caused animals to select the unrewarded spout (contralateral to stimulation) on the next trial (D1-ChR2: I→C no stim: 5.6±1.1% vs. stim: 63.7±3.6%, p=2.4e-7; C→I no stim: 8.9±1.37% vs. stim: 3.7±1.1%, p=0.009), producing a robust increase in erroneous “win-switch” behavior when the stimulation was ipsilateral to the rewarded spout (**Figure 6D; Supp Figure 6B**). To determine whether this effect could be explained by changes in motor output during reward consumption, we examined licking during the stimulation on trial n. dSPN stimulation did not alter the total lick count (no stim: 13.6±0.6 vs. stim: 14.7±0.7 licks, p=0.17), but reduced lick frequency within a licking bout (no stim: 7.7±0.2 vs. stim: 6.2±0.2 Hz, p=5.96e-7) and shifted the proportion of licks during the consumption bout toward the non-chosen spout (chosen no stim: 0.86±0.01 vs. chosen stim: 0.76±0.02, p=2.5e-5) (**Supp Figure 6C**). Although reaction times appeared slower in erroneous “win-switch” (stim WS) trials compared to control WR trials (no stim), the difference was not significant (no stim WR: 146±9ms vs. stim WS: 200±42ms, p=0.32) (**Supp Figure 6C**).

iSPN activation produced a similar increase in erroneous “win-switch” behavior, but with an opposing directional bias. Whereas dSPN stimulation biased switching toward the contralateral spout, iSPN stimulation biased switching toward the ipsilateral unrewarded spout on the next trial (A2a-ChR2: I→C no stim: 4.6±0.6% vs. stim: 8.1±2.9%, p=0.75; C→I no stim: 6.4±1.4% vs. stim: 48.1± 5.2%, p=0.0004) (**Figure 6E; Supp Figure 6F**). Motor effects following stimulation were distinct from those observed in dSPNs. iSPN stimulation reduced the total lick count (no stim: 11.9±1.3 vs. stim: 6.2±0.5 licks, p=0.002), but did not change the lick frequency within a bout (no stim: 7.9±0.3 vs. stim: 8.8±0.7 Hz, p=0.16) or directional bias (chosen no stim: 0.94±0.02 vs. chosen stim: 0.90±0.03, p=0.13) (**Supp Figure 6G**). No changes in next trial action updating or lick metrics were observed in control D1-eYFP or A2a-eYFP animals with equivalent light delivery (**Supp Figures 6D-E, H-I**; **Table S1**).

These results demonstrate that increasing dSPN and iSPN activity during the action-outcome association period is sufficient to perturb future choice, with opposing influences across pathways. To determine whether this activity is also necessary for action updating during ongoing behavior, we expressed soma-localized GtACR1 in dSPNs or iSPNs by crossing the floxed-GtACR1 mouse (Li et al., 2019) to *Drd1a-Cre* or *Adora2a-Cre* mice, respectively (**Figure 6F-G**). We confirmed GtACR1 expression in SPNs with postmortem histology, slice physiology demonstrating light-evoked inhibition of spiking at low light powers, and confocal imaging verifying GtACR1 specificity to SPNs and not striatal interneurons (**Supp Figure 7A-F**). Inhibition of SPNs activity during the action-outcome association period of rewarded trials also disrupted action updating, but in the *opposite direction* of activation. dSPN inhibition biased animals to switch to the unrewarded ipsilateral spout (D1;GtACR1: I→C no stim: 5.4±0.6% vs. stim: 7.0±2.1%, p=0.67; C→I no stim: 5.3±1.1% vs. stim: 38.1±5.0%, p=0.001), whereas iSPN inhibition biased switching toward the unrewarded contralateral spout on the next trial (A2a;GtACR1: I→C no stim: 6.1±0.7% vs. stim: 58.8±5.9%, p=6.1e-5; C→I no stim: 5.6±1.0% vs. stim: 5.5±2.1%, p=0.82) (**Figure 6H-J; Supp Figure 7B, D**).

As with activation, we assessed changes in motor output associated with inhibition. dSPN inhibition reduced total lick count (no stim: 15.2±0.8 licks vs. stim: 9.9±1.1 licks, p=0.0005) without affecting lick frequency (no stim: 7.5±0.2 Hz vs. stim: 7.8±0.1 Hz, p=0.13) or direction (chosen no stim: 0.88±0.02 vs. chosen stim: 0.88±0.05, p=0.39; not chosen no stim: 0.12±0.02 vs. not chosen stim: 0.12±0.05, p=0.38) (**Supp Figure 7H**). iSPN inhibition did not alter the total lick count (no stim: 12.9±1.0 licks vs. stim: 13.1±1.0 licks, p=0.78), but reduced lick frequency (no stim: 7.1±0.2 vs. stim: 5.8±0.2 Hz, p=0.0003) and shifted the proportion of licks during the consumption bout toward the non-chosen spout (chosen no stim: 0.89±0.02 vs. chosen stim: 0.75±0.02, p=0.001; not chosen no stim: 0.11±0.02 vs. not chosen stim: 0.25±0.02, p=0.001) (**Supp Figure 7J**). Inhibition did not change the next trial reaction time for either pathway (**Supp Figure 7H, J**). No changes in action updating or lick metrics (lick count, frequency, direction, or reaction time) were observed in Cre-negative floxed-GtACR1 control animals (**Supp Figures 7K-L; Table S1**).

### Cell-type specific inhibition of dSPNs and iSPNs during the delay period bidirectionally modulates action updating

Our results, thus far, are consistent with SPN activity holding a memory of the motor plan in the inter-trial interval that determines future action selection. To more rigorously test this interpretation, we manipulated dSPNs and iSPNs during the ENL/delay period separating the action-outcome association (trial n) from the subsequent choice (trial n+1) (**Figure 7A-C; Supp Figure 8A**). During this time, animals were required to withhold actions; any frank movements or premature licks triggered an ENL penalty, which restarted the delay period. Thus, manipulations during this period allowed us to separate a motor plan memory from ongoing movement. Similar to the results of action-outcome association period manipulations, inhibition of dSPNs and iSPNs during the delay period robustly biased future choice in opposing directions. Indeed, dSPN inhibition caused animals to switch toward the unrewarded ipsilateral spout (D1;GtACR1: I→C no stim: 3.6±0.6% vs. stim: 6.1±3.8%, p=0.65; C→I no stim: 5.4±0.7% vs. stim: 67.0±11.1%, p=0.01), whereas iSPN inhibition biased switching toward the unrewarded contralateral spout on the next trials (A2a;GtACR1: I→C no stim: 4.0±1.03% vs. stim: 86.0±5.1%, p=0.008; C→I no stim: 9.9±1.8% vs. stim: 5.1±3.8%, p=0.2) (**Figure 7D-E; Supp Figure 8B, D**). Reaction times on trial n+1 were not affected by ENL-period inhibition (**Supp Figure 8C, E**), and no changes in action selection or reaction times were observed in Cre-negative floxed-GtACR1 control animals (**Supp Figures 8F-G; Table S1**). Thus, striatal signals are actively maintained across the delay period and required for proper future action selection. This indicates that inhibition of striatal activity in a period free of task-related movements and rewards disrupts future action selection, consistent with a requirement for striatal activity during this interval to maintain the motor planning memory. Together, these results demonstrate that striatal projection neurons maintain a short-term memory of recent action-outcome associations across the delay period, which is required to guide subsequent choice.

**Figure 7.**
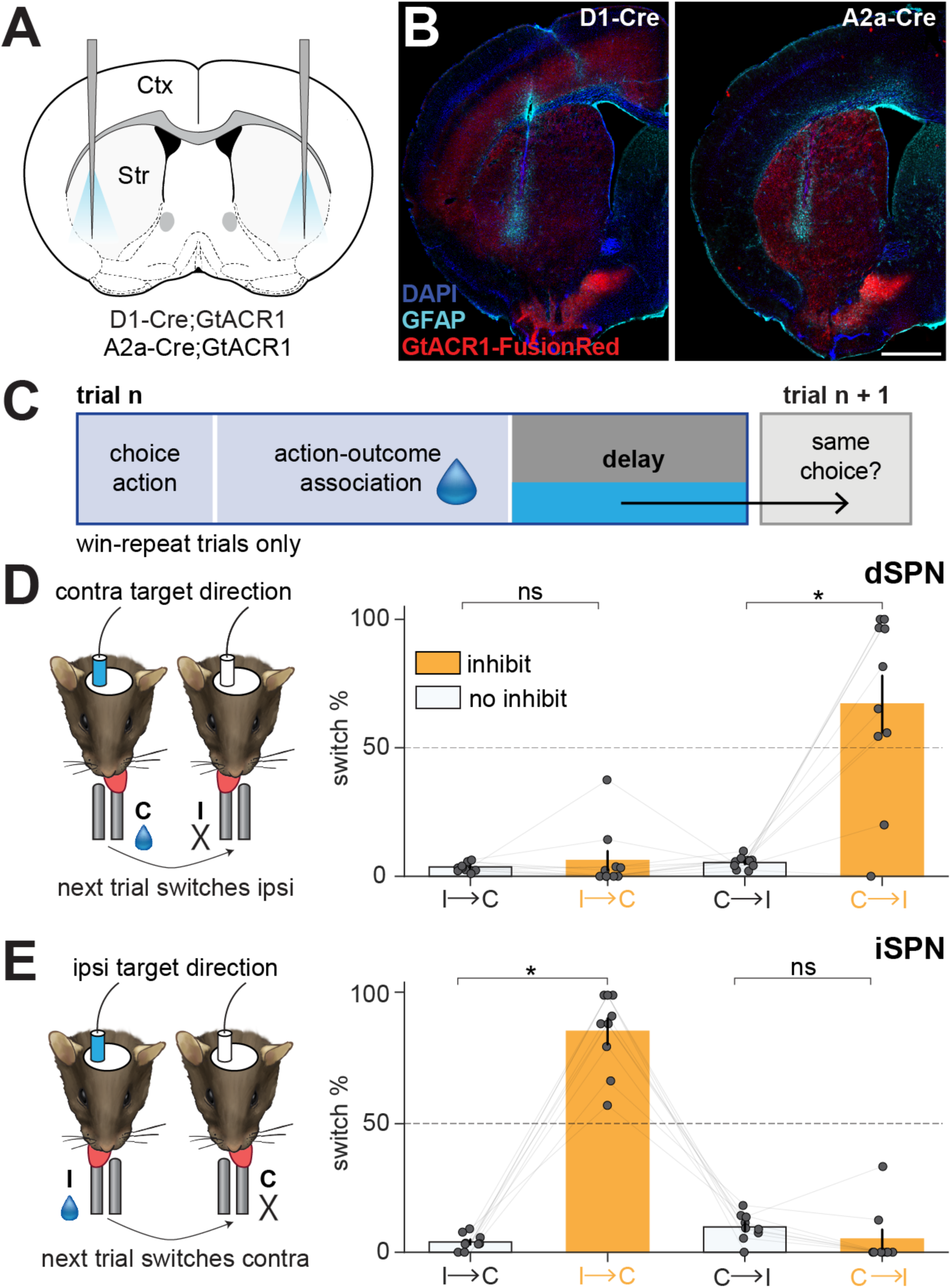
Cell-type specific inhibition of dSPNs and iSPNs during the delay period bidirectionally modulate action updating. **A.** Coronal schematic showing tapered fiber implantation in the ventrolateral striatum. **B.** Representative low-magnification histology showing GtACR1-FusionRed (red), GFAP (cyan), and DAPI fluorescence (blue) in dSPNs and iSPNs. Scale bar: 1 mm. **C.** Schematic of the task structure showing the period of optogenetic inhibition. Inhibition was delivered during the delay period following rewarded trials (continuous inhibition, 0.5 mW, 1.5 seconds, 488 nm blue light). **D.** Schematic (*left*) and summary (*right*) of the effects that dSPN inhibition during the delay on trial n has on the action selected in trial n+1. dSPN inhibition promotes choice of the spout ipsilateral to the stimulated site such that it greatly increases the percent of contralateral to ipsilateral spout (C→I) reward-switch trials. Orange indicates inhibition trials; white indicates no stimulation trials. **E.** Same as in D, except for iSPNs. iSPN inhibition promotes choice of the spout contralateral to the stimulated site such that it greatly increases the fraction of ipsilateral to contralateral spout (I→C) reward-switch trials. **D-E.** N=7 mice, n=28 sessions (D1;GtACR1); N=6 mice, n=18 sessions (A2a;GtACR1). Each dot represents one hemisphere per mouse. Bar plots represent mean across mice ± SEM. For all panels: ns, not significant; ∗*p* < 0.05.

## DISCUSSION

Here, using a memory-guided decision-making task in which mice use recent reward history to decide whether to repeat or switch their upcoming choices, we show that VLS SPN activity and DA fluctuations link action-outcomes to future motor outputs. VLS DA signals are modulated by reward receipt, reward omission, and recent outcome history. dSPN and iSPN calcium activity reflect recent action-outcome history and predict whether animals will switch or repeat on the next trial. Causal manipulations further demonstrate that this activity is necessary for outcome-guided choice: gain- and loss-of-function perturbations of dSPNs and iSPNs during either the action-outcome association period or the delay period bias future choices away from the action favored by recent reward history. Together, our findings indicate that striatal SPN activity maintains an updated action plan across the inter-action interval, providing a short-term action-outcome memory that links a completed action and its outcome to the next choice.

### Striatal activity links recent outcomes to future action selection

Prior work attributes striatal activity to a variety of functions, including movement, reinforcement and selection of actions, value-based decision-making, vigor, and kinematics (Kravitz et al., 2010; Bateup et al., 2010; Kravitz et al., 2012; Tai et al., 2012; Cui et al., 2013; Yttri and Dudman, 2016; Akhlaghpour et al., 2016; Donahue, 2018; Dhawale et al., 2021; Bolkan et al., 2022; Mizes et al., 2023; van Beest et al., 2024; Hardcastle et al., 2025; Reinhold et al., 2025). This diversity has made it difficult to determine whether striatal activity reflects ongoing movement, reward processing, or contributes to the decision process itself. Our task helps separate these possibilities because the action-outcome association period is temporally separated from the next choice. Reward receipt or omission provides the information necessary to determine the optimal next action, but the animal expresses that choice only after a delay.

Several findings indicate that VLS SPN activity is necessary for the maintenance of the memory of the optimal action in this interval. First, spontaneous licks during the delay period are aligned with the direction of the upcoming choice, suggesting that the future motor plan begins to form before the go cue. Second, dSPN and iSPN activity during the outcome period predict whether animals will switch or repeat on the next trial. Finally, optogenetic perturbations during either the action-outcome association period or the delay period alter the subsequent choice. Together, these findings suggest that VLS activity does not simply encode reward consumption or ongoing licking. Instead, this activity helps maintain or bias the future motor plan selected on the basis of recent action-outcome history.

### Direct and indirect pathway balance controls future motor plans

Classical models proposed that dSPNs promote movement whereas iSPNs suppress movement (Alexander et al., 1986; Albin et al., 1989; DeLong, 1990). However, this view has been largely revised by studies, including the results observed here, showing that dSPNs and iSPNs are coactive during movement and can display action-specific activity (Cui et al., 2013; Isomura et al., 2013; Barbera et al., 2016; Markowitz et al., 2018; Parker et al., 2018). Current models, including the competitive and refinement model, propose that dSPN and iSPN activity operates within action-related ensembles to shape the selection, refinement, and execution of specific motor programs (Bariselli et al., 2019; Lee and Sabatini, 2025). In the competitive model, the balance between dSPN and iSPN activity determines the probability, duration, or vigor of a motor program (Bariselli et al., 2019). Whereas, in the refinement model, iSPNs help sharpen or constrain selected actions rather than broadly suppressing all competing actions (Lee and Sabatini, 2025).

Our results fit within this updated framework. We find that both dSPNs and iSPNs are active during the task, carry recent action-outcome information, and predict future choice. However, as previously reported, pathway-specific perturbations produce opposing directional biases in the animal’s next action (Tai et al., 2012; Tecuapetla et al., 2016; Wang et al., 2018; Geddes et al., 2018; Lee et al., 2020; Lee and Sabatini, 2021). These findings suggest that the relevant computation is not whether to move or withhold movement, but how imbalanced activity between dSPN and iSPNs biases the animal toward one lateralized action plan over another. Rewarded outcomes may stabilize the recently selected action plan, favoring repetition, whereas reward omission may shift the circuit away from the suboptimal action and toward the alternative choice. Thus, dSPNs and iSPNs may jointly maintain an action-specific motor-plan state, with perturbations shifting the balance of that state and causing animals to execute choices that conflict with recent reward history.

### A circuit and cellular substrate for action-outcome memory

Our data support a role for VLS in maintaining a short-term action-outcome-dependent memory of the motor plan, but they do not require that this memory is stored in the striatum alone. A more likely model is that this information is maintained within interconnected cortico-basal ganglia-thalamo-cortical loops (Alexander et al., 1986; Hunnicutt et al., 2016; Lee et al., 2020). Frontal motor regions such as ALM can maintain preparatory activity before action selection (Li et al., 2015; Chen et al., 2017; Guo et al., 2017; Svoboda and Li, 2018), and VLS activity may update, gate, or bias this cortical motor-plan state according to recent reward history. In this framework, the striatum is not simply a downstream motor structure or a site of gradual reinforcement learning. Rather, it participates in a recurrent circuit that updates future actions on the timescale of individual trials.

However, the form of this short-term action-outcome memory remains unresolved. Persistent activity across cortical, striatal, thalamic, and downstream motor circuits could maintain the upcoming action plan electrically (Wang et al., 2021). Alternatively, DA- and calcium-dependent intracellular signaling could maintain a biochemical trace of the recent action and outcome. DA is well positioned to provide this updating signal because VLS DA reflects reward receipt, reward omission, and recent outcome history (Bromberg-Martin et al., 2010). Midbrain DA acts on D1 receptors in dSPNs and D2 receptors in iSPNs (Gerfen and Surmeier, 2011), which differentially regulate cAMP/PKA signaling and thereby influence excitability, synaptic integration, and plasticity (Tritsch and Sabatini, 2012). Recent work suggests that PKA signaling in dSPNs and iSPNs can be differentially sensitive to DA increases and decreases (Lee et al., 2021), raising the possibility that pathway-specific biochemical signaling could contribute to the maintenance of action-outcome information across the inter-trial interval. Consistent with this idea, dSPN and iSPN calcium dynamics in our task do not simply mirror DA transients, suggesting that DA signals are transformed into pathway-specific representations of outcome history and future action bias. Thus, the relevant memory trace may reflect persistent circuit activity, a DA-dependent biochemical state, or an interaction between the two.

### Caveats and limitations

Fiber photometry reports population-level calcium dynamics and cannot determine how individual SPNs encode action direction, outcome, history, or future choice. This limitation is important in the striatum, where heterogeneous neural signals may be obscured in bulk fluorescence measurements (Legaria et al., 2022). In addition, striatal photometry signals may not directly correspond to somatic spiking. The persistent GCaMP signals observed here could reflect action potentials, subthreshold synaptic input, dendritic or neuropil signals, intracellular biochemical signaling, or some combination of these processes.

### Conclusions

Together, our findings suggest that dSPN and iSPN activity in the VLS links recent action-outcome history to future action selection. This expands the role of the striatum beyond reinforcement learning and motor control to include the short-term maintenance and updating of action-outcome information. Such a mechanism may be particularly relevant to disorders involving basal BG dysfunction, including Parkinson’s disease, obsessive-compulsive disorder, Tourette’s syndrome, and addiction, in which action selection, behavioral flexibility, and outcome-guided updating are disrupted.

## Supplemental Figures

**Supplemental Figure 1.**
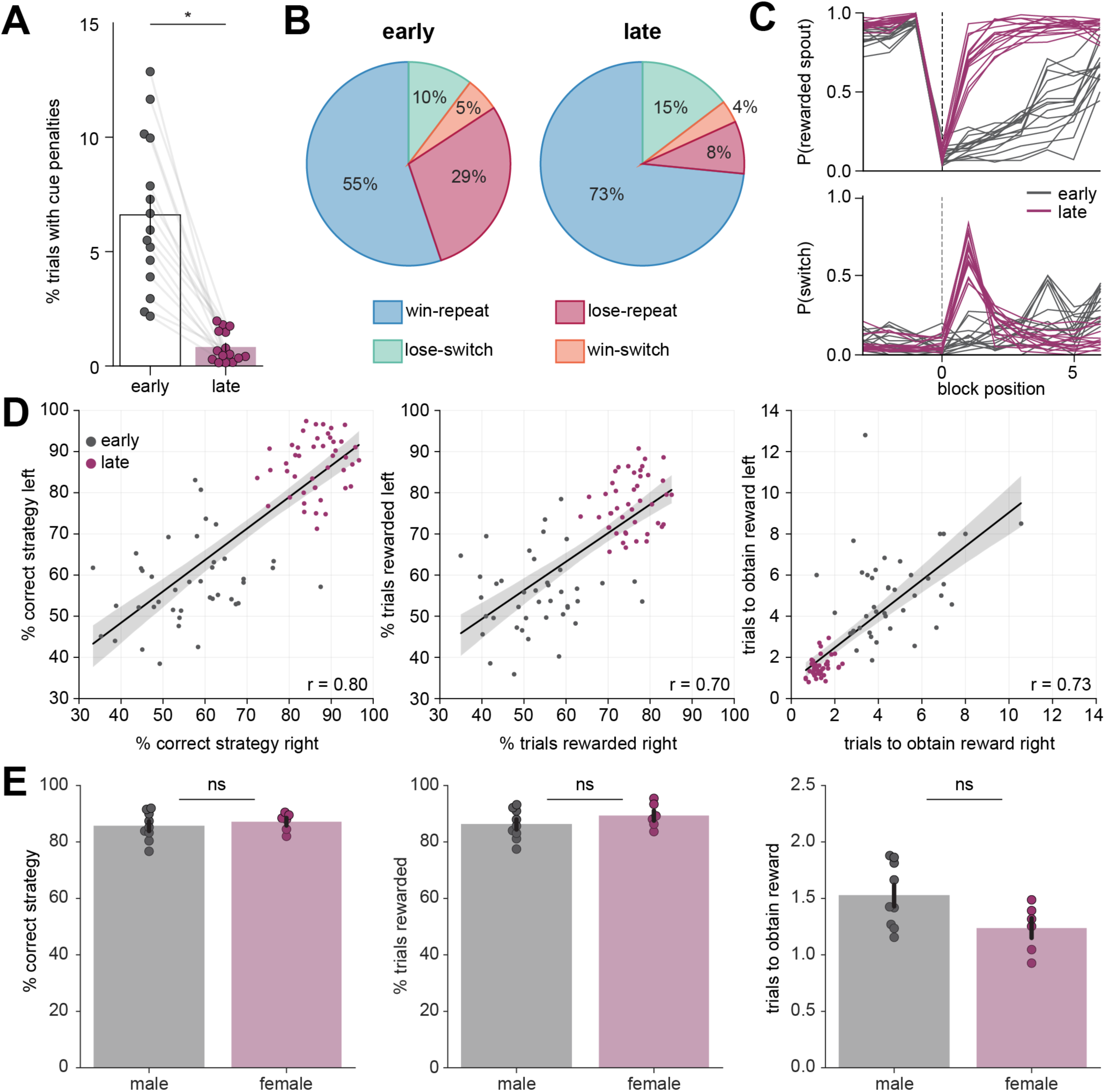
A head-fixed dynamic foraging task for memory guided decision making. **A.** Percentage of trials with lick penalties in the cue period early and late in training. Each circle represents one mouse (an average of 2-3 sessions early and 3 sessions late in training per mouse). **B.** Pie charts showing the proportion of trials corresponding to each behavioral strategy (win-switch, win-repeat, lose-repeat, and lose-switch) early and late in training. Percentages represent averages across sessions. **C.** Probability of selecting the rewarded spout (*top*) and the probability of switching between spouts on the current trial relative to the last trial (*bottom*) as a function of trial position around block transitions (x=0). Solid lines represent average probabilities of individual mice early (grey) and late (purple) in training. Each line represents one mouse (an average of 2-3 sessions early and 3 sessions late in training per mouse). **D.** Comparison of the percentage of trials in which the animal used the correct task strategy (*left*), percentage of rewarded trials (*middle*), and number of trials to first reward after a block switch (*right*) for left and right choice blocks. Each dot represents an individual session early (grey) or late (purple) in training. Linear regression fits to the data are overlaid; gray shading indicates a 95% confidence interval. **E.** Comparison of the percentage of trials in which animals used the correct task strategy (*left*), percentage of rewarded trials (*middle*), and the number of trials to first reward after a block switch (*right*) for male (N=9) and female (N=6) mice late in training. **A-E**. N=15 mice, n=42 sessions early in training, n=45 sessions late in training. Bar plots represent mean across mice ± SEM. For all panels: ns, not significant; ∗*p* < 0.05.

**Supplemental Figure 2.**
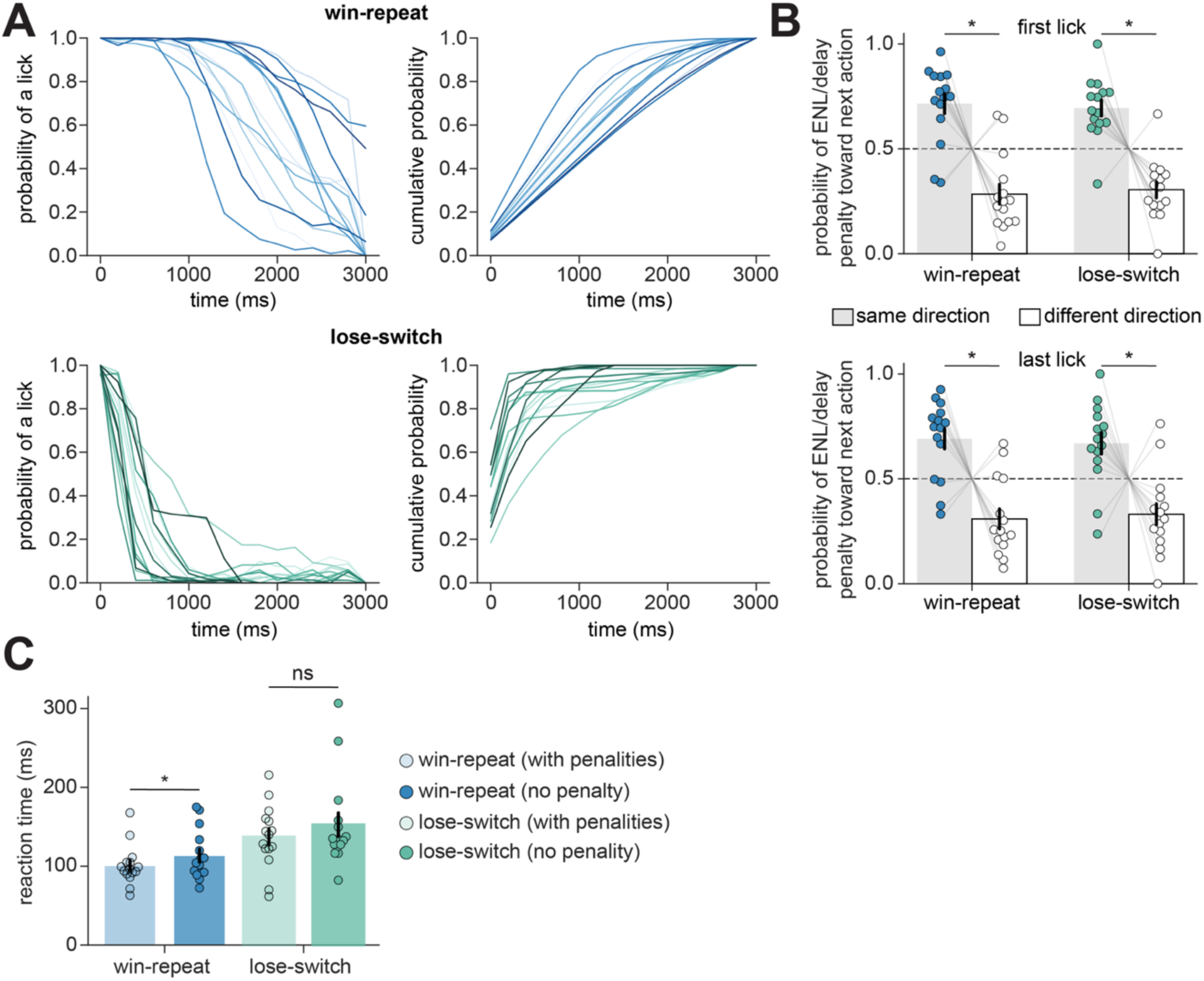
Choice is guided by the previous action-outcome association. **A.** Probability of licking (*left*) and cumulative probability of licking (*right*) across the action-outcome association period in 200 ms bins for win-repeat (*top*, blue) and lose-switch (*bottom,* green) trials for individual mice. t=0 represents the choice lick. Each line represents 1 mouse (average of 3 well trained sessions per mouse). **B.** On trials with lick penalties during the delay period, the probability that the first (*top*) or last (*bottom*) lick penalty occurs in the direction of the next choice action (grey) or not (white) is shown for win-repeat and lose-switch trials. Each circle represents one mouse (average of 3 well trained sessions per mouse). **C.** Average response times for win-repeat and lose-switch trials, separated by whether mice withheld licking during the delay period (darker colors) or made a penalty lick during the delay period (lighter colors). Each circle represents the average reaction time, with and without penalties, per mouse. **A-C**. N=15 mice, n=45 sessions late in training. Bar plots represent mean across mice ± SEM. Each circle represents one mouse. For all panels: ns, not significant; ∗*p* < 0.05.

**Supplemental Figure 3.**
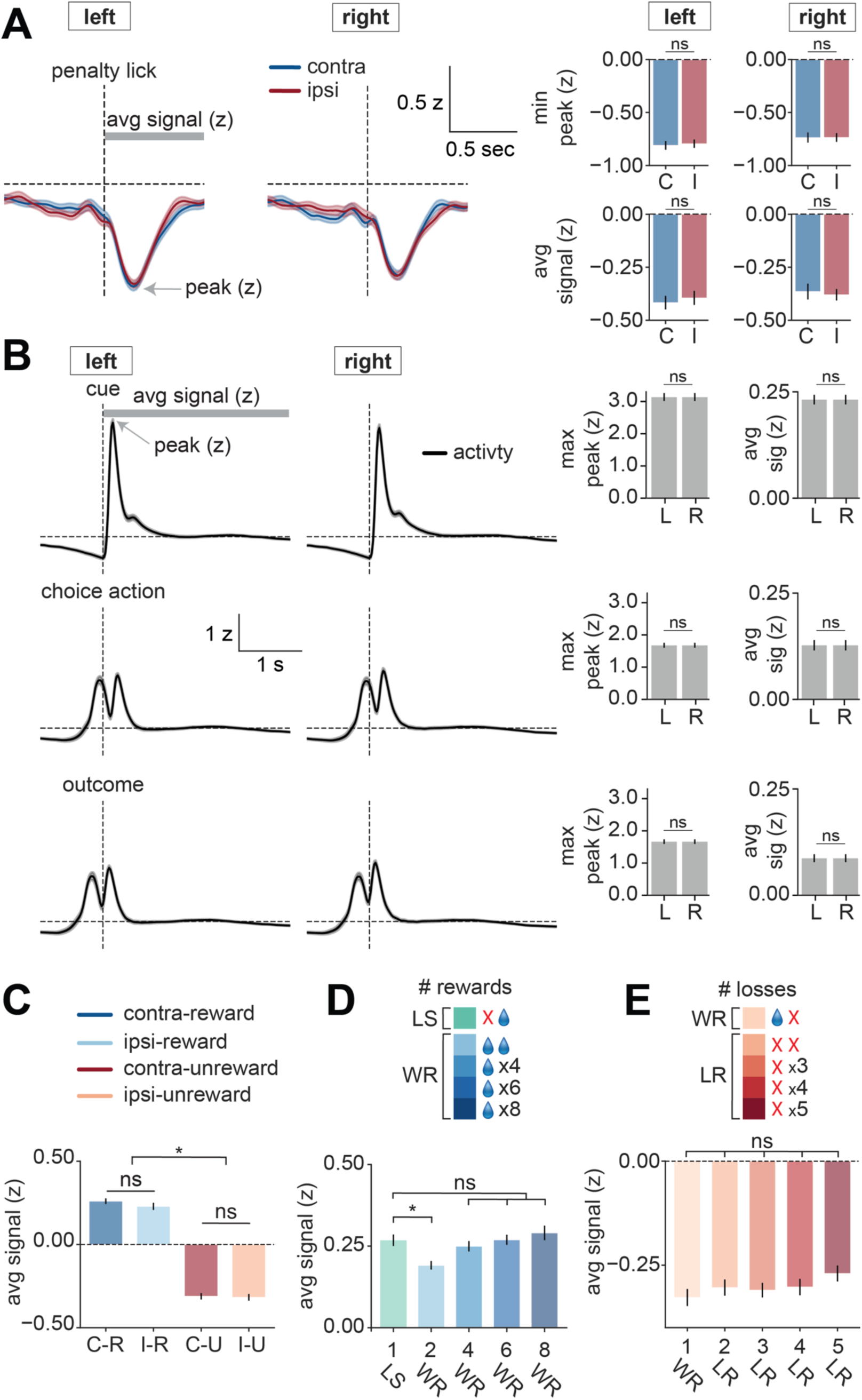
Dopamine in the ventrolateral striatum encodes reward and action history. **A.** DA signals aligned to spontaneous penalty licks during the delay period. *left*: Activity during contralateral (C, blue) and ipsilateral (I, red) penalty licks recorded from the left and right hemispheres. Thick grey line denotes quantification area of the average z-score across time (“avg signal”); arrow denotes maximum or minimum peak z-score. *right*: Quantification (*top*: minimum peak z-score; *bottom*: average z-score signal) of the response magnitudes for contralateral (blue) and ipsilateral (red) penalty licks recorded from the left and right hemispheres. **B.** DA dynamics aligned to task-specific events. *left*: Activity aligned to the cue (*top*), choice action lick (*middle*), and outcome lick (*bottom*) for the left and right hemispheres. *right*: Quantification (*left*: maximum peak z-score; right: average z-score signal) of the response magnitudes across the cue, choice action, and the outcome period for the left and right hemisphere. **C** Quantifications of the average signal of the response magnitudes contralateral (C) vs. ipsilateral (I) and rewarded (R) vs. unrewarded (U) trial types. **D.** Quantifications of the average z-scored signal of consecutive rewarded choice actions. WR: win-repeat, LS: lose-switch trials. **E.** Quantifications of the average z-scored signal of consecutive unrewarded choice actions. WR: win-repeat, LR: lose-repeat trials. **A-E**. N=17 mice (6 D1-Cre and 11 A2a-Cre), n=110 sessions (60,276 trials). Signals are z-scored, aligned to event of interest, and averaged across mice. Shaded bars around mean signal represents SEM. Bar plots represent mean across mice ± SEM. For all panels: ns, not significant; ∗*p* < 0.05.

**Supplemental Figure 4.**
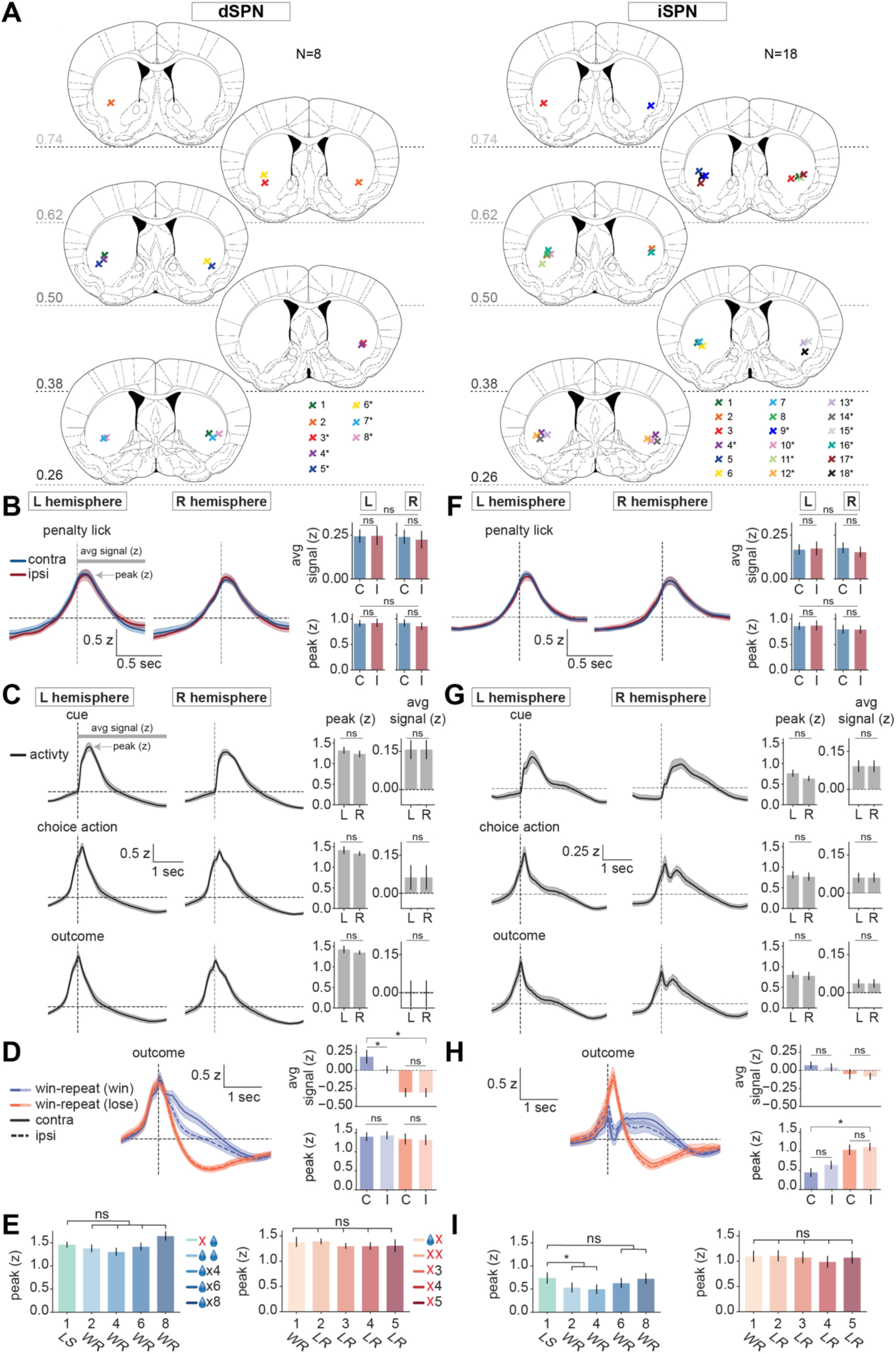
dSPN and iSPN calcium activity encodes action-specific outcome value and reward history. **A.** Summary of tip locations of fiber placements for dSPN (N=8, *left*) and iSPN recordings (N=18, *right*). Coronal sections are shown at the indicated anterior-posterior coordinates (mm from bregma). Colored symbols denote recording sites from hemispheres in individual animals. Asterisks (*) indicate animals that we co-injected with AAV9-hSyn-rDAm. **B.** Calcium activity in dSPNs aligned to spontaneous penalty licks during the delay period. *left*: Activity during contralateral (C, blue) and ipsilateral (I, red) penalty licks recorded from the left (L) and right (R) hemispheres. Grey line denotes quantification area of the average z-score across time (“avg signal”); arrow denotes peak z-score. *right*: Quantification (*top*: average z-scored signal; *bottom*: peak z-score) of the response magnitudes for contralateral (blue) and ipsilateral (red) penalty licks recorded from the left (L) and right (R) hemispheres. **C.** Calcium activity in dSPNs aligned to task-specific events. *left*: Activity aligned to the cue (*top*), choice action lick (*middle*), and outcome period (*bottom*) for the left and right hemispheres. *right*: Quantification (*left*: peak z-score; right: average z-scored signal) of the response magnitudes across the cue, choice action, and the outcome period for the left (L) and right hemisphere (R). **D.** *left*: Calcium activity in dSPNs aligned to the outcome lick on trials in which the same spout was selected on consecutive trials following a rewarded trial. Trials are segregated by current outcome (win, purple; loss, orange). Traces are shown for contralateral (solid) and ipsilateral (dashed) choice actions. *right:* Quantifications (*top*: average z-scored signal; *bottom*: peak z-score) of the response magnitudes for contralateral (C) vs. ipsilateral (I) and win (purple) vs. lose (orange) trial types. **E.** Quantifications of the peak z-score of consecutive rewarded (*left*) or unrewarded (*right*) choice actions. WR: win-repeat, LS: lose-switch, and LR: lose-repeat trials. **F-I.** As in B-E, except for iSPN photometry recordings. **A-I**. N=8 D1-Cre mice, n=65 sessions (35,538 trials). N=18 A2a-Cre mice, n=120 sessions (65,846 trials). Signals are z-scored, aligned to event of interest, and averaged across mice. Shaded bars around mean signal represents SEM. Bar plots represent mean across mice ± SEM. For all panels: ns, not significant; ∗*p* < 0.05.

**Supplemental Figure 5.**
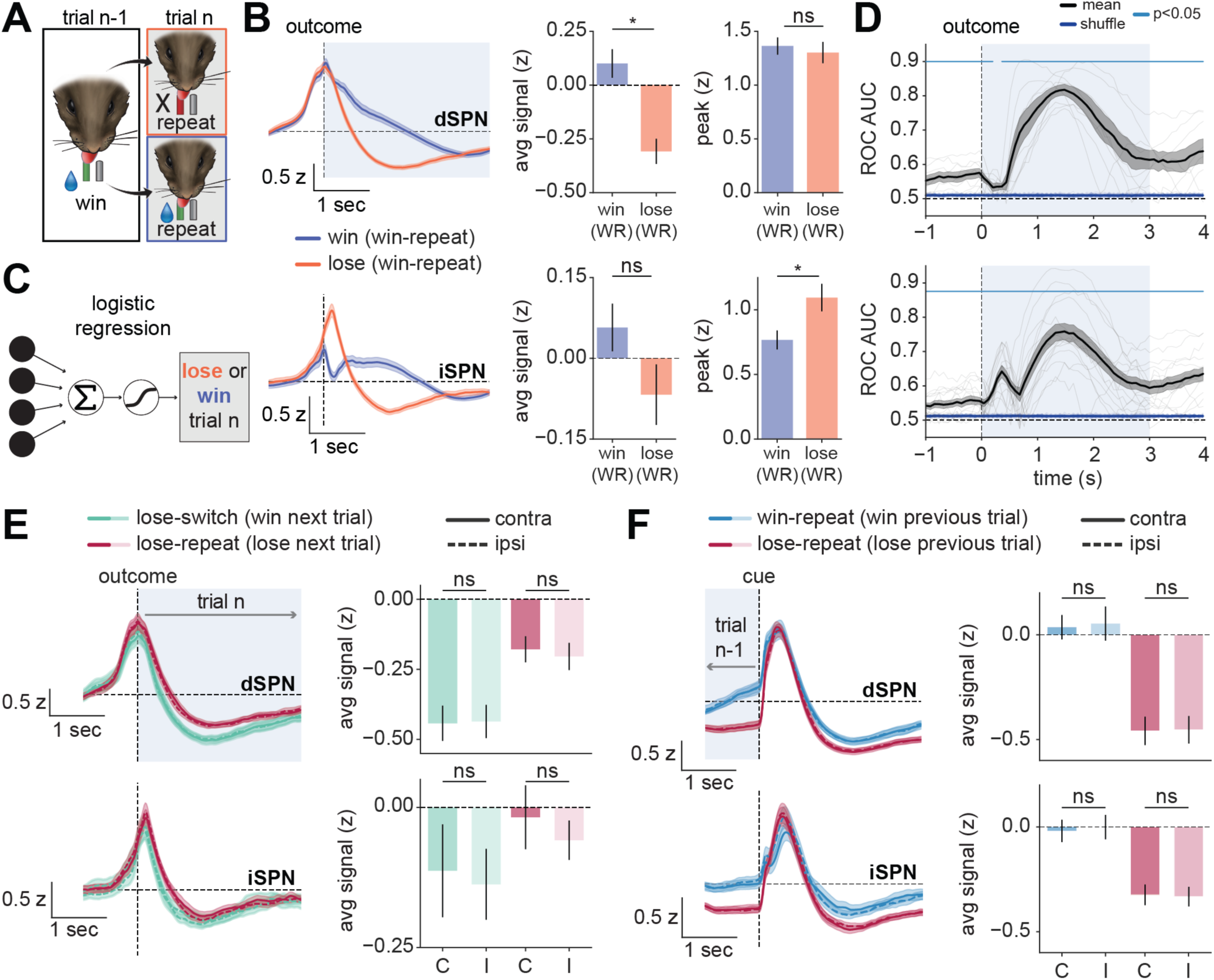
Future action and recent outcome history can be decoded from dSPN and iSPN population activity. **A.** Behavioral schematic illustrating rewarded and unrewarded outcomes after matched reward and action history. On trial n-1, mice selected the rewarded spout and received reward. On trial n, mice repeated the same action and either received reward again, classified as a win-repeat, or failed to receive reward, classified as a loss after a win-repeat sequence. **B.** *left*: Calcium activity in dSPNs (*top*) or iSPNs (*bottom*) aligned to the outcome lick on trial n, segregated by trial outcome (win/reward, purple; lose/unreward, orange). Shaded region denotes analysis window to predict a win or lose on trial n. *right:* Quantification of the average z-scored and peak signal (trial n window) for win (win-WR) and lose (lose-WR) conditions. **C.** Schematic of a logistic regression classifier trained to predict current-trial outcome from striatal activity on trial n (win or loss on trial n). To control for trial history, the classifier compared trials with the same preceding win-repeat history that differed in the outcome received on trial n. **D.** Time-resolved ROC AUC analysis quantifying decoding of current trial outcome (win vs. lose) from calcium in dSPNs (*top*) or iSPNs (*bottom*). Thick black line: mean ROC AUC; thin black lines: individual mice; thick blue line: mean shuffle, thin blue lines: individual shuffles. Shaded region indicates analysis window. Significant time points are indicated (cyan line, *p* < 0.05, see Methods). **E.** *left*: Calcium activity in dSPNs (*top*) or iSPNs (*bottom*) aligned to the outcome lick on trial n, segregated by next-trial behavior (switch, green; repeat, magenta) for ipsilateral and contralateral choices. Shaded region denotes the quantification window. *right:* Quantification of the average z-scored signal (trial n window) for lose-switch (win next trial) and lose-repeat (lose next trial) trials for ipsilateral (I) and contralateral (C) choices. **F.** *left*: Calcium activity in dSPNs (*top*) or iSPNs (*bottom*) aligned to the cue on trial n, segregated by previous trial outcome (win-repeat, blue; lose-repeat, magenta) for ipsilateral vs. contralateral choices. Shaded region denotes the quantification window. *right:* Quantification of the average z-scored signal (trial n-1) for win-repeat (win previous trial) and lose-repeat (lose previous trial) trials for ipsilateral (I) and contralateral (C) choices. **A-F.** N=8 D1-Cre mice, n=65 sessions (35,538 trials). N=18 A2a-Cre mice, n=120 sessions (65,846 trials). Signals are z-scored, aligned to event of interest, and averaged across mice. Shaded bars around mean signal represents SEM. Bar plots represent mean across mice ± SEM. For all panels: ns, not significant; ∗*p* < 0.05. Abbreviations: ROC AUC: (receiver operating characteristic area under the curve).

**Supplemental Figure 6.**
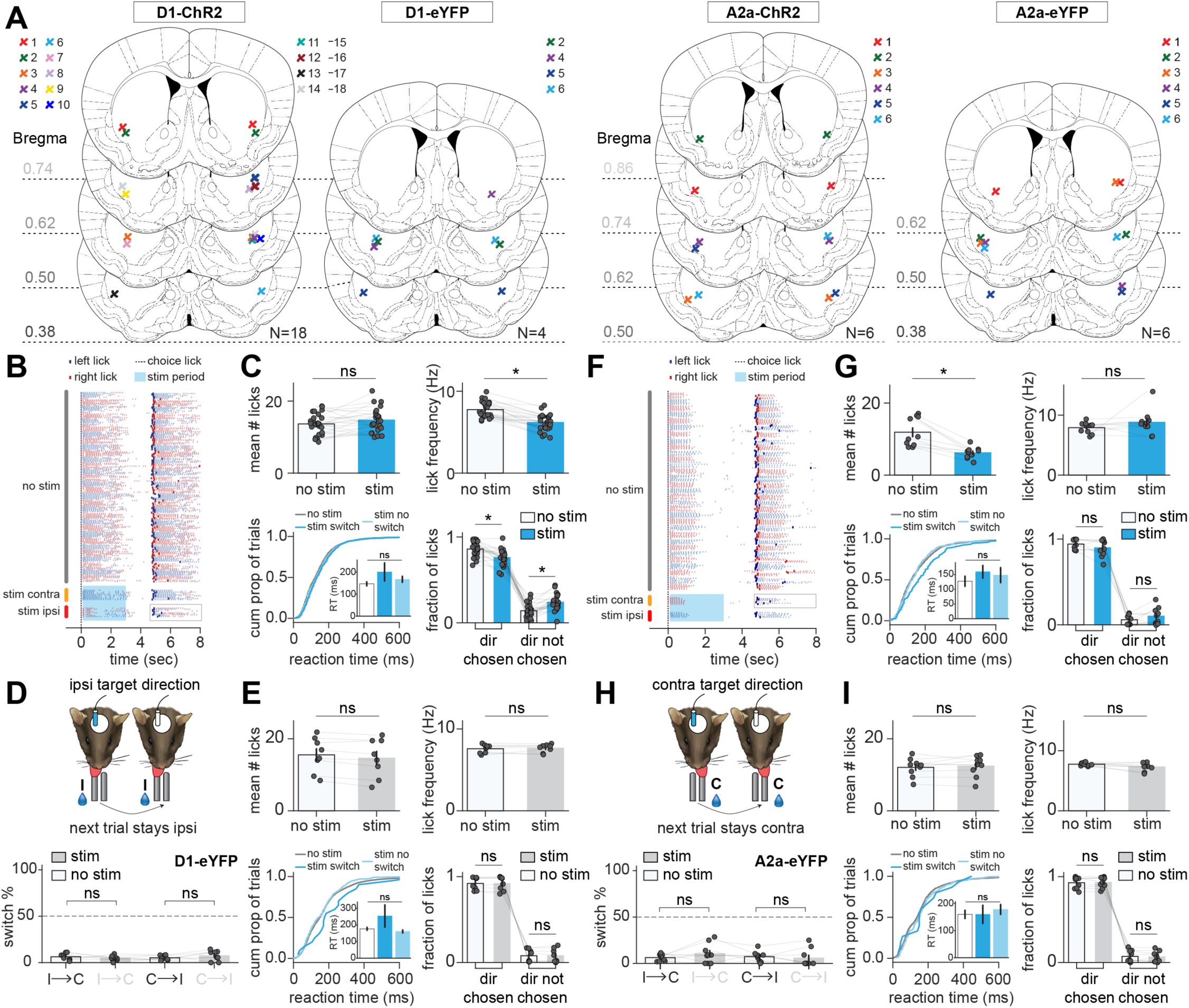
Cell-type specific activation of dSPNs and iSPNs during the action-outcome association period bidirectionally modulate action updating. **A.** Summary of tip locations of fiber placements for dSPNs and iSPNs expressing ChR2 or eYFP controls. Coronal sections are shown at the indicated anterior-posterior coordinates (mm from bregma). Colored symbols denote recording sites from hemispheres in individual animals. Note: precise fiber tips could not be located in 4 D1-ChR2 animals (15-18). **B.** Example licking rasters aligned to the choice lick without stimulation (no stim) and with stimulation of dSPNs during the action-outcome association period. Stimulated trials are grouped by whether stimulation was delivered contralateral (stim contra) or ipsilateral (stim ipsi) to the choice lick direction. Blue ticks indicate left licks; red ticks indicate right licks; shaded region indicates the stimulation period. **C.** Effects of dSPN stimulation on lick output (trial n) and reaction time (trial n+1). *top*: Mean number of licks and lick frequency (Hz) with and without dSPN stimulation. *bottom left*: Cumulative distribution of reaction times for no-stimulation trials (no stim), stimulation trials that resulted in a switch (stim switch), and stimulation trials that did not result in a switch (stim no switch) on trial n+1. Each circle represents the average reaction time per trial across all sessions. Inset: bar plot showing the average response time per condition. *bottom right*: Fraction of licks during the action-outcome association period directed toward the chosen action on trial n (dir chosen) vs. the non-chosen action on trial n (dir not chosen), with and without dSPN stimulation. **D.** Schematic (*top*) and summary (*bottom*) of the effects that control dSPN-eYFP activation during rewarded trial n has on the action selected in trial n+1. Gray indicates stimulation trials; white indicates no stimulation trials. **E.** Same in C, except for stimulation of dSPNs expressing control eYFP fluorophore. **F-G**. Same as in B-C, except for stimulation of iSPNs expressing ChR2. **H.** Same as in D, except for stimulation of iSPNs expressing control eYFP fluorophore. **I**. Same as in C, except for stimulation of iSPNs expressing control eYFP fluorophore. **A-I.** N=18 mice, n=75 sessions (D1-ChR2); N=4 mice, n=21 sessions (D1-eYFP); N=6 mice, n=32 sessions (A2a-ChR1); N=6 mice, n=22 sessions (A2a-eYFP). Each dot represents one hemisphere per mouse. Bar plots represent mean across mice ± SEM. For all panels: ns, not significant; ∗*p* < 0.05.

**Supplemental Figure 7.**
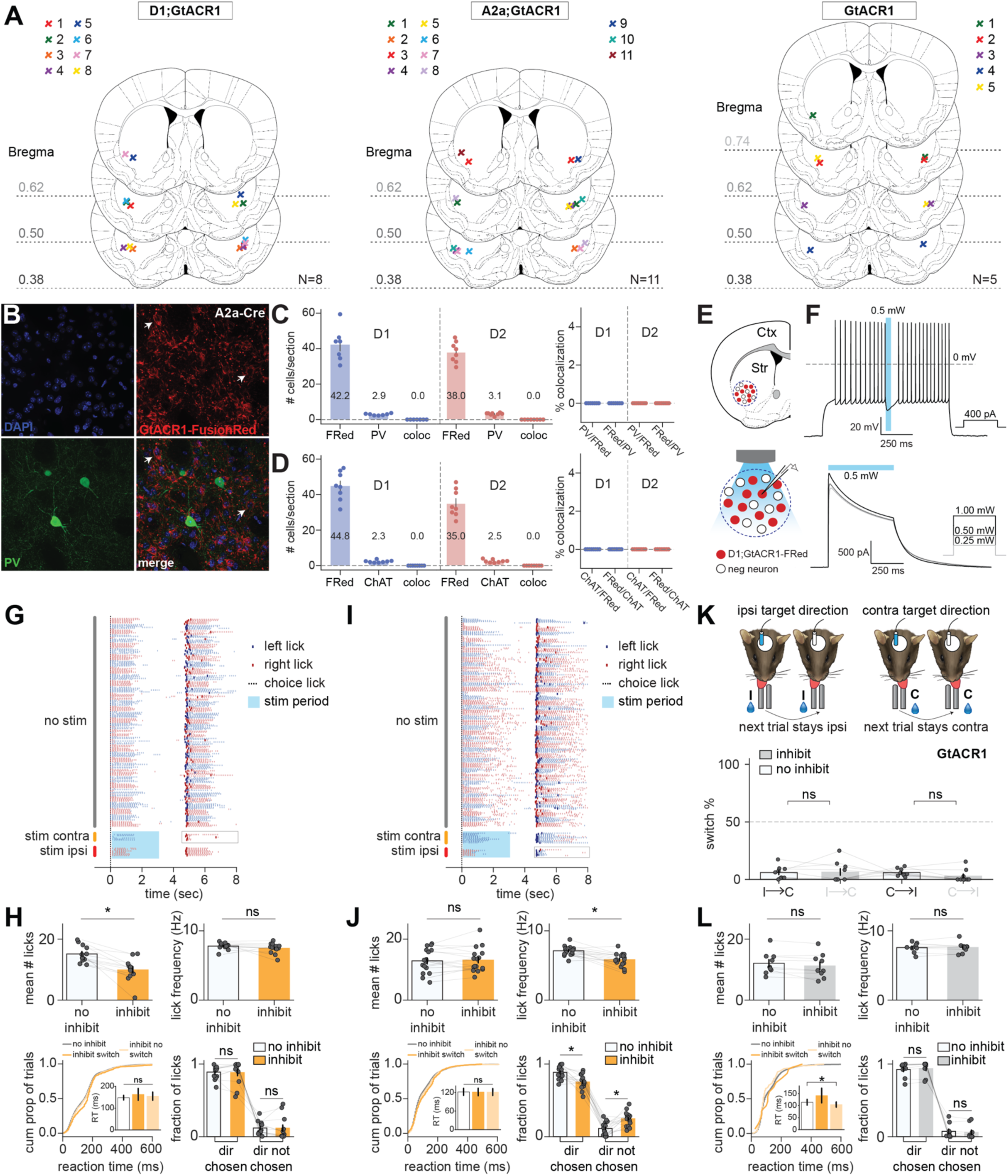
Cell-type specific inhibition of dSPNs and iSPNs during the action-outcome association period bidirectionally modulate action updating. **A.** Summary of tip locations of fiber placements for dSPNs or iSPNs expressing soma localized GtACR1 and floxed GtACR1 control animals. Coronal sections are shown at the indicated anterior-posterior coordinates (mm from bregma). Colored symbols denote recording sites from hemispheres in individual animals. **B.** Representative 40x confocal images from the striatum of A2a;GtACR1-FusionRed mouse immunostained for parvalbumin (PV), showing no overlap between FusionRed and PV positive cells. Arrowheads denote of A2a;GtACR1-FusionRed positive cells are not colocalized with PV positive cells. **C.** *left:* Average number of FusionRed-positive (FRed), PV-positive, and colocalized cells per section in dSPNs (*left*, blue) or iSPNs (*right,* red). *right*: Percent colocalization of PV with FRed (PV/FRed) and FRed with PV (FRed/PV) in dSPNs (*left*, blue) or iSPNs (*right*, red). Each dot represents a unique striatal section per mouse, D1/PV: N=3 mice, D2/PV: N=3 mice. **D.** Same as in B, except for ChAT immunostaining. Each dot represents a unique striatal section per mouse, D1/ChAT: N=3 mice, D2/ChAT: N=3 mice. E. Schematic of whole-cell recording configuration for *in vitro* slice validation of GtACR1. **F**. *top*: Representative whole-cell current-clamp recording of a GtACR1 expressing dSPN illuminated with 488 nm light (0.5 mW) during a 400 pA current injection to inhibit spiking (N=3 mice, n=8 cells,). *bottom*: Representative whole-cell voltage-clamp photocurrent recording of a GtACR1 expressing dSPN illuminated (488 nm) with increasing light power for 500 ms (N=3 mice, n=9 cells). Blue bar represents the period of inhibition. **G.** Example licking rasters aligned to the choice lick without inhibition (no stim) and with inhibition of dSPNs during the action-outcome association period. Trials are grouped by whether inhibition was delivered contralateral (stim contra) or ipsilateral (stim ipsi) to the choice lick direction. Blue ticks indicate left licks; red ticks indicate right licks; shaded region indicates the inhibition period. **H.** Effects of dSPN inhibition on lick output (trial n) and reaction time (trial n+1). *top*: Mean number of licks and lick frequency (Hz) with and without dSPN inhibition. *bottom left*: Cumulative distribution of reaction times for no-inhibition trials (no inhibit), inhibition trials that resulted in a switch (inhibit switch), and inhibition trials that did not result in a switch (inhibit no switch) on trial n+1. Each circle represents the average reaction time per trial across all sessions. Inset: bar plot showing the average response time per condition. *bottom right*: Fraction of licks during the action-outcome association period directed toward the chosen action on trial n (dir chosen) versus the non-chosen action on trial n (dir not chosen), with and without dSPN inhibition. **I-J**. Same as in B-C, except for iSPNs expressing GtACR1. **K.** Schematic (*top*) and summary (*bottom*) of the effects that blue light activation during rewarded trial n has on the action selected in trial n+1 in GtACR1 control animals. Gray indicates stimulation trials; white indicates no inhibition trials. **L**. Same as in C, except for Cre-negative floxed-GtACR1 control animals. **G-L.** N=8 mice, n=27 sessions (D1;GtACR1), N=11 mice, n=24 sessions (A2a;GtACR1), N=5 mice, n=18 sessions (Cre-negative floxed-GtACR1). Each dot represents one hemisphere per mouse. Bar plots represent mean across mice ± SEM. For all panels: ns, not significant; ∗*p* < 0.05

**Supplemental Figure 8.**
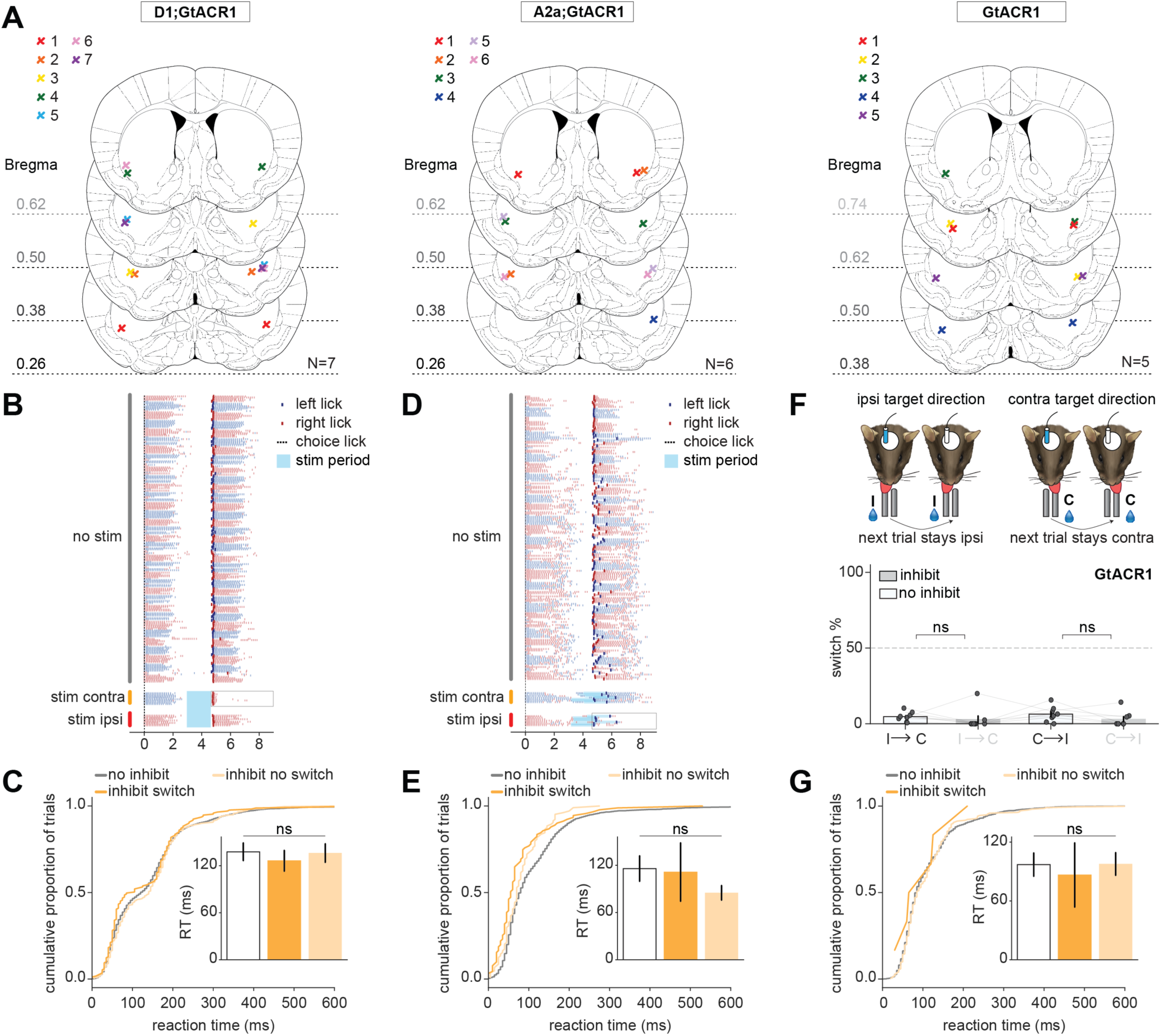
Cell-type specific inhibition of dSPNs and iSPNs during the delay period bidirectionally modulate action updating. **A**. Summary of tip locations of fiber placements for dSPNs or iSPNs expressing GtACR1 and GtACR1 control animals. Coronal sections are shown at the indicated anterior-posterior coordinates (mm from bregma). Colored symbols denote recording sites from hemispheres in individual animals. **B**. Example licking rasters aligned to the choice lick without inhibition (no stim) and with inhibition of dSPNs during the delay/ENL period. Trials are grouped by whether inhibition was delivered contralateral (stim contra) or ipsilateral (stim ipsi) to the choice lick direction. Blue ticks indicate left licks; red ticks indicate right licks; shaded region indicates the inhibition period. **C**. Effects of dSPN inhibition on the reaction time of trial n+1. Cumulative distribution of reaction times for no-inhibition trials (no inhibit), inhibition trials that resulted in a switch (inhibit switch), and inhibition trials that did not result in a switch (inhibit no switch) on trial n+1. Each circle represents the average reaction time per trial across all sessions. Inset: bar plot showing the average response time per condition. **D-E**. Same as in B-C, except for iSPNs expressing GtACR1. **F.** Schematic (*top*) and summary (*bottom*) of the effects that blue light activation during the delay on trial n has on the action selected in trial n+1 in Cre-negative floxed-GtACR1 control animals. Gray indicates stimulation trials; white indicates no inhibition trials. Each dot represents one hemisphere per mouse. **G.** Same as in C, except for Cre-negative floxed-GtACR1 control animals. **A-G.** N=7 mice, n=28 session (D1;GtACR1); N=6 mice, n=18 sessions (D2;GtACR1), N = 5 mice, n=18 sessions (Cre-negative floxed-GtACR1). Bar plots represent mean across mice ± SEM. For all panels: ns, not significant; ∗*p* < 0.05.

## ACKNOWLEDGMENTS

We thank John Assad, Byungwoo Kang, Tarun Kamath, Shijia Liu, Kevin Mastro, Priya Srikanth, Match McGregor, Alyson Iolin, and all members of the Sabatini lab for helpful conversations, analysis advice, technical assistance, and support. We thank Elizabeth Drewry and CT King for administrative support and mouse colony maintenance. We thank Ofer Mazor and Pavel Gorelik of the Harvard Medical School Research Instrumentation Core and Paul Jonak for their assistance in designing and implementing the memory-guided decision-making task hardware and software. We thank the Core for Imaging Technology & Education (CITE) at Harvard Medical School for assistance on confocal imaging. This work was supported by a NIH BRAIN Initiative Postdoctoral Fellowship (F32MH125596, A.E.G.), BBRF Young Investigator Award (A.E.G.), Harvard Medical School Mahoney Postdoctoral Fellowship (A.E.G.), the Howard Hughes Medical Institute (B.L.S.), the Foundation for OCD Research (B.L.S.), and the NIH (NIH R35NS137336, B.L.S.).

## AUTHOR CONTRIBUTIONS

A.E.G., G.M., and B.L.S. conceptualized the project. A.E.G, G.M., and B.L.S designed all experiments. A.E.G., M.A.A., R.Y.Z., R.N.A., G.M., T.M.H., and M.D.B. collected all the data. W.W. performed slice electrophysiology recordings. A.E.G., L.C.M., and C.C.B. analyzed all the data, with input from B.J.G.V.D.B and G.M. D.R.H. and B.J.G.V.D.B. provided technical assistance, data analysis expertise, and software. A.E.G. and B.L.S. wrote and edited the manuscript.

## METHODS

### RESOURCE AVAILABILITY

#### Lead contact

Further information and requests for resources, reagents, and data should be directed to and will be fulfilled by the lead contact, Bernardo Sabatini.

### EXPERIMENTAL MODEL AND SUBJECT DETAILS

#### Animals

Male and female C57Bl6/J mice (WT, Jackson Labs) aged 8 weeks to 4 months were used. For behavior and photometry experiments, hemizygous *Drdla-Cre* (B6.FVB(Cg)-Tg(Drd1-cre)EY262Gsat/Mmucd, 030989-UCDj) (Gerfen et al., 2013) and *Adora2a-Cre* (B6.FVB(Cg)-Tg(Adora2a-cre)KG139Gsat/Mmucd, 036158-UCD) (Gerfen et al., 2013) of either sex were acquired from MMRRC UC Davis and back crossed to C57Bl6/J for at least 6 generations. For optogenetic experiments, hemizygous *Drdla-Cre* or *Adora2a*-Cre of either sex were crossed to hemizygous *R26-CAG-LNL-GtACR1-ts-FRed-Kv2.1* reporter mouse (Jackson Laboratory, stock #033089) (Li et al., 2019) to yield D1;GtACR1 or A2a;GtACR1 mice, and hemizygous *Adora2a*-Cre mice of either sex were crossed with homozygous Ai32(RCL-ChR2(H134R)/EYFP) (Jackson Laboratory, stock # 024109) to yield A2a;Ai32 mice. All behavioral animals were singly housed on a 12-hour reverse light cycle with free access to food and no access to free water. All behavioral manipulations were performed during the dark light cycle phase. Animal care and experimental manipulations were performed in accordance with protocols approved by the Harvard Standing Committee on Animal Care, following guidelines described in the US NIH Guide for the Care and Use of Laboratory Animals.

### METHOD DETAILS

#### Surgical procedures

Surgical procedures were performed on male and female mice 8-12 weeks of age. Mice were anesthetized with 2-3% isoflurane and 80% oxygen under aseptic conditions using a stereotaxic frame (David Kopf Instruments Model 900). For intracranial injections, 250 µm diameter craniotomies were made with an electric drill (Foredom Electric Company K.1070) with a ball bur (Busch and Co. S33289) attached to the manipulator. All reagents were injected through a pulled glass pipette (Drummond Scientific Company pipettes) with a tip of approximate 50 µm (pulled with a P-97 model Sutter Instrument Co. pipette puller). To avoid leak into other brain regions and back spill through the pipette track the injection pipette was lowered 100 µm ventral to the region of injection before being brought up to the point of injection. The pipette was left in place for 1-3 minutes prior to injection and solution was delivered at a rate of 50-100 nL/min using a UMP3 micro-syringe pump (Micro4, World Precision Instruments). Following injection, the pipette was left in place for 5 min at the injection site before raising it 200 µm above the injection site. After an additional 5 min, the pipette was retracted from the brain. To minimize dehydration during surgery, mice received a subcutaneous injection of 1 mL of sterile saline (Teknova S5819). To reduce inflammation, mice received an injection of Ketoprofen (Zoetis 07-803-7389, 0.01 mg/gram). Postoperatively, mice were monitored on a heat pad for one hour before being returned to their home cage. Mice were then monitored daily for at least 5 days and received either oral carprofen (Rimadyl, bacon flavor, 2 mg/tablet) or a MediGel Carprofe cup (Clear H_2_O) in their home cage.

For hardware implantation (e.g., headplates), one long incision was made to expose the entire dorsal portion of the skull. Excess scalp was removed, and a tissue adhesive (VetBond3M 1469SB) and a catalyst (Zipkicker, Zap PT-29) was applied around the circumference (across 2 rounds of application) of the exposed skull to minimize moisture leaking from surrounding tissue. The skull was cleaned, dried, and scored using a scalpel blade to enhance cohesion of implanted hardware. For fiber implantations, 250 µm craniotomies were made with an electric drill (Foredom Electric Company K.1070) with a ball bur (Busch and Co. S33289) attached to the manipulator. Using a fiber implantation clamp attached to the manipulator (KOPF), fibers were slowly lowered into the brain to the desired coordinates. Glue (Loctite 454) was applied around the base of the fibers and Zipkicker was used to quickly harden the glue. The clamp was released, and more glue and catalyst were applied around the ferrule to ensure stable cohesion to the skull. A final round of cement (C&B Metabond) was applied to the hardened surface for extra security. For headpost implants, titanium headposts were attached on the surface of the skull across the lambda skull suture. This location allowed for injections or fiber implantations in anterior regions. A small amount of glue was applied to the bottom portion of the headpost which was then carefully placed by hand on to the skull. Any necessary adjustments to the position and angle of the headposts were made to keep the implants as flat as possible before applying a catalyst to harden the glue. More glue, catalyst, and cement were applied around the points of contact with the skull and built up over the middle portion of the implant to ensure stable cohesion to the skull. After the hardware or headpost implantation was completed, exposed skull was covered with a UV-Cured dental cement (Pentron N11XVD) to avoid exposure to air, moisture, or contaminants. The same postoperative monitoring and measures were taken for all animal with hardware implants.

Brain coordinates in this study are given with respect to Bregma; anterior–posterior (A/P), medial–lateral (M/L), and dorsal–ventral (D/V). D/V coordinates are measured from the top of the brain.

Coordinates for bilateral virus injections for photometry (**Figure 3-5; Supp Figure 3-5**):

A/P: +0.6
M/L: ± 2.4
D/V: −3.4
(250 nL injected per site)

Coordinates for bilateral photometry fiber implants (**Figure 3-5; Supp Figure 3-5**):

A/P: +0.6
M/L: ± 2.4
D/V: −3.2

Coordinates for bilateral virus injections for optogenetics (**Figure 6; Supp Figure 6**):

A/P: +0.65
M/L: ± 2.4
D/V: −3.2 and −2.5
(200 nL injected per site, 400nL total)

Coordinates for bilateral tapered fiber implants (**Figure 6-7; Supp Figure 6-8**):

A/P: +0.65
M/L: ± 2.4
D/V: −3.4

Viruses (**Figures 3-6, Supp Figure 3-6**):

1) AAV9-syn-FLEX-jGCaMP8m-WPRE (Addgene 162378), final titer=5.0e+12 OR 8e+12
2) AAV9-hsyn-rDA1m(rDA2.5m) (WZ Biosciences Inc.), final titer=5.0e+12
3) AAV5-EF1a-DIO-hChR2(H134R)-mCherry-WPRE-pA (UNC), final titer=5.1e+12
4) AAV5-EF1a-DIO-eYFP (Addgene 27056), final titer=1.0e+13

#### Behavior system & training

Behavioral experiments were conducted in a sound-attenuating chamber (Med Associates ENV-018V) containing a custom-built two-tiered apparatus. Mice were positioned on the upper tier in a detachable head-fixation stage consisting of a clear, copper-lined polycarbonate tube (3.8 cm inner diameter × 10 cm length; McMaster-Carr) with a 2 cm-long half-cylinder opening. The tube was mounted to a circular aluminum breadboard (15 cm × 0.127 cm; Thorlabs MBR6) using an adjustable-height optics clamp (Thorlabs VG100). Custom metal headpost holders were mounted on either side of the tube opening using hex-locking post holders (2.5 cm; Thorlabs PH1) and mounting bases (2.5 cm × 5.8 cm × 1 cm; Thorlabs BA1s). The stage was secured within the apparatus using magnetically coupled kinematic bases (7.6 cm × 7.6 cm; Newport BK-3A). Two steel lick spouts (0.05-inch outer diameter, 0.033-inch inner diameter), separated by 4.5 mm, were mounted on a 2-axis motorized stage (Zaber A-MCB2-KS10A) attached to a post with a manual 3-axis fine-positioning stage (Narishige U-3C). Pure tones were delivered through speakers (Audax TW025A20) mounted on the lower tier. Water rewards delivery was controlled with silent solenoid valves (Lee LHQA0531220H) positioned outside the chamber.

Behavior events (such as licks, cue, and reward dispensation via solenoids) were registered at 200 Hz using custom software running on a MyRio-1900 (LabVIEW 2014, National Instruments). Lick detection was done using a 2-spout lick port and licks were registered by a custom designed resistor-capacitor based system. The system recorded the lick timing at the offset of the tongue touch. This ensured a separation of the first lick (response lick) from the reward delivery and reward collection licks.

Mice were allowed at least 5 days to recover from titanium headpost and/or hardware implantation surgery before water restriction. Following recovery, mice were singly housed, weighed to establish baseline body weight, and then water restricted to 1 mL of water per day. Body weight was maintained above 80% of baseline throughout training. Mice were weighed before and after each behavioral session. Habituation to the task and behavioral apparatus proceeded gradually over several days. On the first day of water restriction, mice were manually handled and given 1 mL of water by syringe. On the second day, mice were introduced to the holding tube and gently passed through it 5-10 times while receiving 1 mL of water. On the third day, mice were head-fixed for 15 minutes outside the behavioral apparatus to monitor for signs of discomfort or stress and again received 1 mL of water by syringe. Over the next 1-2 days, mice were head-fixed in the behavioral apparatus and received “free rewards” from the spouts following a single 75 ms, 5 kHz pure tone, without needing to make a choice lick. Up to 200 rewards were delivered across the acclimatization session in blocks of 5 alternating rewards per side. This procedure familiarized mice to head fixation, trained them to collect rewards efficiently from both spouts, and established that the auditory cue did not predict reward side. If a side bias emerged, manual training was extended at the experimenter’s discretion to encourage equal sampling of both spouts.

After habituation, mice were trained on the head-fixed, two-choice dynamic foraging task. Each trial began with an enforced no-lick (ENL) delay period of 1-2 s (uniformly sampled as 1.0, 1.25, 1.5, 1.75, or 2.0 sec), during which any lick restarted the trial with the same ENL length. A single 75 ms, 5 kHz tone (same as in the acclimatization sessions) signaled the start of a 3 sec selection period, during which the mouse could choose one of the two spouts. Reward contingencies were deterministic and assigned in software such that one spout was rewarded with 100% probability and the other with 0% probability. If the rewarded spout was chosen, 4-5 µL of water was delivered from that spout. A choice lick was followed by a 3 sec consumption period, which occurred regardless of outcome.

Trials were organized into blocks in which one spout was designated as rewarded and the other as unrewarded. At the end of each block, the reward contingencies reversed. Training began with block lengths consisting of 8-9 rewarded trials, allowing mice to learn a “win-repeat” strategy. This block structure generally lasted 1-2 weeks. The number of rewards per block was then reduced to 6-7 rewarded trials and finally to 4-8 rewarded trials (∼4-6 weeks of training), which constituted the late-training and experimental condition. During early training, mice received a free reward after 6 consecutive unrewarded trials to encourage switching to the rewarded spout. These free rewards were omitted during late training and all experimental recording or manipulation sessions. Each session lasted up to 60 minutes or until the mouse collected 200-230 rewards, equivalent to ∼1 mL of water (4-5 µL per reward). If a mouse failed to collect at least 200 rewards, supplemental water was provided to maintain 1 mL/day total intake. Performance criterion for data inclusion, i.e. “late” stage training, was met by achieving a 5-day window with an average of at least >80% rewarded trials, <2 trials to reward after a block switch, >80% correct strategy usage, and <30% of trials with lick penalties. Mice typically reached criterion performance within 4-6 weeks (see **Figure 1; Supp Figure 1**).

#### Fiber photometry

Fiber implants for frequency modulated photometry recordings were connected to a 0.37 NA patch cord (Doric Lenses, MFP_200/220/900-0.37_2m_FCM-MF1.25_LAF, low autofluorescence), which received excitation light and sent its emission light to a Doric filter club: blue excitation light (465-480 nm); red excitation light (555-570 nm); green emission light (500-540 nm); red emission light (580-680 nm) (FMC5_E1(465-480)_F1(500-540)_E2(555-570)_F2(580-680)_S, Doric Lenses). Excitation light originated from LED drivers (Thorlabs) and was amplitude-modulated at 211 (right) or 223 (left) Hz (470-nm excitation light, M470F3, Thorlabs; LED driver LEDD1B, Thorlabs) and 311 (right) or 383 (left) Hz (565-nm excitation light, M565F3, Thorlabs; LED driver LEDD1B, Thorlabs) using THT RX8 digital signal processor lock-in amplifier. The following excitation light powers were used for the indicated sensors: GcAMP8m (25 µW) and rDAm (45 µW). Raw photometry signals from the photodetectors were amplified in DC mode with Doric photodetectors and collected through the TDT interface at 6103.52 Hz. The TDT interface also received trial start-pulses from the LabVIEW behavioral system. Behavior and photometry were aligned post-hoc using timestamps from LabVIEW and trial start-pulses fed into the TDT system. Recordings occurred simultaneously on both hemispheres. To minimize data loss from z-scoring in post-hoc analysis, a 5-10-minute baseline period was acquired pre and post behavior.

Photometry signals were processed in Python using custom scripts. Raw frequency-modulated photometry traces were first detrended with a rolling z-score using a 180 sec window. This step was applied before demodulation to reduce slow baseline drift and photobleaching. Because fluorescence-dependent changes in the raw modulated signal are small relative to the carrier modulation, and because behavioral events occurred on a much faster timescale than the detrending window, this preprocessing removed slow changes in baseline without substantially distorting trial-related fluctuations. Detrended signals were then demodulated using a rolling spectrogram (scipy.signal.spectrogram) with a Hamming window. Spectrogram segments were computed with nperseg=958 samples and noverlap=835.93 samples, yielding a step size of 122.07 samples and an effective output sampling rate of 50 Hz. The corresponding spectral resolution was 6.37 Hz per bin. For each channel, the demodulated signal was defined as the mean power in the frequency bins nearest the measured carrier frequency, rather than as power within a fixed-width frequency band. No additional low-pass filter was applied beyond the smoothing inherent to the spectrogram windowing.

For subsequent fluorescence quantification, the demodulated detrended trace was subjected to a second rolling z-score normalization using a 180 sec window to generate the final z-scored photometry signal used for analysis. To align photometry and behavior, trial-start pulses from the behavioral control system and selection licks were matched to the digital event stream marking trial starts. This alignment was then used to register all other behavioral events and lick times to the photometry signal. Trial-aligned fluorescence traces were then averaged across trials and mice, with no normalization after z-scoring.

For analysis, only behavioral sessions from well-trained mice were accepted. Behavioral analysis and quality control for all photometry animals was performed as described previously. Every fiber photometry signal from all sessions was quality controlled for subsequent analysis. Measurements of skew, kurtosis, bleaching, and signal to noise ratio were calculated on each sessions z-scored signal per hemisphere. Further, all raw, detrended, and z-scored signals were manually inspected to ensure reliability of transients across the session. Only sessions with skew >1, kurtosis >3 (excess kurtosis), and bleaching <15% were included for subsequent analysis. Finally, post-hoc histology confirmed viral expression and fiber placement in all animals. After signal quality control and histological verification, 65 sessions (7-9 sessions per animal) for 8 D1-Cre mice and 120 sessions (6-7 sessions per animal) for 18 A2a-Cre mice were included in the final dataset. A subset of *Drdla-Cre* (N=6) and *Adora2a-Cre* (N=11) mice were co-injected with AAV9-hSyn-rDAm and GcAMP8m at the exact same viral coordinates and fiber implantation (**Supp Figure 4**, mice denoted with *).

#### Photostimulation and inhibition

Once mice were fully trained, optogenetic stimulation or inhibition was accomplished using a 473 nm laser launched into a fiber optic (Laser Quantum, Ventus + MCP 600, DPSS) and controlled by an acousto-optic modulator (AOM; AA Opto-Electronic, MTS110-A3-VIS) and a shutter (Vincent Associates). The AOM and shutter were controlled using custom LabVIEW code. For all optogenetic experiments, tapered optical fibers (Optogenix, 0.39 NA/200 µm fiber, active length 2.5 mm) were bilaterally implanted in the ventrolateral striatum (VLS), and only one fiber was stimulated per session. Output power was calibrated daily at the end of the patch cord prior to each experiment and adjusted to 0.5-1 mW depending on genotype and condition. Light was delivered on approximately 15% of trials to minimize persistent behavioral effects caused by repeated stimulation.

Each session included stimulated trials during both left- and right-rewarded blocks to assess ipsilateral and contralateral effects, and 2-4 stimulation sessions were performed per fiber/hemisphere per mouse. For each mouse, experiments were completed on one fiber, alternating between left and right fibers across animals, before moving to the second fiber. Although we did not observe gross persistent effects on baseline performance across sessions, all optogenetic experiments were separated by at least 1 baseline day to monitor for any lasting effects of stimulation or inhibition on task performance.

For photoactivation experiments, ChR2-mCherry or eYFP was expressed in dSPNs or iSPNs in the VLS. Stimulation was delivered during the action-outcome association period after the choice lick and was triggered by the correct choice lick on win-repeat trials only, with an onset latency of approximately 10 µs. To control for reward history, stimulation was delivered only on the third rewarded trial within a block. Light was delivered at 20 Hz for 3 sec at 1 mW, spanning the entire action-outcome association period. The dSPN photoactivation cohort consisted of D1-ChR2 mice (N=18 mice, 75 sessions) and D1-eYFP controls (N=4 mice, 21 sessions). The iSPN photoactivation cohort consisted of A2a-ChR2 mice (N=6 mice, 32 sessions) and A2a-eYFP controls (N=6 mice, 22 sessions). Within the A2a-ChR2 mice, 3 mice expressed ChR2 via the Ai32 allele (A2a; Ai32) rather than viral delivery.

For photoinhibition experiments, D1-Cre or A2a-Cre mice were crossed with floxed GtACR1 mice. Hemizygous Cre-negative GtACR1-floxed mice were used as light controls. As in the stimulation experiments, inhibition during the action-outcome association period was triggered by the correct choice lick on win-repeat trials only and delivered on the third rewarded trial within a block to control for reward history. Continuous light was delivered for 3 sec at 0.5 mW in D1;GtACR1 mice, and 3 sec at 1 mW in A2a;GtACR1 and GtACR1 control mice. For action-outcome association-period inhibition experiments, the cohorts consisted of D1;GtACR1 mice (N=8 mice, 27 sessions), A2a;GtACR1 mice (N=11 mice, 24 sessions), and GtACR1 control mice (N=5 mice, 18 sessions). For ENL delay-period inhibition experiments, continuous light was delivered for 1.5 sec using the same genotype-specific power levels, again on the third rewarded trial within a block to control for reward history. The ENL delay-period cohorts consisted of D1;GtACR1 mice (N=7 mice, 28 sessions), A2a;GtACR1 mice (N=6 mice, 18 sessions), and GtACR1 control mice (N=5 mice, 18 sessions).

All five Cre-negative GtACR1-floxed mice were used as controls for both the action-outcome association-period and ENL delay-period inhibition experiments; action-outcome association-period testing was performed first in these control animals. In addition, 3 D1;GtACR1 mice and 3 A2a;GtACR1 mice were tested in both the action-outcome association-period and the ENL delay-period inhibition paradigms. Among these overlapping experimental animals, two mice were tested first in the ENL delay period, and one mouse was tested first in the action-outcome association-period manipulation.

For all photostimulation and inhibition experiments, post-hoc histology confirmed viral expression and fiber placement in all animals. As tapered fibers have sub-micron-sized tips and produce smaller tissue damage compared to flat-cleaved fibers (Pisanello et al., 2017), glial fibrillary acidic protein (GFAP) immunohistochemistry was used to localize fiber tips. Fiber tips were unable to be located in 4 D1-ChR2 mice (**Supp Figure 6**, labeled as mice 15-18 on the figure), although viral expression was consistent with other animals in the cohort.

To prevent mice from distinguishing stimulation and control trials based on visual cues from blue light, carbon powder was added to the dental cement holding the fiber in place to mask emitted light, and black heat-shrink tubing was used to physically cover the connection between the tapered fiber and patch cord.

#### *Ex vivo* electrophysiology

Brain slices were prepared from 2-3 month-old A2a;GtACR1 or D1;GtACR1 mice of both sexes using standard techniques. The experiments closely followed the procedures outlined in previous studies (Ganesh et al., 2026; Kim et al., 2022; Wallace et al., 2017). Mice were anesthetized using isoflurane inhalation and subsequently subjected to transcardial perfusion with ice-cold artificial cerebrospinal fluid (ACSF) composed of the following (in mM): 125 NaCl, 2.5 KCl, 25 NaHCO3, 2 CaCl2, 1 MgCl2, 1.25 NaH2PO4, and 11 glucose (final osmolarity of 300–305 mOsm/kg). The brain was removed from the skull, placed in ice-cold ACSF, and 300 μm coronal brain slices were obtained on a Leica VT1000 S vibratome. Slices were initially placed in a holding chamber for 10 minutes at 34°C in a choline-based solution composed of (in mM): 110 choline chloride, 25 NaHCO3, 2.5 KCl, 7 MgCl2, 0.5 CaCl2, 1.25 NaH2PO4, 25 glucose, 11.6 ascorbic acid, and 3.1 pyruvic acid. Following the choline incubation, slices were transferred to a second chamber with ACSF also maintained at 34°C for a minimum of 30 minutes. The chamber was subsequently shifted to room temperature for the duration of the experiment. All recordings were made within 4-5 hours of slicing.

Striatal projection neurons were targeted for recordings using differential interference contrast (DIC) optics on an upright Scientifica microscope equipped with a digital camera (Hamamatsu), an epifluorescence LED light source (CoolLED), and a 60x/0.9 NA LUMPlanFl/IR water-immersion objective (Olympus). Patch pipettes were pulled from borosilicate glass (Sutter Instruments) with resistances of 2.0-3.5 MΩ. Recordings were performed using a MultiClamp 700B amplifier (Molecular Devices) and a data acquisition board (National Instruments), and data were collected using MATLAB custom-written software and scripts (R2023a). Striatal neurons were identified by their FusionRed-positive cell bodies. Fluorescence-negative neurons with GABAergic interneuron physiological properties (membrane tau decay<0.8 ms for both fast-spiking and persistent low-threshold spiking subtypes; input resistance>500 MΩ in persistent low-threshold spiking subtype) were excluded from the analysis. The temperature was maintained at 32°C and carbogen-bubbled ACSF was perfused at a rate of 2-3 mL/min for all experiments

For whole-cell current-clamp recordings, patch pipettes were filled with low chloride potassium-based internal solution (in mM): 135 KMeSO_3_, 3 KCl, 10 HEPES, 1 EGTA, 0.1 CaCl_2_, 4 Mg-ATP, 0.3 Na-GTP, and 8 Na_2_Phosphocreatine (pH 7.3 adjusted with KOH, 290-300 mOsm/kg). Signals were filtered at 10 kHz and digitized at 50 kHz. To validate GtACR1 function in slice, cells were given current injections (200-600 pA) to produce stable spiking. After 500 ms, a 2 ms light pulse (473 nm) was delivered at light powers of 0.5 or 1 mW whose power output was controlled through an acousto-optic modulator to assess the ability of GtACR1 to inhibit spiking. Holding current and input resistance were continuously monitored as proxies of recording stability. No series resistance compensation was used. Series resistance did not exceed 20 MΩ.

For whole-cell voltage clamp recordings, patch pipettes were filled with a cesium-based internal solution (in mM): 135 CsMeSO_3_, 10 HEPES, 1 EGTA, 3.3 QX-314 (Cl^−^ salt), 4 Mg-ATP, 0.3 Na-GTP, 8 Na_2_-Phosphocreatine, with pH adjusted to 7.3 using CsOH, resulting in an osmolarity of 295 mOsm/kg. All recorded neurons exhibited electrophysiological characteristics of spiny projection neurons. Photocurrents were recorded at a holding potential of 0 mV. Series resistance and leak currents were monitored continuously. GtACR1-mediated currents were optically evoked using 500 ms light pulses (473 nm continuous light) at light powers of 0.25, 0.5, and 1 mW whose power output was controlled through an acousto-optic modulator. Quality control criteria were a stable baseline membrane resistance (variation in R_m_<10%) and a stable baseline resting potential with median standard deviation of ≤0.5 mV across recordings. All data were analyzed post-hoc using custom scripts in Igor Pro 6.0 software (Wavemetrics).

#### Histology and microscopy

Animals were anesthetized with isoflurane and transcardially perfused with phosphate-buffered solution (PBS, 0.081 mM 1701 Na2HPO4, 0.017 mM NaH2PO4, pH 7.2-7.4), followed by 4% paraformaldehyde (PFA) in PBS. Brains were dissected from the skull, post-fixed overnight in 4% PFA at 4°C, and transferred to a 30% sucrose solution the following day. Brains were stored at 4°C until slicing. Coronal slices (50–80 µm) were obtained using a cryostat. For immunohistochemistry, sections were washed in PBST (0.1% Triton X-100), blocked in 6% normal goat serum (NGS; Jackson ImmunoResearch, #AB144P) or bovine serum albumin (BSA), and permeabilized in 0.1% Triton X-100 for 2 hours at room temperature on a shaker. Primary antibodies were diluted in 3% NGS with 0.1% Triton X-100 and incubated with sections overnight at 4°C in Eppendorf tubes on a rotating tube rack. Primary antibodies used were Rabbit anti-Gilial Fibrillary Acidic Protein (GFAP, Agilent, 1:1,000), Goat anti-ChAT (Millipore, 1:500), Rabbit anti-DARPP-32 (Cell Signaling Technology, 1:1,000), Rabbit anti-PV (Swant, 1:2,000), Chicken anti-GFP (Abcam, 1:1,500). Sections were then washed in PBST, incubated in secondary antibodies (donkey anti-rabbit, goat, or chicken Alexa Fluor 488, 593, or 647, 1:500, JacksonImmuno Research) for 2-4 hours at room temperature on a shaker, washed, and stained with DAPI (1:10,000). Finally, sections were washed, mounted onto slides, cover slipped with Vectashield Plus Antifade Mounting Medium. All images were acquired on a VS200 slide scanner (Olympus) or a spinning-disk confocal microscope (Core for Imaging Technology and Education).

### QUANTIFICATION AND STATISTICAL ANALYSIS

#### Statistics

All statistical comparisons can be found in Table S1. Data were analyzed using Python v3.11.3 and statistical tests were performed using SciPy functions. No statistical method was used to pre-determine sample size. Data were not systematically tested for normalcy; thus, non-parametric, two-sided statistical tests were used for all analysis. Statistical significance was taken as *p* < 0.05. In all figures, n.s: *p* > 0.05 and ∗*: p* < 0.05. All graphs show the mean and standard error about the mean (SEM), with “N” referring to the number of animals and “n” referring to the number of sessions, unless otherwise stated.

#### Behavior

Unless otherwise specified, all behavioral metrics were computed by pooling data across all sessions within each mouse to generate a single value per mouse, and then averaging across mice within each condition; group data are shown as mean ± SEM. To quantify learning (**Figure 1**), we compared behavior within mice during “early” training (days 5-7) versus “late” training, defined as three randomly selected sessions after performance criterion was reached, ∼4-6 weeks after training. Thus, each mouse contributed a single value representing the average across the selected sessions. For regression analysis, Pearson’s correlation was used (**Supp Figure 1**). Data in Figure 2 included only sessions from well-trained (late) animals. For probability and cumulative distribution plots (**Figure 2B; Supp Figure 2A)**, rewarded win-repeat trials and unrewarded lose-switch trials were identified, and lick times were binned into 200 ms intervals. For each mouse, we calculated the probability of observing at least one lick in each bin, normalized by the total number of included trials, as well as the corresponding cumulative distribution function (CDF). These data were plotted as across-mouse averages ± SEM. For directional delay-period licking analyses (**Figure 2E**), only trials containing at least one penalty lick were included. Within each trial, penalty licks were classified as occurring in the same or opposite direction relative to the subsequent choice before, normalized by the total number of ENL periods, and averaged across sessions within each mouse for win-repeat and lose-switch conditions. For first/last delay-period lick analyses (**Supplementary Figure 2B**), only trials containing more than one ENL period were included. The first and last ENL penalty licks were classified as occurring in the same or opposite direction relative to the subsequent choice before averaging probabilities across sessions within each mouse for win-repeat and lose-switch conditions. Statistical comparisons for reaction times for all win-repeat and lose-switch trials (**Figure 2F; Supp Figure 2C**) were performed on the mean reaction time for each trial type in each mouse. Figure 2F includes trials regardless of whether they were preceded by lick penalties, whereas trials with and without preceding lick penalties are shown separately in Supplementary Figure 2C.

#### Fiber photometry

Only sessions passing behavioral and photometry signal quality control were included. For each hemisphere, right and left choices were remapped as ipsilateral and contralateral relative to the recording site. Each trial contributed one observation per hemisphere. We quantified photometry data in two ways. First, we measured the mean z-scored signal (“avg signal”) within a defined time window after the event of interest (**Supp Figure 3B-C, Supp Figure 4B-C**). This was calculated as the average z-scored value of the photometry trace within that window. The analysis window was 750 ms for spontaneous licks, 3 sec for the outcome period, and 1 sec for the ENL preceding the cue. Second, we measured the peak z-scored signal within a defined time window, calculated as the maximum or minimum value of the mouse-averaged photometry trace over that window. The window was 0.25 s for spontaneous licks and the outcome period. To test for potential differences between the recorded signal in the right and left hemisphere for task-independent and task-related events, we compared transients across hemispheres and quantified the average z-scored and peak signal. In general, trials with and without preceding ENL penalty licks were included. Repeated-measures group differences were assessed using the Friedman test, followed by post hoc pairwise Wilcoxon signed-rank tests; p-values were adjusted for multiple comparisons using the Benjamini-Hochberg false discovery rate (FDR) procedure. Adjusted p-values are reported.

#### Decoding of future choice and outcome history

To examine if striatal activity carried information predictive of the animal’s upcoming (trial n+1) and past (trial n-1) spout choice, we used a sliding-window receiver operating characteristic (ROC) analysis to discriminate future switching or outcome history. Trials were grouped according to the animals upcoming choice or previous trial outcome.

To determine if striatal activity held information about upcoming choice, calcium activity in dSPNs or iSPNs was aligned to the action-outcome association period (trial n), and predictive labels were assigned on whether the animal would switch spouts on the subsequent trial (trial n+1) **(Figure 5A-E**). Prior to decoding, the photometry signal was downsampled using a non-overlapping mean filter with a window of 4 samples (effective sampling rate: 12.5 Hz). Classes were balanced prior to model fitting by randomly downsampling the majority class to match the size of the minority class. A logistic regression classifier was then fit across the same time bin extracted from each trial to discriminate the binary label of interest (switch vs. stay on the subsequent trial), using the mean signal within that bin as the sole predictor. Model performance was quantified as the area under the ROC curve (defined as ROC AUC) such that 0.5 indicates chance-level discrimination. This procedure was repeated across successive non-overlapping time bins spanning the full consumption window, yielding a continuous AUC time course. AUC values were computed separately for each mouse.

To test whether striatal activity retained information about reward history from the previous trial, we aligned photometry signals to the cue on trial n and labeled each trial according to the outcome of trial n-1 (**Figure 5F-J**). Specifically, the binary logistic regression distinguished whether the preceding trial had been rewarded or unrewarded. Analyses were conducted separately for each ENL duration to account for the variable delay preceding the cue. To summarize decoding performance across ENL conditions, peak AUC was computed per mouse as the maximum AUC value across the full alignment window, and group data are shown as mean ± SEM across mice as a function of ENL duration.

To assess statistical significance, a permutation-based null distribution was generated by randomly shuffling the behavioral labels 1000 times and recomputing the mean AUC across mice at each time bin on each iteration. Significance at each time point was determined by comparing the observed mouse-averaged AUC time course against the permutation-based null distribution using a nonparametric two-sided Mann-Whitney U test, with p-values corrected for multiple comparisons across time bins using the Holm-Bonferroni sequential correction procedure (α=0.05).

#### Photostimulation and inhibition

For switching analyses, the stimulated trial was defined as trial n, and the behavioral effect of stimulation was assessed on the subsequent trial (n+1). Switching was quantified by comparing the animal’s choice on trial n+1 with its choice on trial n. If the animal selected the same spout on trial n+1 as on trial n, the trial was classified as a stay (win-repeat); if it chose the opposite spout, the trial was classified as a switch (win-switch). Choices were defined relative to the stimulated hemisphere: selections toward the stimulated side were labeled ipsilateral, and selections toward the opposite side were labeled contralateral. Each post-stimulation trial was therefore assigned to one of four categories: ipsilateral stimulation leading to an ipsilateral stay (I→I), ipsilateral stimulation leading to a contralateral switch (I→C), contralateral stimulation leading to a contralateral stay (C→C), or contralateral stimulation leading to an ipsilateral switch (C→I).

Non-optogenetic (non-opto) control trials were drawn from the same sessions and were defined using the same criteria as the stimulated trials, except that any trials immediately following an optogenetic manipulation as excluded from the control set. These non-opto trials were classified using the same ipsilateral/contralateral framework and the same stay-versus-switch criteria as optogenetic trials. Trials in which the animal failed to make a choice were excluded from all switching analyses. These omission trials represented ∼1.1% of stimulated trials across (**Figures 6-7; Supp Figures 6-8**). Percent switch was calculated separately for ipsilateral and contralateral trial types in both optogenetic and non-optogenetic conditions.

Ipsilateral switch probability was calculated as: I→C / (I→C + I→I) × 100
Contralateral switch probability was calculated as C→I / (C→I + C→C) × 100

Lick metrics were quantified for trials with stimulation or inhibition during the action-outcome association period (**Supp Figures 6-7**). The mean number of licks was calculated for the manipulated trials (trial n) and compared with all other win-repeat trials in the same session in which no stimulation or inhibition occurred; manipulated trials were excluded from the control set. Lick frequency was calculated as the inverse of the mean interlick interval within each lick bout. For analyses of licking direction, the chosen spout was defined as the spout contacted on the choice lick. During consumption, the fraction of licks directed to the chosen spout was calculated as chosen-spout consumption licks divided by total consumption licks, and the fraction directed to the non-chosen spout was calculated as non-chosen-spout consumption licks divided by total consumption licks. Reaction times were plotted as cumulative distributions and separated into non-stimulated trials, stimulated trials that did not result in switching, and stimulated trials that resulted in switching (**Supp Figures 6-8**). Two to four stimulation sessions were performed for each fiber/hemisphere per mouse. Sessions were averaged within each mouse, and the resulting mouse values were averaged to obtain group mean ± SEM. Repeated-measures group differences were assessed using the Friedman test, followed by post hoc pairwise Wilcoxon signed-rank tests; p values were adjusted for multiple comparisons using the Benjamini-Hochberg FDR procedure and adjusted p-values are reported.

#### Histology, Microscopy, and Cell Counting

For immunohistochemical experiments to investigate colocalization between D1 or A2a;GtACR1-FusionRed positives cells and markers of striatal cell types (**Supp Figure 7**), three representative coronal slices per animal were chosen, focusing next on three specific fields of view in the VLS. Confocal z stacks in each field of view were acquired, matching exposure time and laser power for each fluorophore in all images captured. To assess colocalization between FusionRed cells and different neuronal markers, maximum z-projections were made and merged to produce an overlay with FusionRed cells and the corresponding cellular marker. For each animal, a minimum of 50 cells of each type (FusionRed or cellular marker) were manually counted using custom scripts in FIJI/ImageJ. For a cell to be counted as colocalized, it needed to appear on both individual and merged channels in the z-stack and maximum z-projection showing identical overlap with the other cell. Any cell showing ambiguous overlap was not counted as colocalized.

## Notes

### Competing Interest Statement

The authors have declared no competing interest.

