## Supplemental Table 1 for "Striatal activity maintains a short-term action-outcome memory to guide future choice"

| Key Experiments & Analysis | Figure | Type of Comparison | Statistical Test | n (sessions) | N (mice) | Comparison values (mean ± SEM) | Stastic, p-value | Post hoc corrected p-value |
|---|---|---|---|---|---|---|---|---|
| Number of trials | 1d | Within-mouse, early vs. late | WSR | 42 (early), 45 (late) | 15 | Early: 158.27 ± 10.38, Late: 221.47 ± 4.15 trials | 1.22E-04 | N/A |
| % Trials rewarded trials | 1d | Within-mouse, early vs. late | WSR | 42 (early), 45 (late) | 15 | Early: 61.2 ± 2.0%, Late: 90.0 ± 1.0% reward rate | 6.10E-05 | N/A |
| Trials to obtain reward | 1d | Within-mouse, early vs. late | WSR | 42 (early), 45 (late) | 15 | Early: 5.32 ± 0.42, Late: 1.44 ± 0.07 trials to reward | 6.10E-05 | N/A |
| % Trials with lick penalities during ENL delay epoch | 1d | Within-mouse, early vs. late | WSR | 42 (early), 45 (late) | 15 | Early: 57.71 ± 3.35%, Late: 20.90 ± 2.77% trials with penalities | 6.10E-05 | N/A |
| Correct strategy | 1f | Within-mouse, early vs. late | WSR | 42 (early), 45 (late) | 15 | Early: 66.64 ± 1.58%, Late: 88.63 ± 1.11% correct strategy | 6.10E-05 | N/A |
| Log lose-switch/lose-stay | 1f | Within-mouse, early vs. late | WSR | 42 (early), 45 (late) | 15 | Early: -0.47 ± 0.05, Late: 0.35 ± 0.07 log(lose-switch/lose-stay) | 6.10E-05 | N/A |
| % Trials with lick penalities during cue epoch | Supp 1a | Within-mouse, early vs. late | WSR | 42 (early), 45 (late) | 15 | Early: 6.61 ± 0.86%, Late: 0.83 ± 0.17% trials with cue penalities | 6.10E-05 | N/A |
| Side bias, correct strategy | Supp 1d | Across sesions, left vs. right | Pearson Corr | 42 (early), 45 (late) | 15 | r = 0.80, $R^2$ = 0.64 | 3.24E-20 | N/A |
| Side bias, fraction of rewarded trials | Supp 1d | Across sesions, left vs. right | Pearson Corr | 42 (early), 45 (late) | 15 | r = 0.70, $R^2$ = 0.49 | 1.89E-13 | N/A |
| Side bias, trials to obtain reward | Supp 1d | Across sesions, left vs. right | Pearson Corr | 42 (early), 45 (late) | 15 | r = 0.73, $R^2$ = 0.53 | 1.5E-15 | N/A |
| Sex bias, correct strategy | Supp 1e | Across-mice, M vs. F, late | MWU | 27 (9 M), 18 (6 F) | 15 | Male: 85.60 ± 1.72%, Female: 87.17 ± 1.32 % correct strategy | 0.69 | N/A |
| Sex bias, fraction of rewarded trials | Supp 1e | Across-mice, M vs. F, late | MWU | 27 (9 M), 18 (6 F) | 15 | Male: 86.20 ± 1.77%, Female: 89.34 ± 1.80 % trials rewarded | 0.22 | N/A |
| Sex bias, trials to obtain reward | Supp 1e | Across-mice, M vs. F, late | MWU | 27 (9 M), 18 (6 F) | 15 | Male: 1.52 ± 0.09, Female: 1.24 ± 0.09 trials to obtain reward | 0.11 | N/A |
| Mean # licks during consumption bout | 2c | Within-mouse, late | WSR | 45 | 15 | win-repeat: 16.30 ± 1.03 licks, lose-switch: 3.32 ± 0.35 licks | 6.10E-05 | N/A |

| | | | | | | | | |
|---|---|---|---|---|---|---|---|---|
| Lick direction toward current choice action: win-repeat | 2d | Within-mouse, late | N/A | 45 | 15 | win-repeat chosen: 92.21 ± 2.27% & win-repeat not chosen: 7.64 ± 2.26% | N/A | N/A |
| Lick direction toward current choice action: lose-switch | 2d | Within-mouse, late | N/A | 45 | 15 | lose-switch chosen: 74.25 ± 3.34% & lose-switch not chosen: 25.61 ± 3.32% | N/A | N/A |
| Probability of ENLP penalty, win-repeat chosen vs not chosen | 2e | Within-mouse, late | WSR | 45 | 15 | win-repeat chosen: 0.76 ± 0.031 & not chosen: 0.24 ± 0.030 | 6.10E-05 | N/A |
| Probability of ENLP penalty, lose-switch chosen vs not chosen | 2e | Within-mouse, late | WSR | 45 | 15 | lose-switch chosen: 0.84 ± 0.030 & not chosen: 0.16 ± 0.030 | 6.10E-05 | N/A |
| Selection time, win-repeat vs. lose-switch | 2f | Within-mouse, late | WSR | 45 | 15 | win-repeat: 111.07 ± 7.65 ms, lose-switch: 149.20 ± 13.25 ms | 2.00E-04 | N/A |
| Probability of ENLP, First Lick, win-repeat chosen vs not chosen | Supp 2b | Within-mouse, late | WSR | 45 | 15 | win-repeat chosen: 0.72 ± 0.05 & not chosen: 0.28 ± 0.05 | 0.0012 | N/A |
| Probability of ENLP, First Lick, lose-switch chosen vs not chosen | Supp 2b | Within-mouse, late | WSR | 45 | 15 | lose-switch chosen: 0.70 ± 0.04 & not chosen: 0.30 ± 0.04 | 8.60E-04 | N/A |
| Probability of ENLP, Last Lick, win-repeat chosen vs not chosen | Supp 2b | Within-mouse, late | WSR | 45 | 15 | win-repeat chosen: 0.69 ± 0.05 & not chosen: 0.31 ± 0.05 | 0.0034 | N/A |
| Probability of ENLP, Last Lick, lose-switch chosen vs not chosen | Supp 2b | Within-mouse, late | WSR | 45 | 15 | lose-switch chosen: 0.67 ± 0.05 & not chosen: 0.33 ± 0.05 | 0.012 | N/A |
| Selection time, win-repeat no penalty vs. win-repeat with penalty | Supp 2c | Within-mouse, late | WSR | 45 | 15 | WR no penalty: 113 ± 8.16 ms vs. WR with penalty: 100.80 ± 6.52 ms | 0.0053 | N/A |
| Selection time, lose-switch no penalty vs. lose-switch with penalty | Supp 2c | Within-mouse, late | WSR | 45 | 15 | LS no penalty: 152 ± 15.0 ms vs. LS with penalty: 137.27 ± 10.4 ms | 0.21 | N/A |
| DA, spont lick, min peak | 3e | ipsi vs. contra, within-mouse | WSR | 110 | 17 | contra: -0.71 ± 0.03 vs. ipsi: -0.73 ± 0.03 | Statistic = 49.0, p = 0.21 | N/A |
| DA-outcome, reward, no reward, ipsi, contra, max peak | 3f | reward vs. no reward (ipsi/contra), within-mouse | Friedman + WSR post hoc | 110 | 17 | C-R: 2.54 ± 0.12, I-R 2.41 ± 0.12, C-U: -1.11 ± 0.04, I-U: -1.12 ± 0.05 | Friedman χ² = 43.0, p = 0.000002 | C-R vs. I-R: 0.1, C-R vs. C-U: 0.00009, C-R vs. I-U: 0.00009, I-R vs. C-U: 0.00009, I-R vs. I-U: 0.00009, C-U vs. I-U: 1.0 |
| DA, consecutive rewards, max peak | 3h | 1, 2, 4, 6, 8 rewards, within-mouse | Friedman + WSR post hoc | 110 | 17 | 1: 3.26 ± 0.15, 2: 2.12 ± 0.11, 4: 2.27 ± 0.10, 6: 2.53 ± 0.10, 8: 2.90 ± 0.22 | Friedman χ² = 56.0, p = 0.000001 | 1.0 vs. 2.0: 0.00015, 1.0 vs. 4.0: 0.00015, 1.0 vs. 6.0: 0.00015, 1.0 vs. 8.0: 0.45 |
| DA, consecutive losses, min peak | 3g | 1, 2, 3, 4, 5, losses, within-mouse | Friedman + WSR post hoc | 110 | 17 | 1: -1.27 ± 0.044, 2: -1.03 ± 0.042, 3: -0.99 ± 0.043, 4: -0.93 ± 0.046, 5: -0.95 ± 0.050 | Friedman χ² = 39.7, p = 0.000005 | 1.0 vs. 2.0: 0.00015, 1.0 vs. 3.0: 0.00015, 1.0 vs. 4.0: 0.00015, 1.0 vs. 5.0: 0.00015 |

| | | | | | | | | |
|---|---|---|---|---|---|---|---|---|
| DA, spont lick, min peak, right hemisphere | Supp 3a | contra vs. ipsi (right hemisphere), within-mouse | WSR | 110 | 17 | contra: -0.74 ± 0.050 vs. ipsi: -0.74 ± 0.042 | Statistic = 71.0, p = 0.82 | N/A |
| DA, spont lick, avg signal, right hemisphere | Supp 3a | contra vs. ipsi (right hemisphere), within-mouse | WSR | 110 | 17 | contra: -0.36 ± 0.038 vs. ipsi: -0.38 ± 0.030 | Statistic = 75.0, p = 0.96 | N/A |
| DA, spont lick, min peak, left hemisphere | Supp 3a | contra vs. ipsi (left hemisphere), within-mouse | WSR | 110 | 17 | contra: -0.81 ± 0.042 vs. ipsi: -0.79 ± 0.040 | Statistic = 73.0, p = 0.89 | N/A |
| DA, spont lick, avg signal, left hemisphere | Supp 3a | contra vs. ipsi (left hemisphere), within-mouse | WSR | 110 | 17 | contra: -0.42 ± 0.033 vs. ipsi: -0.39 ± 0.034 | Statistic = 71.0, p = 0.82 | N/A |
| DA, ipsi spont lick, min peak | Supp 3a | right vs. left hemisphere, within-mouse | MWU | 110 | 17 | ipsi right:-0.74 ± 0.042 vs. ipsi left: -0.79 ± 0.040 | Statistic = 128.0, p = 0.58 | N/A |
| DA, contra spont lick, min peak | Supp 3a | right vs. left hemisphere, within-mouse | MWU | 110 | 17 | contra right: -0.74 ± 0.050 vs. contra left: -0.81 ± 0.042 | Statistic = 101.0, p = 0.14 | N/A |
| DA, ipsi spont lick, avg signal | Supp 3a | right vs. left hemisphere, within-mouse | MWU | 110 | 17 | ipsi right: -0.38 ± 0.028 vs. ipsi left: -0.39 ± 0.034 | Statistic = 140.0, p = 0.89 | N/A |
| DA, contra spont lick, avg signal | Supp 3a | right vs. left hemisphere, within-mouse | MWU | 110 | 17 | contra right: -0.36 ± 0.038 vs. contra left: -0.42 ± 0.033 | Statistic = 118.0, p = 0.37 | N/A |
| DA, Cue event response, max peak | Supp 3b | right vs. left hemisphere, within-mouse | MWU | 110 | 17 | cue right: 3.00 ± 0.13, cue left: 3.14 ± 0.13 | Statistic = 130.0, p = 0.63 | N/A |
| DA, Choice event response, max peak | Supp 3b | right vs. left hemisphere, within-mouse | MWU | 110 | 17 | choice right: 1.73 ± 0.077, choice left: 1.68 ± 0.078 | Statistic = 152.0, p = 0.81 | N/A |
| DA, Outcome event response, max peak | Supp 3b | right vs. left hemisphere, within-mouse | MWU | 110 | 17 | outcome right: 1.74 ± 0.069, outcome left: 1.67 ± 0.074 | Statistic = 163.0, p = 0.54 | N/A |
| DA, Cue event response, avg signal | Supp 3b | right vs. left hemisphere, within-mouse | MWU | 110 | 17 | cue right: 0.23 ± 0.012, cue left: 0.22 ± 0.013 | Statistic = 157.0, p = 0.68 | N/A |
| DA, Choice event response, avg signal | Supp 3b | right vs. left hemisphere, within-mouse | MWU | 110 | 17 | choice right: 0.134 ± 0.013, choice left: 0.12 ± 0.015 | Statistic = 167.0, p = 0.45 | N/A |
| DA, Outcome event response, avg signal | Supp 3b | right vs. left hemisphere, within-mouse | MWU | 110 | 17 | outcome right: 0.087 ± 0.0097, outcome left: 0.081 ± 0.012 | Statistic = 165.0, p = 0.49 | N/A |
| DA-outcome, reward, no reward, ipsi, contra, avg signal | Supp 3c | reward vs. no reward (ipsi/contra), within-mouse | Friedman + WSR post hoc | 110 | 17 | C-R: 0.26 ± 0.02, I-R 0.23 ± 0.02, C-U: -0.31 ± 0.02, I-U: -0.32 ± 0.02 | Friedman $\chi^2$ = 41.3, p = 0.00002 | C-R vs. I-R: 1.0, C-R vs. C-U: 0.00009, C-R vs. I-U: 0.00009, I-R vs. C-U: 0.00009, I-R vs. I-U: 0.00009, C-U vs. I-U: 1.0 |

| | | | | | | | | |
|---|---|---|---|---|---|---|---|---|
| DA, consecutive rewards, avg signal | Supp 3d | 1, 2, 4, 6, 8 rewards, within-mouse | Friedman + WSR post hoc | 110 | 17 | 1: 0.27 ± 0.02, 2: 0.19 ± 0.013, 4: 0.25 ± 0.016, 6: 0.27 ± 0.016, 8: 0.29 ± 0.02 | Friedman χ² = 35.6, p = 0.00007 | 1.0 vs. 2.0: 0.00015, 1.0 vs. 4.0: 1.0, 1.0 vs. 6.0: 1.0, 1.0 vs. 8.0: 0.51 |
| DA, consecutive losses, avg signal | Supp 3e | 1, 2, 3, 4, 5, losses, within-mouse | Friedman + WSR post hoc | 110 | 17 | 1: -0.33 ± 0.02, 2: -0.30 ± 0.020, 3: -0.31 ± 0.02, 4: -0.30 ± 0.02, 5: -0.27 ± 0.02 | Friedman χ² = 15.9, p = 0.0032 | N/A |
| dSPN, spont lick, max peak | 4d | ipsi vs. contra, within-mouse | WSR | 65 | 8 | contra: 0.79 ± 0.04 vs. ipsi: 0.73 ± 0.06 | Statistic = 14.0, p = 0.64 | N/A |
| dSPN, reward, no reward, ipsi, contra, max peak | 4f | reward vs. no reward (ipsi/contra), within-mouse | Friedman | 65 | 8 | C-R: 1.34 ± 0.09, I-R 1.43 ± 0.12, C-U: 1.33 ± 0.07, I-U: 1.3 ± 0.13 | Friedman χ² = 1.95, p = 0.58 | N/A |
| dSPN, reward, no reward, ipsi, contra, avg signal | 4f | reward vs. no reward (ipsi/contra), within-mouse | Friedman + WSR post hoc | 65 | 8 | C-R: 0.24 ± 0.1, I-R 0.06 ± 0.06, C-U: -0.4 ± 0.04, I-U: -0.39 ± 0.04 | Friedman χ² = 19.80, p = 0.0002 | C-R vs. I-R: 0.031, C-R vs. C-U: 0.011, C-R vs. I-U: 0.011, I-R vs. C-U: 0.011, I-R vs. I-U: 0.011, C-U vs. I-U: 0.74 |
| dSPN, consecutive rewards, avg signal | 4g | 1, 2, 4, 6, 8 rewards, within-mouse | Friedman + WSR post hoc | 65 | 8 | 1: 0.36 ± 0.1, 2: 0.14 ± 0.07, 4: 0.07 ± 0.07, 6: 0.13 ± 0.07, 8: 0.14 ± 0.06 | Friedman χ² = 21.14, p = 0.0003 | 1.0 vs. 2.0: 0.016, 1.0 vs. 4.0: 0.03, 1.0 vs. 6.0: 0.03, 1.0 vs. 8.0: 0.03 |
| dSPN, consecutive losses, avg signal | 4h | 1, 2, 3, 4, 5, losses, within-mouse | Friedman + WSR post hoc | 65 | 8 | 1: -0.26 ± 0.06, 2: -0.48 ± 0.05, 3: -0.57 ± 0.05, 4: -0.56 ± 0.05, 5: -0.58 ± 0.06 | Friedman χ² = 19.50, p = 0.0006 | 1.0 vs. 2.0: 0.01 1.0 vs. 3.0: 0.01, 1.0 vs. 4.0: 0.01, 1.0 vs. 5.0: 0.01 |
| iSPN, spont lick, max peak | 4i | ipsi vs. contra, within-mouse | WSR | 120 | 18 | contra: 0.83 ± 0.07 vs. ipsi: 0.74 ± 0.07 | Statistic = 47.0, p = 0.10 | N/A |
| iSPN, contra, ipsi, rewarded, unrewarded, max peak | 4k | reward vs. no reward (ipsi/contra), within-mouse | Friedman + WSR post hoc | 120 | 18 | C-R: 0.5 ± 0.12, I-R: 0.69 ± 0.11, C-U: 1.06 ± 0.12, I-U: 1.05 ± 0.09 | Friedman χ² = 30.20, p = 0.000001 | C-R vs. I-R: 0.09, C-R vs. C-U: 0.00007, C-R vs. I-U: 0.00007, I-R vs. C-U: 0.001, I-R vs. I-U: 0.0003, C-U vs. I-U: 0.97 |
| iSPN, contra, ipsi, rewarded, unrewarded, avg signal | 4k | reward vs. no reward (ipsi/contra), within-mouse | Friedman + WSR post hoc | 120 | 18 | C-R: 0.12 ± 0.05, I-R 0.08 ± 0.06, C-U: -0.19 ± 0.05, I-U: -0.19 ± 0.04 | Friedman χ² = 10.80, p = 0.01 | C-R vs. I-R: 0.5, C-R vs. C-U: 0.008, C-R vs. I-U: 0.006, I-R vs. C-U: 0.009, I-R vs. I-U: 0.009, C-U vs. I-U: 0.93 |
| iSPN, consecutive rewards, avg signal | 4m | 1, 2, 4, 6, 8 rewards, within-mouse | Friedman + WSR post hoc | 120 | 18 | 1: 0.25 ± 0.06, 2: 0.06 ± 0.05, 4: 0.05 ± 0.05, 6: 0.1 ± 0.05, 8: 0.11 ± 0.05 | Friedman χ² = 22.13, p = 0.0002 | 1.0 vs. 2.0: 0.0021, 1.0 vs. 4.0: 0.0021, 1.0 vs. 6.0: 0.0084, 1.0 vs. 8.0: 0.0043 |
| iSPN, consecutive losses, avg signal | 4m | 1, 2, 3, 4, 5, losses, within-mouse | Friedman + WSR post hoc | 120 | 18 | 1: -0.05 ± 0.05, 2: -0.26 ± 0.04, 3: -0.35 ± 0.04, 4: -0.35 ± 0.05, 5: -0.34 ± 0.05 | Friedman χ² = 30.44, p = 0.00001 | 1.0 vs. 2.0: 0.00018, 1.0 vs. 3.0: 0.00015, 1.0 vs. 4.0: 0.00018, 1.0 vs. 5.0: 0.00066 |
| dSPN, spont lick, max peak, right hemisphere | Supp 4b | contra vs. ipsi (right hemisphere), within-mouse | WSR | 65 | 8 | contra: 0.92 ± 0.08 vs. ipsi: 0.86 ± 0.07 | Statistic = 16.0, p = 0.84 | N/A |
| dSPN, spont lick, avg signal, right hemisphere | Supp 4b | contra vs. ipsi (right hemisphere), within-mouse | WSR | 65 | 8 | contra: 0.24 ± 0.04 vs. ipsi: 0.22 ± 0.05 | Statistic = 16.0, p = 0.84 | N/A |
| dSPN, spont lick, max peak, left hemisphere | Supp 4b | contra vs. ipsi (left hemisphere), within-mouse | WSR | 65 | 8 | contra: 0.91 ± 0.07 vs. ipsi: 0.92 ± 0.09 | Statistic = 18.0, p = 1.0 | N/A |

| | | | | | | | | |
|---|---|---|---|---|---|---|---|---|
| dSPN, spont lick, avg signal, left hemisphere | Supp 4b | contra vs. ipsi (left hemisphere), within-mouse | WSR | 65 | 8 | contra: 0.24 ± 0.04 vs. ipsi: 0.25 ± 0.06 | Statistic = 16.0, p = 0.84 | N/A |
| dSPN, ipsi spont lick, max peak | Supp 4b | right vs. left hemisphere, within-mouse | MWU | 65 | 8 | ipsi right: 0.86 ± 0.07 vs. ipsi left: 0.92 ± 0.09 | Statistic = 36.0, p = 0.72 | N/A |
| dSPN, contra spont lick, max signal | Supp 4b | right vs. left hemisphere, within-mouse | MWU | 65 | 8 | contra right: 0.92 ± 0.08 vs. contra left: 0.91 ± 0.07 | Statistic = 38.0, p = 0.57 | N/A |
| dSPN, ipsi spont lick, avg signal | Supp 4b | right vs. left hemisphere, within-mouse | MWU | 65 | 8 | ipsi right: 0.22 ± 0.05 vs. ipsi left: 0.25 ± 0.06 | Statistic = 38.0, p = 0.57 | N/A |
| dSPN, contra spont lick, avg signal | Supp 4b | right vs. left hemisphere, within-mouse | MWU | 65 | 8 | contra right: 0.24 ± 0.04 vs. contra left: 0.24 ± 0.04 | Statistic = 33.0, p = 0.96 | N/A |
| dSPN, Cue event response, max peak | Supp 4c | right vs. left hemisphere, within-mouse | MWU | 65 | 8 | cue right: 1.23 ± 0.09, cue left: 1.29 ± 0.09 | Statistic = 28.0, p = 0.72 | N/A |
| dSPN, Choice event response, max peak | Supp 4c | right vs. left hemisphere, within-mouse | MWU | 65 | 8 | choice right: 1.34 ± 0.06, choice left: 1.39 ± 0.1 | Statistic = 25.0, p = 0.51 | N/A |
| dSPN, Outcome event response, max peak | Supp 4c | right vs. left hemisphere, within-mouse | MWU | 65 | 8 | outcome right: 1.35 ± 0.06, outcome left: 1.4 ± 0.1 | Statistic = 27.0, p = 0.65 | N/A |
| dSPN, Cue event response, avg signal | Supp 4c | right vs. left hemisphere, within-mouse | MWU | 65 | 8 | cue right: 0.16 ± 0.14, cue left: 0.14 ± 0.03 | Statistic = 35.0, p = 0.80 | N/A |
| dSPN, Choice event response, avg signal | Supp 4c | right vs. left hemisphere, within-mouse | MWU | 65 | 8 | choice right: 0.06 ± 0.05, choice left: 0.05 ± 0.04 | Statistic = 33.0, p = 0.96 | N/A |
| dSPN, Outcome event response, avg signal | Supp 4c | right vs. left hemisphere, within-mouse | MWU | 65 | 8 | outcome right: -0.004 ± 0.06, outcome left: -0.02 ± 0.0.4 | Statistic = 34.0, p = 0.89 | N/A |
| dSPN, reward, no reward, ipsi, contra, history, max peak | Supp 4d | reward vs. no reward (ipsi/contra), within-mouse | Friedman | 65 | 8 | WR (win) contra:1.32 ± 0.10, WR (win) ipsi: 1.42 ± 0.12, WR (lose) contra: 1.24 ± 0.12, WR (lose) ipsi: 1.31 ± 0.16 | Friedman $\chi^2$ = 0.60, p = 0.90 | N/A |
| dSPN, reward, no reward, ipsi, contra, history, avg signal | Supp 4d | reward vs. no reward (ipsi/contra), within-mouse | Friedman + WSR post hoc | 65 | 8 | WR (win) contra:0.19 ± 0.1, WR (win) ipsi: 0.01 ± 0.06, WR(lose) contra: -0.31 ± 0.07, WR (lose) ipsi: -0.31 ± 0.06 | Friedman $\chi^2$ = 15.45, p = 0.0015 | WR-win-C vs. WR-win-I: 0.041, WR-win-C vs. WR-lose-C: 0.023, WR-win-C vs. WR-lose-I: 0.023, WR-win-I vs. WR-lose-C: 0.023, WR-win-I vs. WR-lose-I: 0.023, WR-lose-C vs. WR-lose-I: 0.84 |
| dSPN, consecutive rewards, max peak | Supp 4e | 1, 2, 4, 6, 8 rewards, within-mouse | Friedman + WSR post hoc | 65 | 8 | 1: 1.46 ± 0.07, 2: 1.38 ± 0.09, 4: 1.31 ± 0.09, 6: 1.42 ± 0.09, 8: 1.65 ± 0.11 | Friedman $\chi^2$ = 14.63, p = 0.0055 | 1.0 vs. 2.0: 0.24, 1.0 vs. 4.0: 0.22, 1.0 vs. 6.0: 0.81, 1.0 vs. 8.0: 0.24 |
| dSPN, consecutive losses, max peak | Supp 4e | 1, 2, 3, 4, 5, losses, within-mouse | Friedman | 65 | 8 | 1: 1.38 ± 0.11, 2: 1.39 ± 0.06, 3: 1.3 ± 0.06, 4: 1.3 ± 0.08, 5: 1.31 ± 0.14 | Friedman $\chi^2$ = 3.60, p = 0.46 | N/A |

| | | | | | | | | |
|---|---|---|---|---|---|---|---|---|
| iSPN, spont lick, max peak, right hemisphere | Supp 4f | contra vs. ipsi (right hemisphere), within-mouse | WSR | 120 | 18 | contra: 0.8 ± 0.11 vs. ipsi: 0.79 ± 0.1 | Statistic=28.0, p =0.42 | N/A |
| iSPN, spont lick, avg signal, right hemisphere | Supp 4f | contra vs. ipsi (right hemisphere), within-mouse | WSR | 120 | 18 | contra: 0.18 ± 0.04 vs. ipsi: 0.15 ± 0.04 | Statistic=28.0, value=0.42 | N/A |
| iSPN, spont lick, max peak, left hemisphere | Supp 4f | contra vs. ipsi (left hemisphere), within-mouse | WSR | 120 | 18 | contra: 0.86 ± 0.09 vs. ipsi: 0.87 ± 0.11 | Statistic=55.0, p =0.80 | N/A |
| iSPN, spont lick, avg signal, left hemisphere | Supp 4f | contra vs. ipsi (left hemisphere), within-mouse | WSR | 120 | 18 | contra: 0.17 ± 0.03 vs. ipsi: 0.17 ± 0.04 | Statistic=55.0, p =0.80 | N/A |
| iSPN, ipsi spont lick, max peak | Supp 4f | right vs. left hemisphere, within-mouse | MWU | 120 | 18 | ipsi right: 0.79 ± 0.1 vs. ipsi left: 0.87 ± 0.11 | Statistic = 101.0, p = 0.61 | N/A |
| iSPN, contra spont lick, max signal | Supp 4f | right vs. left hemisphere, within-mouse | MWU | 120 | 18 | contra right: 0.8 ± 0.11 vs. contra left: 0.86 ± 0.09 | Statistic = 98.0, p = 0.71 | N/A |
| iSPN, ipsi spont lick, avg signal | Supp 4f | right vs. left hemisphere, within-mouse | MWU | 120 | 18 | ipsi right: 0.15 ± 0.04 vs. ipsi left: 0.17 ± 0.04 | Statistic = 101.0, p = 0.61 | N/A |
| iSPN, contra spont lick, averge signal | Supp 4f | right vs. left hemisphere, within-mouse | MWU | 120 | 18 | contra right:0.18 ± 0.04 vs. contra left: 0.17 ± 0.03 | Statistic = 98.0, p = 0.71 | N/A |
| iSPN, Cue event response, max peak | Supp 4g | right vs. left hemisphere, within-mouse | MWU | 120 | 18 | cue right: 0.64 ± 0.08, cue left: 0.77 ± 0.1 | Statistic = 76.0, p = 0.51 | N/A |
| iSPN, Choice event response, max peak | Supp 4g | right vs. left hemisphere, within-mouse | MWU | 120 | 18 | choice right: 0.77 ± 0.13, choice left: 0.81 ± 0.1 | Statistic = 76.0, p = 0.51 | N/A |
| iSPN, Outcome event response, max peak | Supp 4g | right vs. left hemisphere, within-mouse | MWU | 120 | 18 | outcome right: 0.78 ± 0.13, outcome left: 0.8 ± 0.1 | Statistic = 79.0, p = 0.60 | N/A |
| iSPN, Cue event response, mean signal | Supp 4g | right vs. left hemisphere, within-mouse | MWU | 120 | 18 | cue right: 0.09 ± 0.03, cue left: 0.07 ± 0.03 | Statistic = 101.0, p = 0.61 | N/A |
| iSPN, Choice event response, mean signal | Supp 4g | right vs. left hemisphere, within-mouse | MWU | 120 | 18 | choice right: 0.06 ± 0.02, choice left: 0.02 ± 0.03 | Statistic = 105.0, p = 0.48 | N/A |
| iSPN, Outcome event response, mean signal | Supp 4g | right vs. left hemisphere, within-mouse | MWU | 120 | 18 | outcome right: 0.04 ± 0.02, outcome left: -0.01 ± 0.03 | Statistic = 108.0, p = 0.39 | N/A |
| iSPN, reward, no reward, ipsi, contra, history, max peak | Supp 4h | reward vs. no reward (ipsi/contra), within-mouse | Friedman + WSR post hoc | 120 | 18 | WR (win) contra: 0.45 ± 0.12, WR (win) ipsi: 0.65 ± 0.11, WR (lose) contra: 1.05 ± 0.14, WR (lose) ipsi: 1.12 ± 0.11 | Friedman $\chi^2$ = 30.87, p = 0.000001 | WR-win-C vs. WR-win-I: 0.070, WR-win-C vs. WR-lose-C: 0.00005, WR-win-C vs. WR-lose-I: 0.00007, WR-win-I vs. WR-lose-C: 0.013, WR-win-I vs. WR-lose-I: 0.0005, WR-lose-C vs. WR-lose-I: 0.37 |

| | | | | | | | | |
|---|---|---|---|---|---|---|---|---|
| iSPN, reward, no reward, ipsi, contra, history, avg signal | Supp 4h | reward vs. no reward (ipsi/contra), within-mouse | Friedman | 120 | 18 | WR (win) contra: 0.07 ± 0.05, WR (win) ipsi: 0.04 ± 0.06, WR (lose) contra: -0.05 ± 0.07, WR (lose) ipsi: -0.08 ± 0.05 | Friedman χ² = 4.60, p = 0.20 | N/A |
| iSPN, consecutive rewards, max peak | Supp 4i | 1, 2, 4, 6, 8 rewards, within-mouse | Friedman + WSR post hoc | 120 | 18 | 1: 0.74 ± 0.13, 2: 0.54 ± 0.11, 4: 0.5 ± 0.11, 6: 0.63 ± 0.11, 8: 0.73 ± 0.12 | Friedman χ² = 23.56, p = 0.0001 | 1.0 vs. 2.0: 0.34, 1.0 vs. 4.0: 0.34, 1.0 vs. 6.0: 0.34, 1.0 vs. 8.0: 0.70 |
| iSPN, consecutive losses, max peak | Supp 4i | 1, 2, 3, 4, 5, losses, within-mouse | Friedman | 120 | 18 | 1: 1.1 ± 0.11, 2: 1.11 ± 0.11, 3: 1.08 ± 0.12, 4: 0.99 ± 0.12, 5: 1.07 ± 0.13 | Friedman χ² = 3.07, p = 0.55 | N/A |
| dSPN, future swich, avg signal | 5b | LS (win next trial) vs. LR (lose next trial), within-mouse | WSR | 65 | 8 | LS (win next trial): -0.44 ± 0.06 vs. LR (lose next trial): -0.19 ± 0.05 | 0.0078 | N/A |
| iSPN, future switch, avg signal | 5b | LS (win next trial) vs. LR (lose next trial), within-mouse | WSR | 120 | 18 | LS (win next trial): -0.13 ± 0.07 vs. LR (lose next trial): -0.04 ± 0.04 | 0.074 | N/A |
| dSPN, outcome history, avg signal | 5g | WR (win previous trial) vs. LR (lose previous trial), within-mouse | WSR | 65 | 8 | WR (win previous trial): 0.04 ± 0.07 vs. LR (lose previous trial): -0.46 ± 0.07 | 0.0078 | N/A |
| iSPN, outcome history, avg signal | 5g | WR (win previous trial) vs. LR (lose previous trial), within-mouse | WSR | 120 | 18 | WR (win previous trial): -0.01 ± 0.05 vs. LR (lose previous trial): -0.33 ± 0.05 | 3.8147E-05 | N/A |
| dSPN, current outcome, avg signal | Supp 5b | WR (win current trial) vs. WR (lose current trial), within-mouse | WSR | 65 | 8 | WR (win current trial): 0.10 ± 0.07 vs. WR (lose current trial): -0.31 ± 0.06 | 0.0078 | N/A |
| dSPN, current outcome, max peak | Supp 5b | WR (win current trial) vs. WR (lose current trial), within-mouse | WSR | 65 | 8 | WR (win current trial): 1.36 ± 0.09 vs. WR (lose current trial): 1.30 ± 0.11 | 0.38 | N/A |
| iSPN, current outcome, avg signal | Supp 5b | WR (win current trial) vs. WR (lose current trial), within-mouse | WSR | 120 | 18 | WR (win current trial): 0.06 ± 0.05 vs. WR (lose current trial): -0.07 ± 0.06 | 0.25 | N/A |
| iSPN current outcome, max peak | Supp 5b | WR (win current trial) vs. WR (lose current trial), within-mouse | WSR | 120 | 18 | WR (win current trial): 0.77 ± 0.08 vs. WR (lose current trial): 1.1 ± 0.11 | 4.00E-03 | N/A |
| dSPN, future swich, ipsi, contra, avg signal | Supp 5e | LS (win next trial) vs. LR (lose next trial), within-mouse | Friedman + WSR post hoc | 65 | 8 | LS (win) contra: -0.44 ± 0.07, LS (win) ipsi: -0.44 ± 0.06, LR (lose) contra: -0.18 ± 0.05, LR (lose) ipsi: -0.2 ± 0.05 | Friedman χ² = 14.85, p = 0.002 | LS contra vs. LS ipsi: 0.95, LS contra vs. LR contra: 0.023, LS contra vs. LR ipsi: 0.023, LS ipsi vs. LR contra: 0.023, LS ipsi vs. LR ipsi: 0.023, LR contra vs. LR ipsi: 0.95 |
| iSPN, future switch, ipsi, contra, avg signal | Supp 5e | LS (win next trial) vs. LR (lose next trial), within-mouse | Friedman | 120 | 18 | LS (win) contra: -0.11 ± 0.09, LS (win) ipsi: -0.14 ± 0.07, LR (lose) contra: -0.02 ± 0.06, LR (lose) ipsi: -0.06 ± 0.04 | Friedman χ² = 5.0, p = 0.17 | N/A |
| dSPN, outcome history, ipsi, contra, avg signal | Supp 5f | WR (win previous trial) vs. LR (lose previous trial), within-mouse | Friedman + WSR post hoc | 65 | 8 | WR contra: 0.04 ± 0.06, WR ipsi: 0.05 ± 0.09, LR contra: -0.46 ± 0.07, LR ipsi: -0.45 ± 0.07 | Friedman χ² = 19.35, p = 0.0002 | WR contra vs. WR ipsi: 0.55, WR contra vs. WR ipsi: 0.012, WR contra vs. LR ipsi: 0.012, WR ipsi vs. LR contra: 0.012, WR ipsi vs. LR ipsi: 0.012, LR contra vs. LR ipsi: 0.84 |
| iSPN, outcome history, ipsi, contra, avg signal | Supp5f | WR (win previous trial) vs. LR (lose previous trial), within-mouse | Friedman + WSR post hoc | 120 | 18 | WR contra: -0.02 ± 0.06, WR ipsi: -0.001 ± 0.06, LR contra: -0.32 ± 0.05, LR ipsi: -0.33 ± 0.05 | Friedman χ² = 32.07, p = 0.000002 | WR contra vs. WR ipsi: 0.87, WR contra vs. WR ipsi: 0.00008, WR contra vs. LR ipsi: 0.00008, WR ipsi vs. LR contra: 0.0001, WR ipsi vs. LR ipsi: 0.00008, LR contra vs. LR ipsi: 0.73 |

| | | | | | | | | |
|---|---|---|---|---|---|---|---|---|
| dSPN-ChR2, action updating (switching) | 6d | stim vs. no stim (ipsi/contra), within mouse | Friedman + WSR post hoc | 75 | 18 | I→C no stim: 5.6 ± 1.1% vs. stim: 63.7 ± 3.6%; C→I no stim: 8.9 ± 1.37% vs. stim: 3.7 ± 1.1% | Friedman χ² = 49.75, p = 9.10e-11 | C→I: no stim vs. I→C stim, p=2.4e-7; C→I stim vs. I→C stim, p=2.4e-7; I→C: no stim vs. stim, p=2.4e-7; C→I: no stim vs. stim, p=0.009; C→I no stim vs. I→C no stim, p=0.07; C→I stim vs. I→C no stim, p=0.30 |
| iSPN-ChR2, action updating (switching) | 6e | stim vs. no stim (ipsi/contra), within mouse | Friedman + WSR post hoc | 32 | 6 | I→C no stim: 4.6 ± 0.06% vs. stim: 8.1 ± 2.9%; C→I no stim: 6.4 ± 1.4% vs. stim: 48.1 ± 5.2% | Friedman χ² = 18.84, p = 0.0003 | C→I no stim vs. stim, p=0.004; C→I stim vs. I→C no stim, p=0.004; C→I stim vs. I→C stim, p=0.004; C→I no stim vs. I→C no stim, p=0.65; I→C: no stim vs. stim, p=0.75; C→I no stim vs. I→C stim, p=0.92 |
| dSPN-ChR2, number of licks | Supp 6c | stim vs. no stim, within mouse | WSR | 75 | 18 | no stim: 13.6 ± 0.6 vs. stim: 14.7 ± 0.7 licks | 0.17 | N/A |
| dSPN-ChR2, frequency of licks | Supp 6c | stim vs. no stim, within mouse | WSR | 75 | 18 | no stim: 7.7 ± 0.2 vs. stim: 6.2 ± 0.2 Hz | 5.96E-07 | N/A |
| dSPN-ChR2, lick direction | Supp 6c | stim vs. no stim (chosen vs. not chosen direction), within mouse | Friedman + WSR post hoc | 75 | 18 | C-NS: 0.86 ± 0.01, C-S: 0.76 ± 0.02, NC-NS: 0.14 ± 0.02, NC-S: 0.24 ± 0.02 | Friedman χ² = 64.0, p = 8.2e-14 | C-NS vs. NC-NS, p=1.8e-7; C-NS vs. NC-S, p=1.8e-7; C-S vs. NC-NS, p=1.8e-7; C-S vs. NC-S, p=1.8e-7; C-NS vs. C-S, p=2.5e-5; NC-NS vs. NC-S, p=2.5e-5 |
| dSPN-ChR2, reaction time | Supp 6c | stim vs. no stim, within mouse | Friedman + WSR post hoc | 75 | 18 | no stim: 146 ± 9ms, stim switch: 200 ± 42ms, stim no switch: 167 ± 14ms | Friedman χ² = 2.3, p = 0.32 | N/A |
| dSPN-eYFP, action updating (switching) | Supp 6d | stim vs. no stim (ipsi/contra), within mouse | Friedman | 21 | 4 | I→C no stim: 6.30 ± 1.30% vs. stim: 4.75 ± 1.00%; C→I no stim: 5.15 ± 0.88% vs. stim: 7.44 ± 1.94% | Friedman χ² = 4.65, p = 0.20 | N/A |
| dSPN-eYFP, number of licks | Supp 6e | stim vs. no stim, within mouse | WSR | 21 | 4 | no stim: 15.5 ± 1.7 vs. stim: 14.7 ± 1.9 licks | Statistic = 35.0, p = 0.80 | N/A |
| dSPN-eYFP, frequency of licks | Supp 6e | stim vs. no stim, within mouse | WSR | 21 | 4 | no stim: 7.6 ± 0.2 vs. stim: 7.6 ± 0.2 Hz | Statistic = 16.0, p = 0.84 | N/A |
| dSPN-eYFP, lick direction | Supp 6e | stim vs. no stim (chosen vs. not chosen direction), within mouse | Friedman + WSR post hoc | 21 | 4 | C-NS: 0.92 ± 0.03, C-S: 0.92 ± 0.02, NC-NS: 0.08 ± 0.03, NC-S: 0.08 ± 0.03 | Friedman χ² = 19.77, p = 0.00019 | C-NS vs. NC-NS, p=0.01; C-NS vs. NC-S, p=0.01; C-S vs. NC-NS, p=0.01; C-S vs. NC-S, p=0.01; C-NS vs. C-S, p=0.9; NC-NS vs. NC-S, p=0.9 |
| dSPN-eYFP, reaction time | Supp 6e | stim vs. no stim, within mouse | WSR | 21 | 4 | no stim: 146 ± 9ms, stim switch: 200 ± 42ms, stim no switch: 167 ± 14ms | Friedman χ² = 1.75, p = 0.42 | N/A |
| iSPN-ChR2, number of licks | Supp 6g | stim vs. no stim, within mouse | WSR | 32 | 6 | no stim: 11.9 ± 1.3 licks vs. stim: 6.2 ± 0.5 licks | 0.002 | N/A |
| iSPN-ChR2, frequency of licks | Supp 6g | stim vs. no stim, within mouse | WSR | 32 | 6 | no stim: 7.9 ± 0.3 Hz vs. stim: 8.8 ± 0.7 Hz | 0.16 | N/A |
| iSPN-ChR2, lick direction | Supp 6g | stim vs. no stim (chosen vs. not chosen direction), within mouse | Friedman + WSR post hoc | 32 | 6 | C-NS: 0.94 ± 0.02, C-S: 0.90 ± 0.03, NC-NS: 0.06 ± 0.02, NC-S: 0.1 ± 0.03 | Friedman χ² = 26.16, p = 8.83e-06 | C-NS vs. NC-NS, p=0.002; C-NS vs. NC-S, p=0.002; C-S vs. NC-NS, p=0.002; C-S vs. NC-S, p=0.002; NC-NS vs. NC-S, p=0.13; C-NS vs. C-S, p=0.13 |
| iSPN-ChR2, reaction time | Supp 6g | stim vs. no stim, within mouse | WSR | 32 | 6 | no stim: 128 ± 17 ms, stim switch: 160 ± 22 ms, stim no switch: 148 ± 26 ms | Friedman χ² = 2.60, p = 0.27 | N/A |

| | | | | | | | | |
|---|---|---|---|---|---|---|---|---|
| iSPN-eYFP, action updating (switching) | Supp 6h | stim vs. no stim (ipsi/contra), within mouse | Friedman | 22 | 6 | I→C no stim: 6.23 ± 1.21% vs. stim: 10.44 ± 3.51%; C→I no stim: 7.27 ± 1.87% vs. stim: 5.80 ± 3.19% | Friedman χ² = 1.92, p = 0.59 | N/A |
| iSPN-eYFP, number of licks | Supp 6i | stim vs. no stim, within mouse | WSR | 22 | 6 | no stim: 12.06 ± 0.86 licks vs. stim: 12.41 ± 0.96 licks | 0.65 | N/A |
| iSPN-eYFP, frequency of licks | Supp 6i | stim vs. no stim, within mouse | WSR | 22 | 6 | no stim: 7.31 ± 0.22 Hz vs. stim: 7.76 ± 0.08 Hz | 0.055 | N/A |
| iSPN-eYFP, lick direction | Supp 6i | stim vs. no stim (chosen vs. not chosen direction), within mouse | Friedman + WSR post hoc | 22 | 6 | C-NS: 0.93 ± 0.02, C-S: 0.93 ± 0.02, NC-NS: 0.07 ± 0.02, NC-S: 0.07 ± 0.03 | Friedman χ² = 24.29, p = 2.18e-05 | C-NS vs. NC-NS, p=0.005; C-NS vs. NC-S, p=0.005; C-S vs. NC-NS, p=0.005; C-S vs. NC-S, p=0.005; C-NS vs. C-S, p=0.34; NC-NS vs. NC-S, p=0.34 |
| iSPN-eYFP, reaction time | Supp 6i | stim vs. no stim, within mouse | WSR | 22 | 6 | no stim: 160 ± 16 ms, stim switch: 160 ± 34 ms, stim no switch: 178 ± 18 ms | Friedman χ² = 4.33, p = 0.11 | N/A |
| dSPN;GtACR1, action updating (switching) | 6i | inhibti vs. no inhibit (ipsi/contra), within mouse | Friedman + WSR post hoc | 27 | 8 | I→C no stim: 5.4 ± 0.6% vs. stim: 7.0 ± 2.1%; C→I no stim: 5.3 ± 1.1% vs. stim: 38.1 ± 5.0% | Friedman χ² = 19.50, p = 0.00021 | C→I no stim vs. stim, p=0.0015; C→I stim vs. I→C no stim, p=0.0015; C→I stim vs. I→C stim, p=0.0029; C→I no stim vs. I→C no stim, p=0.68; C→I no stim vs. I→C stim, p=0.68; I→C: no stim vs. stim, p=0.68 |
| iSPN;GtACR1, action updating (switching) | 6j | inhibti vs. no inhibit (ipsi/contra), within mouse | Friedman + WSR post hoc | 24 | 11 | I→C no stim: 6.1 ± 0.7% vs. stim: 58.8 ± 5.9%; C→I no stim: 5.6 ± 1.0% vs. stim: 5.5 ± 2.1% | Friedman χ² = 30.15, p = 1.28e-06 | C→I no stim vs. I→C stim, p=6.1e-5; C→I stim vs. I→C stim, p=6.1e-5; I→C: no stim vs. stim, p=6.1e-5; C→I: no stim vs. stim, p=0.83; C→I no stim vs. I→C no stim, p=0.83; C→I stim vs. I→C no stim, p=0.83 |
| dSPN;GtACR1, number of licks | Supp 7h | inhibit vs. no inhibit, within mouse | WSR | 27 | 8 | no stim: 15.2 ± 0.8 licks vs. stim: 9.9 ± 1.1 licks | 5.00E-04 | N/A |
| dSPN;GtACR1, frequency of licks | Supp 7h | inhibit vs. no inhibit, within mouse | WSR | 27 | 8 | no stim: 7.5 ± 0.2 Hz vs. stim: 7.8 ± 0.1 Hz | 0.13 | N/A |
| dSPN;GtACR1, lick direction | Supp 7h | inhibit vs. no inhibit (chosen vs. not chosen direction), within mouse | Friedman + WSR post hoc | 27 | 8 | C-NS: 0.88 ± 0.02 vs. C-S: 0.88 ± 0.05, NC-NS: 0.12 ± 0.02 vs. NC-S: 0.12 ± 0.05 | Friedman χ² = 29.0, p = 2.24e-06 | C-NS vs. NC-NS, p=0.001; C-NS vs. NC-S, p=0.001; C-S vs. NC-NS, p=0.001; C-S vs. NC-S, p=0.001; C-NS vs. C-S, p=0.38; NC-NS vs. NC-S, p=0.38 |
| dSPN;GtACR1, reaction time | Supp 7h | inhibit vs. no inhibit, within mouse | WSR | 27 | 8 | no stim: 149 ± 11 ms, stim switch: 164 ± 28 ms, stim no switch: 155 ± 19 ms | Friedman χ² = 1.17, p = 0.56 | N/A |
| iSPN;GtACR1, number of licks | Supp 7j | inhibit vs. no inhibit, within mouse | WSR | 24 | 11 | no stim: 12.9 ± 1.0 licks vs. stim: 13.1 ± 1.0 licks | 0.78 | N/A |
| iSPN;GtACR1, frequency of licks | Supp 7j | inhibit vs. no inhibit, within mouse | WSR | 24 | 11 | no stim: 7.1 ± 0.2 vs. stim: 5.8 ± 0.2 Hz | 3.00E-04 | N/A |
| iSPN;GtACR1, lick direction | Supp 7j | inhibit vs. no inhibit (chosen vs. not chosen direction), within mouse | Friedman + WSR post hoc | 24 | 11 | C-NS: 0.89 ± 0.02, C-S: 0.75 ± 0.02, NC-NS: 0.11 ± 0.02, NC-S: 0.25 ± 0.02 | Friedman χ² = 43.8, p = 1.67e-09 | C-NS vs. NC-NS, p=4.6e-5; C-NS vs. NC-S, p=4.6e-5; C-S vs. NC-NS, p=4.6e-5; C-S vs. NC-S, p=4.6e-5; C-NS vs. C-S, p=0.001; NC-NS vs. NC-S, p=0.001 |
| iSPN;GtACR1, reaction time | Supp 7j | inhibit vs. no inhibit, within mouse | WSR | 24 | 11 | no stim: 122 ± 12 ms, stim switch: 121 ± 12 ms, stim no switch: 121 ± 12 ms | Friedman χ² = 1.125, p = 0.60 | N/A |

| | | | | | | | | |
|---|---|---|---|---|---|---|---|---|
| GtACR1, action updating (switching) | Supp 7k | inhbiti vs. no inhibit (ipsi/contra), within mouse | Friedman | 18 | 5 | I→C no stim: 6.00 ± 1.87% vs. stim: 6.47 ± 2.96%; C→I no stim: 6.05 ± 1.11% vs. stim: 2.95 ± 1.81% | Friedman $\chi^2$ = 4.071, p = 0.25 | N/A |
| GtACR1, number of licks | Supp 7l | inhibit vs. no inhibit, within mouse | WSR | 18 | 5 | no stim:12.1 ± 1.2 licks vs. stim: 11.2 ± 1.3 licks | 0.48 | N/A |
| GtACR1, frequency of licks | Supp 7l | inhibit vs. no inhibit, within mouse | WSR | 18 | 5 | no stim: 7.6 ± 0.3 vs. stim: 7.6 ± 0.2 Hz | 0.89 | N/A |
| GtACR1, lick direction | Supp 7l | inhibit vs. no inhibit (chosen vs. not chosen direction), within mouse | Friedman + WSR post hoc | 18 | 5 | C-NS: 0.93 ± 0.03, C-S: 0.94 ± 0.03, NC-NS: 0.07 ± 0.03, NC-S: 0.06 ± 0.03 | Friedman $\chi^2$ = 24.55, p = 1.92e-05 | C-NS vs. NC-NS, p=0.005; C-NS vs. NC-S, p=0.005; C-S vs. NC-NS, p=0.005; C-S vs. NC-S, p=0.005; C-NS vs. C-S, p=0.1; NC-NS vs. NC-S, p=0.1 |
| GtACR1, reaction time | Supp 7l | inhibit vs. no inhibit, within mouse | WSR | 18 | 5 | no stim: 114 ± 12 ms, stim switch: 142 ± 31 ms, stim no switch: 104 ± 11 ms | Friedman $\chi^2$ = 3.0, p = 0.22 | N/A |
| dSPN;GtACR1, action updating (switching) | 7d | inhbiti vs. no inhibit (ipsi/contra), within mouse | Friedman + WSR post hoc | 28 | 7 | I→C no stim: 3.64 ± 0.56% vs. stim: 6.11 ± 3.75%; C→I no stim: 5.38 ± 0.72% vs. stim: 67.0 ± 11.11% | Friedman $\chi^2$ = 14.52, p = 0.0002 | C→I no stim vs. stim, p=0.011; C→I stim vs. I→C no stim, p=0.011; C→I stim vs. I→C stim, p=0.011; C→I no stim vs. I→C no stim, p=0.20; C→I no stim vs. I→C stim, p=0.52; I→C: no stim vs. stim, p=0.63 |
| iSPN;GtACR1, action updating (switching) | 7f | inhbiti vs. no inhibit (ipsi/contra), within mouse | Friedman + WSR post hoc | 18 | 6 | I→C no stim: 3.98 ± 1.04% vs. stim: 85.94 ± 5.11%; C→I no stim: 9.87 ± 1.78% vs. stim: 5.10 ± 3.88% | Friedman $\chi^2$ = 21.34, p = 8.96e-5 | C→I no stim vs. I→C stim, p=0.007; C→I stim vs. I→C stim, p=0.007; I→C: no stim vs. stim, p=0.007; C→I no stim vs. I→C no stim, p=0.03; C→I: no stim vs. stim, p=0.2; C→I stim vs. I→C no stim, p=0.6 |
| dSPN;GtACR1, reaction time | Supp 8c | inhibit vs. no inhibit, within mouse | Friedman | 28 | 7 | no stim: 138 ± 11 ms, stim switch: 126 ± 13 ms, stim no switch: 136 ± 12 ms | Friedman $\chi^2$ = 1.40, p = 0.50 | N/A |
| iSPN;GtACR1, reaction time | Supp 8e | inhibit vs. no inhibit, within mouse | Friedman | 18 | 6 | no stim: 116 ± 16 ms, stim switch: 111 ± 37 ms, stim no switch: 85 ± 9 ms | Friedman $\chi^2$ = 2.67, p = 0.26 | N/A |
| GtACR1, action updating (switching) | Supp 8f | inhbiti vs. no inhibit (ipsi/contra), within mouse | Friedman | 18 | 5 | I→C no stim: 4.75 ± 1.33% vs. stim: 2.81 ± 2.27%; C→I no stim: 6.39 ± 1.79% vs. stim: 3.01 ± 1.79% | Friedman $\chi^2$ = 7.50, p = 0.058 | N/A |
| GtACR1, reaction time | Supp 8g | inhibit vs. no inhibit, within mouse | Friedman | 18 | 5 | no stim: 97 ± 12 ms, stim switch: 86 ± 32 ms, stim no switch: 97 ± 12 ms | Friedman $\chi^2$ = 2.00, p = 0.38 | N/A |

**Abbreviations**: WSR (Wilcoxon signed rank test); MWU (Wilcoxon rank sum or Mann-Whitney U Test); Freidman (Non-parametric repeated measures analysis of variance); C-NS=chosen, no stim; NC-NS=not chosen, no stim; C-S=chosen, stim; NC-S=not chosen, stim
